# Lipids in xylem sap of woody plants across the angiosperm phylogeny

**DOI:** 10.1101/763771

**Authors:** H. Jochen Schenk, Joseph M. Michaud, Kerri Mocko, Susana Espino, Tatiana Melendres, Mary R. Roth, Ruth Welti, Lucian Kaack, Steven Jansen

## Abstract

Lipids have been observed attached to lumen-facing surfaces of mature xylem conduits of several plant species, but there has been little research on their functions or effects on water transport, and only one lipidomic study of the xylem apoplast. Therefore, we conducted lipidomic analyses of xylem sap from woody stems of seven plants representing six major angiosperm clades, including basal magnoliids, monocots, and eudicots, to characterize and quantify phospholipids, galactolipids, and sulfolipids in sap using mass spectrometry. Locations of lipids in vessels of *Laurus nobilis* were imaged using TEM and confocal microscopy. Xylem sap contained the galactolipids di- and mono-galactosyldiacylglycerol (DGDG and MGDG), as well as all common plant phospholipids, but only traces of sulfolipids, with total lipid concentrations in extracted sap ranging from 0.18 to 0.63 nmol / mL across all seven species. Contamination of extracted sap from lipids in cut living cells was found to be negligible. Lipid composition of sap was compared to wood in two species and was largely similar, suggesting that sap lipids, including galactolipids, originate from cell content of living vessels. Seasonal changes in lipid composition of sap were observed for one species. Lipid layers coated all lumen-facing vessel surfaces of *Laurus nobilis*, and lipids were highly concentrated in inter-vessel pits. The findings suggest that apoplastic, amphiphilic xylem lipids are a universal feature of angiosperms. The findings require a reinterpretation of the cohesion-tension theory of water transport to account for the effects of apoplastic lipids on dynamic surface tension and hydraulic conductance in xylem.

## Introduction

The lipidome of the plant apoplast has been characterized as a black box (Misra, 2016), because there have been almost no lipidomic analyses of cell walls, intercellular spaces, and xylem conduits and fibers, except for a few studies of surface lipids involved in the making of suberin, cutin, and waxes. Studies of lipids in xylem conduits, including vessels, are especially limited. Wagner *et al*. (2000) observed under transmission electron microscopy (TEM) a dark layer lining vessel and tracheid surfaces of the resurrection plant *Myrothamnus flabellifolia* Welw. (Myrothamnaceae) and after further studies concluded that it was composed of phospholipids (Schneider *et al.*, 2003). Dark layers on vessel walls that appeared under TEM after OsO_4_ fixation were also found in a few other species (Fineran, 1997, Zimmermann *et al.*, 2004, Westhoff *et al.*, 2008). In fact, it had been shown much earlier that a thin coat of lipids remains on the lumen walls of xylem conduits from live cell content after conduit maturation (Scott *et al.*, 1960, Esau, 1965, Esau *et al.*, 1966).

The presence of lipids in xylem conduits may seem difficult to reconcile with the cohesion tension (CT) theory of water transport (Askenasy, 1895, Dixon and Joly, 1895), which posits the existence of negative pressure (i.e., tension) in xylem sap. Lipids are either hydrophobic or amphiphilic, either of which properties would appear to pose a problem for maintaining negative pressure in the sap without forming gas bubbles on hydrophobic or amphiphilic surfaces. Accordingly, Zimmermann *et al*. (2004) cited evidence for lipids in the xylem apoplast to bolster their arguments against the CT theory. However, the CT theory is strongly supported by multiple lines of evidence, both indirect and direct (Angeles *et al.*, 2004, Wheeler and Stroock, 2008, Jansen and Schenk, 2015, Venturas *et al.*, 2017), so the existence of apoplastic xylem lipids would require an explanation within the context of this theory (Schenk *et al.*, 2017).

Any idea that there are no amphiphilic lipids in the xylem apoplast was put to rest when Gonorazky *et al*. (2012) published an analysis of phospholipids in intercellular fluids and xylem fluids of tomato plants. This was followed by reports of phospholipids in xylem sap and on conduit surfaces of five woody angiosperm species from diverse phylogenetic backgrounds (Schenk *et al.*, 2017, Schenk *et al.*, 2018). These findings raise questions about the origins of these lipids, which could be remains from previous living vessel cell content (Scott *et al.*, 1960, Esau, 1965, Esau *et al.*, 1966) or potentially be transported into vessels from conduit-associated parenchyma cells (Morris *et al.*, 2018b). Even more important are questions about their functions in xylem (Schenk *et al.*, 2015, Schenk *et al.*, 2017). However, surely the first question about these lipids regards their chemical nature. What are they? There is clear evidence for phospholipids in the xylem apoplast, but what are their characteristics, and how about galacto- and sulfolipids?

To answer these questions, we conducted a lipidomic analysis of xylem sap extracted from seven species sampled across the angiosperm phylogeny, including five woody angiosperm species included in previous studies (Schenk *et al.*, 2017, Schenk *et al.*, 2018), to quantify and characterize all phospho- and galactolipids in xylem sap and also test for the presence of sulfolipids in a subset of species. Because contamination from cut surfaces is a notorious problem with all xylem sap analyses (Schurr, 1998), we analyzed contamination controls from cut and cleaned xylem surfaces at the sap collection end of stems and compared their lipid conentrations and compositions to xylem sap samples to test if lipids in extracted sap originate from damaged living cells during extraction. We also used a lipid tracer to test for lipid contamination from successive cuts of stems during xylem sap extractions. Because lipids in xylem vessels have been reported to originate from lipid bilayer membranes in the cell content of living vessels (Scott *et al.*, 1960, Esau, 1965, Esau *et al.*, 1966), we compared the lipid composition of xylem sap for two species to that of wood from the same stems to test the prediction that there would be no differences in composition. Lipids in sap sampled for one species were compared in July and March to test for seasonal differences in lipid composition that could indicate developmental effects, lipid transport into sap, or apoplastic enzyme activity. In addition, locations of lipids in xylem were visualized via confocal and transmission electron microscopy in bay laurel, *Laurus nobilis*, a species much studied for its hydraulic properties (e.g., Tyree *et al.*, 1999, Zwieniecki *et al.*, 2001, Hacke and Sperry, 2003, Gascó *et al.*, 2006, Espino and Schenk, 2011, Nardini *et al.*, 2017).

## Material and Methods

### Plant species

The study was conducted with seven plant species from six major angiosperm clades (APG III, 2009) and different growth forms: *Liriodendron tulipifera* L. (winter-deciduous tree, Magnoliaceae, Magnoliales, magnoliid clade), *Laurus nobilis* L., (evergreen tree, Lauraceae, Laurales, magnoliid clade), *Bambusa oldhamii* Munro (bamboo, Poaceae, Poales, commelinid clade), *Triadica sebifera* Small (syn. *Sapium sebiferum*, winter-deciduous tree, Euphorbiaceae, Malpighiales, fabid clade), *Geijera parviflora* Lindl. (evergreen tree, Rutaceae, Sapindales, malvid clade), *Distictis buccinatoria* (DC.) A.H.Gentry (syn. *Amphilophium buccinatorium* (DC.) L.G. Lohmann, evergreen liana, Bignoniaceae, Lamiales, lamiid clade), and *Encelia farinosa* Torr. & A.Gray (drought-deciduous desert shrub, Asteraceae, Asterales, campanulid clade). All grow in the Fullerton Arboretum or on the California State University Fullerton campus in Fullerton, California, United States, except for *Laurus nobilis*, which was collected from the Los Angeles County Arboretum in Arcadia, California, United States. Specimens of *Laurus nobilis* for imaging studies were collected from the Botanical Garden at Ulm University, Germany. All species will be referred to by their generic names in this paper.

Measurements were conducted in two stages: 1. Lipid composition of xylem sap and contamination controls from all seven species: Stems for xylem sap extraction for the first stage of measurements (n = 3 for each species) were collected in August 2017 from *Liriodendron*, *Triadica*, *Geijera*, *Distictis*, and *Encelia*, and in May 2018 from *Bambusa* and *Laurus*. 2. Comparisons of lipid composition in xylem sap and wood in a subset of three species, and additional tests for potential contamination of xylem sap from cut surfaces using lipid tracers in two species (*Geijera* and *Distictis*). Collections for the second stage of measurements (n = 4 for each species) were done in October 2019 for *Liriodendron*, January 2020 for Distictis, and March 2020 for *Geijera*. Some methods were common to both sets of measurements while others differed, as indicated below.

### Xylem sap extraction (all measurements)

Most collections were done at dawn to ensure full hydration of stems. The previously published method for xylem sap extraction (Schenk *et al.*, 2017) was modified to completely eliminate contact of xylem sap with hydrophobic surfaces in order to avoid loss of lipids that could cling to such surfaces (Fig. 1). Samples were only in contact with glass or, during stage 1 measurements, with LoBind surfaces of microcentrifuge tubes (Eppendorf AG, Hamburg Germany), which consist of a two-component polymer mix that creates a hydrophilic surface. Branches were cut from the plants at a length exceeding the longest vessel for each species, which had been previously measured via air injection (Greenidge, 1952). Stems were transported immediately to the lab, and cut under water, with the final cuts made with a fresh razor blade. The bark was removed from the proximal end for about 4 cm length to expose the xylem cylinder (Fig. 1). The cut surface was thoroughly cleaned with deionized water using a high-pressure dental flosser (WP-100 Ultra Water Flosser, Waterpik Inc., Fort Collins, CO, USA) for two minutes to remove cell debris and cytoplasmic content from the surface. A control sample to determine the amount of contamination from living cell remnants from the cleaned, cut surface was taken by inserting the cut end into a glass vial containing 1 mL of nanopure water and leaving it there for one minute. The liquid was then moved in a glass pipette into a pre-weighed 1.5 mL LoBind Eppendorf microcentrifuge tube (for stage 1 extractions using partial freeze drying) or 2 mL glass vials (for stage 2 extractions using a SpeedVac), immediately flash-frozen in liquid nitrogen, and stored in a −20°C freezer until further processing, usually with 1-2 days.

**FIG. 1.**
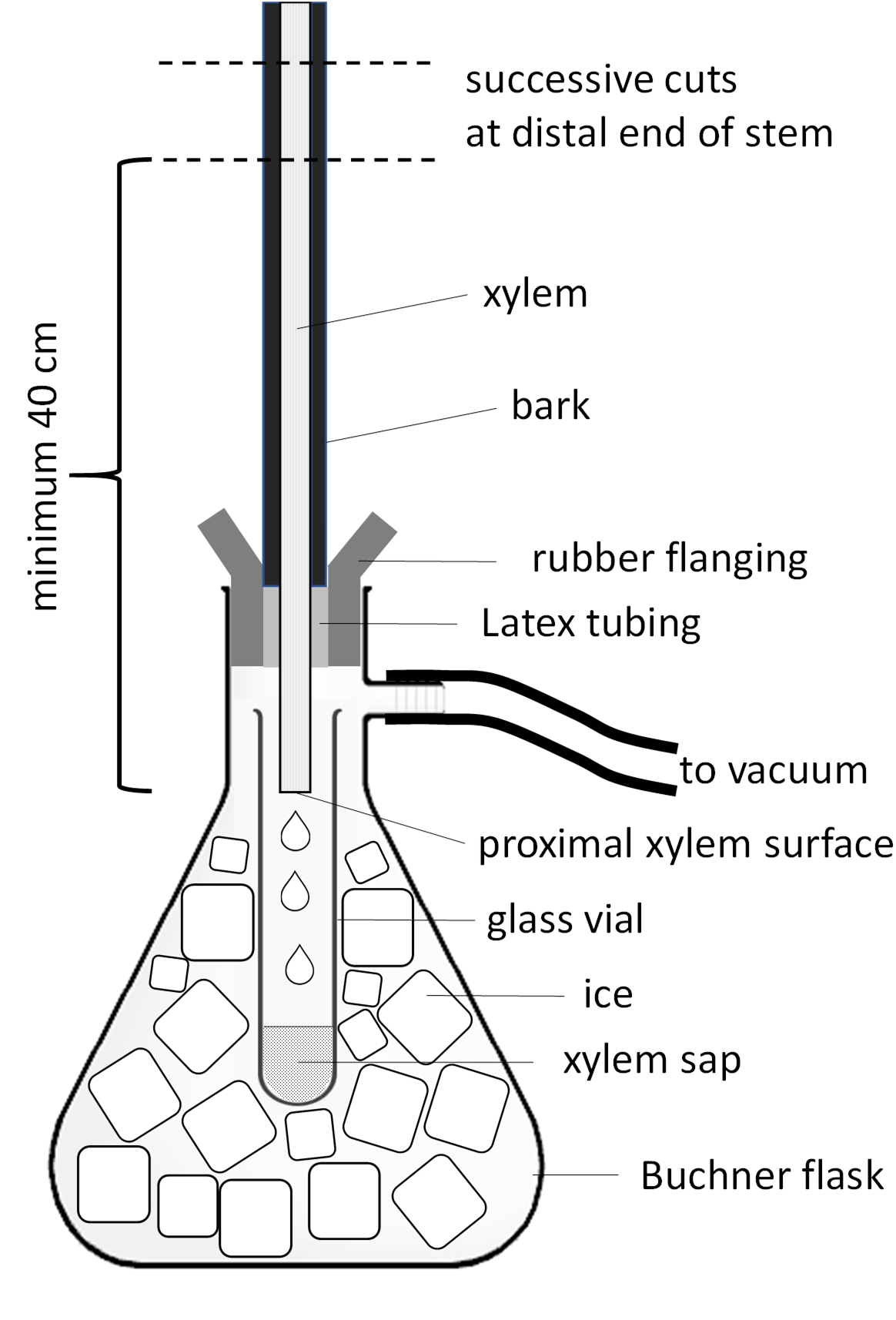
Experimental setup for xylem sap extraction via the vacuum method designed to ensure that lipids in sap only come into contact with glass surfaces and are cooled immediately after extraction to inhibit enzyme activity. Also shown are the proximal xylem surface at the collection end of the stem from which cell contamination controls were collected after high-pressure cleaning of the surface and the successive distal cuts that open vessels for vacuum extraction. In lipid tracer (di17:0 PE) experiments, the tracer was added to the freshly cut distal surfaces of the last three cuts (50, 45, and 40 cm stem length). See Methods section for detailed explanations.

Xylem sap was extracted under vacuum (Fig. 1). Tight-fitting latex tubing, about 2 cm long, was fitted over about half of the exposed xylem cylinder next to the bark, leaving about 2 cm of xylem cylinder freely exposed. The stem was then inserted from the top into a rubber flanged stopper, avoiding any contact of the xylem with the stopper during insertion and creating a tight seal between latex tubing and the stopper. The exposed xylem cylinder was then cleaned again with deionized water using a high-pressure dental flosser and excess water removed with a Kimwipe. Xylem sap was collected in a glass test tube embedded in ice inside a 1 L Buchner flask, which was also embedded in ice. The flask was subjected to lab vacuum for about 30 seconds. The distal end of the branch was then cut back by about 4 cm, followed by successive 2 cm cuts, which were made to all the side branches of the stem until sap was observed dripping into the test tube. Once dripping sap was observed, further 1 cm cuts were made every minute to allow for slow, continuous removal of xylem sap. Depending on stem size, 1 to 2 mL of sap were extracted from each stem, moved with a glass pipette into pre-weighed 1.5 mL LoBind Eppendorf microcentrifuge tubes, (for stage 1 extractions using partial freeze drying) or 2 mL glass vials (for stage 2 extractions using a SpeedVac), flash-frozen in liquid nitrogen, and stored in the freezer until further processing.

Lipids were extracted using two different methods, because a SpeedVac evaporator instrument had become available by the time stage 2 measurements were conducted, which allowed us to simplify procedures. The two methods were compared and validated against each other [**Supplementary Information**].

### Lipid extraction in the stage 1 measurements of seven species: Partial freeze-drying

In preliminary experiments we had found that lipid extraction from completely freeze-dried sap samples was incomplete, most likely due to formation of insoluble aggregates. We therefore developed a new protocol to partially freeze-dry samples for lipid extraction and thereby avoided aggregate formation, which yielded about 8 times more lipids than complete freeze-drying (data not shown). For this protocol, weighed cell contamination controls and xylem sap samples of about 1 mL in pre-weighed LoBind Eppendorf microcentrifuge tunes were partially lyophilized in a freeze-dryer (FreeZone 1 Liter Benchtop Freeze Dry System, Labconco, Kansas City, MO, USA) and observed until only about 100 μL of sample remained. They were then taken from the freeze-dryer and weighed again to determine the remaining volume of aqueous sample. The remaining aqueous samples in two (for *Encelia*, *Bambusa*, and *Laurus* xylem sap and *Bambusa* and *Laurus* controls) or four (all xylem sap and controls from all other species) LoBind microcentrifuge tubes were combined into one sample originating from 2 mL or 4 mL (see breakdown of species and controls above) of xylem sap or control using a glass pipette. Methanol:chloroform 1:1 (both HPLC grader, Fisher Scientific) was added to the combined aqueous sample to create an approximate 5:5:1 methanol:chloroform:water one-phase mixture. The mixture was then vortexed, centrifuged, and the supernatant collected with a 1 mL glass syringe with PrecisionGlide Needle. A new one-phase 5:5:1 mixture of methanol:chloroform:water was then added to the residue, the mixture vortexed, centrifuged, and the supernatant again collected and combined with the previously collected supernatant. Samples were then dried in a desiccator outfitted with an in-line carbon filter capsule (model 6704-7500, Whatman, GE Healthcare Life Sciences, UK) and sent to the Kansas Lipidomics Research Center at Kansas State University for mass spectrometry analysis.

### Lipid extraction in stage 2 measurements of three species: SpeedVac evaporation

The lipid extraction protocol at this stage was changed, because a SpeedVac evaporator designed for chloroform extraction had become available, which allowed us to simplify the procedure. Following xylem sap extraction, samples were immediately freeze-dried using a SpeedVac Concentrator unit (model Savant SPD121P, Thermo Scientific, Waltham, MA, USA). Samples were freeze-dried for 7 hours to permit complete drying of xylem sap in 2 mL glass vials. Freeze-dried samples were then stored at −15 °C.

To isolate lipids from freeze-dried xylem sap samples, a 5:5:1 methanol:chloroform:water one-phase mixture (341 μL methanol, 341 μL chloroform, 68 μL nanopure water) was added to previously fully freeze-dried vials containing xylem sap. Following vortexing for 60 s, the solution was transferred to new 5 mL glass centrifuge tubes via glass pipette and centrifuged at 3500 rpm for 8 min. The supernatant of the centrifuged sample containing lipids was then collected using a glass pipette into a new 2 mL glass vial. This process was repeated to ensure maximal collection of lipids from all glass surfaces, with supernatant of the repeated steps added into the final 2 mL glass vial. Samples were then freeze-dried using the SpeedVac Concentrator unit for 7 h to permit complete removal of all liquid in 2 mL glass vials. Freeze-dried lipid-isolate samples were then stored on dry ice and sent to the Kansas Lipidomics Research Center at Kansas State University for mass spectrometry analysis.

The partial freeze-drying and SpeedVac extraction methods were subjected to a methods comparison and were found to yield similar results (Methods S1) [**Supplementary Information].**

### *Seasonal changes in lipid composition of xylem sap in* Geijera

Xylem sap was extracted from *Geijera* in mid July 2019 (n = 8), and early March 2020 (n = 8). At both times, half of the 8 samples were from lipid tracer experiments (see below), and because no tracer was found in collected xylem sap for *Geijera* (see Results), samples from tracer treatments and controls were combined to increase the statistical power of comparisons. Lipids were extracted as described above, and lipid concentrations and composition compared between sampling times to determine if there were changes between summer and winter collections.

### Lipid extraction from wood in stage 2 measurements of three species

Lipids were extracted from wood by using a modification of methods developed originally for lipid extractions from leaves (Shiva *et al.*, 2018). Following xylem sap extractions (see above) from the same stems, four thin wood sections of exposed xylem (~200 mg) were cut with a sledge microtome (model GSL 1, Schweingruber, Switzerland) and immediately submerged in 0.4 mL of hot isopropanol (75°C) for 15 min in 2 mL glass vials to prevent enzymatic degradation of lipids from cut cells. This procedure made it impossible to determine the precise weight of cut sections, so it allows only determination of lipid composition, not lipid concentrations. To approximate the average fresh weight of wood samples, separate sections from the same stem were cut and weighed. To isolate lipids from wood samples, 0.54 mL chloroform, 0.54 mL methanol, and 0.104 mL nanopure water were added to the vial and softly shaken for 24 h on a compact orbital shaker (Cole-Parmer). The 5/5/1 chloroform/methanol/water solution was then transferred with a glass pipette into a new 2 mL glass vial and freeze-dried using a Speedvac.

### Lipid tracer studies to quantify lipid contamination from cut stem surfaces (stage 2 measurements)

To determine if cutting stems during vacuum extraction of xylem sap introduces lipids into extracted sap, lipid tracer studies were conducted with *Geijera* (short vessels) and *Distictis* (long vessels). The tracer was di17:0 phosphatidylethanolamine (di17:0 PE) (Avanti Polar Lipids, Alabaster, AL, USA), because it was absent in previous measurements of xylem sap. All cutting blades were cleaned vigorously with 1% Alconox solution (product No. 1104, Alconox, White Plains, NY, USA) before each cut to prevent surface contamination with tracer from blades. Sap was extracted from stems as described above until the remaining stems were 50 cm long. At this point, vacuum was relaxed, and 0.1 mL of 0.07 mg/ml di17:0 PE lipid tracer in chloroform was added to the freshly-cut distal stem surface (see Fig. 1). The chloroform evaporated immediately, and the tracer was allowed to absorb into vessels for 15 s. Vacuum was reapplied to the stem until approximately 0.5 mL xylem sap was collected. Two more cuts were done using the same tracer application at 45 cm and 40 cm length, at which point the extraction was concluded. Lipids were extracted from xylem sap as described above for stage 2 measurements.

### Vessel volume analysis

Because lipid micelles are mostly too large to pass through pit membranes (Zhang *et al.*, 2020), we hypothesized that lipid concentrations in extracted xylem sap would be related to the volume of vessels opened when a stem is cut for sap extraction. To determine the average opened vessel volume for each species, we used silicone injections (Sperry *et al.*, 2005, Wheeler *et al.*, 2005). Silicone compound (Rhodorsil RTV-141; Rhodia USA, Cranbury, NJ, USA) mixed with 0.41% blue silicone dye (Blue Silc Pig; Smooth On, Easton, PA, USA) was injected from the proximal end into flushed stems (*n* = 4 per species) that were between 0.9 and 1.7 cm in diameter and 25 cm in length, except 50 cm in length for *Distictis*. Injection took place at approximately 50 kPa for 24 h, after which the silicone was allowed to dry for 72 h prior to sectioning. Cross sections (cut at 0.7, 1.4, 2.9, 5.8, 8.3, 11.8, and 24 cm from the cut surface for all species, except at 0.7, 1.4, 2.8, 5.8, 11.6, 16.6, 23.6, and 48 cm for *Distictis*) were thin-sectioned using a sledge microtome (model GSL 1, Schweingruber, Switzerland), dry-mounted on glass slides, and imaged using a Leica MZ16 stereomicroscope with digital camera (model INFINITY 2-1C-IQ, Lumenera, Ottawa, ONT, Canada) at 20× zoom and backlit using a transmitted light stage connected to a halogen light source (model KL 1500 LCD, Schott, Germany). Adobe Photoshop (version CS6) was used to blend images into one combined image for each cross-section (Fig. S10) [**Supplementary Information**], and an analysis of filled vessels, vessels diameters, vessel wall perimeters, and vessel density was completed using ImageJ (Schneider *et al.*, 2012). Silicone-filled vessels were counted in each image, and the area of silicone-filled vessels in each image was quantified both in mm^2^ and as percent of the total xylem area. The silicone-filled vessel volume was calculated from these data by linear interpolation between each cross-section (Fig. S11) [**Supplementary Information**]. Counts of filled vessels (*C*) at each distance (*d*) from the injected surface were used to estimate median vessel lengths, based on fitting the equation ln *C* = a + b × *d* using the software TableCurve 2D (version 5.01, Systat, San Jose, CA, USA).

### Mass spectrometry

Direct-infusion electrospray ionization triple-quadrupole mass spectrometry was performed using a Waters Xevo TQS mass spectrometer or Applied Biosystems 4000 (Shiva *et al.*, 2013). Instrument-specific details can be found in Supplemental Tables S1 and S2. The dried samples, each originating from 2 mL or 4 mL of xylem sap or cell contamination control, or from wood samples (see breakdown of species above), were dissolved in 1 mL chloroform in a 2 mL glass vial. After a test to determine the optimal amount for analysis, 50-1000 μL of each sample was transferred to a 2-mL vial containing internal standards (Supplemental Table S2). 1.2 mL of chloroform:methanol:300 mM ammonium acetate in water (30:66.5:3.5) was added and the sample mixed. The sample was continuously infused to the mass spectrometer from a 1-mL loop.

In addition to the scans described in Shiva *et al*. (2013), a scan for SQDG (Pre 261.1) in positive ion mode was also acquired (except for samples from *Laurus* and *Bambusa*) (details in Supplemental Table S2). Data were processed as described in Shiva *et al*. (2013) using the LipidomeDB Data Calculation Environment at http://lipidome.bcf.ku.edu:8080/Lipidomics/ (Zhou *et al.*, 2011). Appropriate response factor for the biological galactolipid molecular species in comparison to the saturated galactolipid internal standards were applied. The limit of detection in the samples analyzed was 0.002 nmol for the stage 1 measurements of seven species. And 0.0005 nmol for stage 2 measurements.

### Confocal laser scanning microscopy

To test for lipids in xylem of *Laurus*, fluorescent FM1-43 dye (Molecular Probes, Life Technologies, Eugene, OR, USA) was infiltrated into living xylem and imaged with laser scanning confocal microscopy, as described in Schenk *et al*. (2018). FM1-43 is an amphiphilic fluorophore that is virtually non-fluorescent in water and strongly fluorescent under cyan light excitation when bound to lipids, including amphiphilic lipids, and biological membranes (Jelínková *et al.*, 2010). It was used as a 5 μg/mL solution in nanopure water with 488 nm excitation and an emission window of 578 to 618 nm. Woody stems, approximately 5-8 mm in diameter and about 10 cm long, were cut from the plants and submerged in DI water. About 2.5 cm were cut under water from each end of the stem to remove most bubbles in the xylem that may have been induced by the cutting. The cut ends were then recut with new razor blades to create clean and smooth surfaces. Bark was removed from about 1 cm length at the distal end to expose the xylem. The proximal end of each stem was connected to a vacuum flask using tight-fitting latex tubing around the stem inside Tygon tubing. A short piece of Tygon tubing was fitted to the exposed xylem cylinder at the distal end, using latex tubing around the cylinder to create a tight fit. The distal cut end was cleaned by washing it three times with 500 μL nanopure water using a pipette. 1 mL of fluorophore solution was then added into the tubing and onto the exposed xylem surface. Lab vacuum was turned on and turned off once the solution was absorbed into the xylem, but no air was aspirated. The control was treated the same way but, instead of the fluorophore solution, 1 mL of nanopure water was sucked through the stem. A 2 mm long stem piece was then cut along its transverse plane from the center of the stem segment and the distal surface recut using a sledge microtome (Sledge microtome GSL 1, Schweingruber, Switzerland). The stem piece was then placed onto a cell culture dish with a glass bottom of 175 μm (CELLview™ dish, Greiner Bio-One, Germany) in a drop of water and observed using the inverted microscope of a confocal laser scanning microscope (model TCS SP8 with a HyVolution super-resolution detector, Leica Microsystems, Wetzlar, Germany). Lignin was detected with 405 nm excitation and an emission window of 467 to 509 nm. Imaging was done with a 63.0 × 1.20 objective and 3× digital zoom in water, using four scans per image at 1760 × 1760 resolution.

### Transmission electron microscopy (TEM)

Small sectioning blocks of xylem from fresh *Laurus* stems were given two different treatments as described in Schenk *et al*. (2017): (1) fixation with a solution containing 2.5% glutaraldehyde, 1% sucrose, and 0.1 M phosphate buffer at pH 7.3, and (2) fixation with glutaraldehyde and then postfixation with a 2% aqueous osmium tetroxide (OsO_4_) solution for 2 hours at room temperature. The samples were dehydrated through a gradual ethanol series and embedded in Epon resin. Moreover, samples treated with glutaraldehyde only were observed with and without post staining of TEM grids with aqueous uranyl acetate and lead citrate to test the potential staining effect on vessel-vessel pit membranes (Ellis, 2014). The duration of the staining was 5 minutes for uranyl acetate and 1 minute for lead citrate. Transverse semi-thin sections were cut with an ultramicrotome (Leica Ultracut UCT, Leica Microsystems, Vienna, Austria), stained with 0.5% toluidine blue in 0.1 M phosphate buffer, and mounted on microscope slides using Eukitt. Ultra-thin sections between 60 nm and 90 nm were mounted on copper grids (Athena, Plano GmbH, Wetzlar, Germany) and observed with a JEM-1210 TEM (Jeol, Tokyo, Japan) at 120 kV. Digital images were taken using a MegaView III camera (Soft Imaging System, Münster, Germany).

### Data analysis

To test for the presence of di17:0 PE tracer in tracer experiments, concentrations of di17:0 PE in xylem sap from stems treated with di17:0 PE tracer at cut stem surfaces were compared to controls without tracer applications using two-tailed t-tests (n = 4) assuming equal variance in Microsoft Excel. Mean lipid concentrations were compared to various anatomical parameters of xylem for each species (incl. opened vessel volume, opened vessel volume relative to surface area, mean vessel diameter and wall perimeter, both calculated from the diameter of a circle with the same area as the vessel, and mean vessel density) via reduced major axis regression in the statistical software package PAST (ver. 4.02) (Hammer *et al.*, 2001).

To compare lipid compositions between samples, mass spectrometry data were analyzed using the online MetaboAnalyst software package (https://www.metaboanalyst.ca). Any concentrations below the detection limit were set to zero, and data were log-transformed to normalize distributions. Compositions of polar lipids in xylem sap were compared in MetaboAnalyst as normalized concentrations between seven species and between two sampling periods for *Geijera* using principal component analysis (PCA) and heatmaps, based on Euclidean distance measures, and a Ward clustering algorithm to visualize differences among the species or sampling times. Unpaired t-tests, adjusted for false discovery rates (FDR) were used to identify lipids that differed between sampling times.

Compositions of polar lipids in xylem sap from six species (one sample for *Encelia* was lost, causing the species to be excluded from this analysis) were compared to those in paired contamination controls to test the hypothesis that xylem sap lipids originate from damaged living cells and therefore do not differ in lipid composition to contamination controls. For these comparisons, concentrations were converted into percentages, data were log-transformed and analyzed using PCA in MetaboAnalyst.

Polar lipid compositions were compared between wood and xylem sap for *Geijera* and *Distictis* by converting concentrations to percentages, then excluding any measurements that were below the detection limit, and using PCA (95% CI). Paired t-tests adjusted for FDR were used to identify lipids that differed between sap and wood. Because wood samples included xylem sap lipids, the two data sets were not independent, so PCA and t-tests were used qualitatively to determine if there were differences between sap and wood and identified lipids that differed the most.

## Results

### Comparisons of xylem sap and contamination controls (seven species)

Polar lipid concentrations in cell contamination controls ranged from 1.1% of the amount detected in xylem sap in *Distictis* to 9.0% in *Laurus*, averaging 4.49 (± 2.80 SD) percent across species (Table 1). The complete lipidomics data set may be found in Table S2 for xylem sap and Table S3 for cell contamination controls [**Supplementary Information**]. All contamination controls were far more variable and distinctly different in polar lipid composition compared to xylem sap, as shown in principal component analyses (PCA; Fig. S12), clearly demonstrating that xylem sap lipids do not originate from cells cut open during sap extraction [**Supplementary Information**].

**Table 1.** Composition of xylem sap lipids (in nmol / mL) for the seven studied species determined by mass spectrometry, arranged by head groups, and expressed as mean ± standard deviation (n = 3). LysoPG was excluded from these measurements, because trace amounts of a contaminant that was also found in solvent interfered with the detection of LPG in stage 1 measurements.

|  | *Laurus* |  | *Liriodendron* |  | *Bambusa* |  | *Triadica* |  | *Geijera* |  | *Distictis* |  | *Encelia* |  |
| --- | --- | --- | --- | --- | --- | --- | --- | --- | --- | --- | --- | --- | --- | --- |
|  | Mean | STD | Mean | STD | Mean | STD | mean | STD | mean | STD | mean | STD | Mean | STD |
| Xylem sap |  |  |  |  |  |  |  |  |  |  |  |  |  |  |
| DGDG | 0.038 ± 0.013 | | 0.015 ± 0.010 | | 0.203 ± 0.046 | | 0.051 ± 0.009 | | 0.015 ± 0.006 | | 0.041 ± 0.01 | | 0.015 ± 0.008 | |
| MGDG | 0.099 ± 0.052 | | 0.050 ± 0.041 | | 0.158 ± 0.022 | | 0.042 ± 0.007 | | 0.032 ± 0.012 | | 0.050 ± 0.016 | | 0.021 ± 0.010 | |
| PG | 0.001 ± 0.000 | | 0.002 ± 0.002 | | 0.003 ± 0.001 | | 0.007 ± 0.002 | | 0.003 ± 0.003 | | 0.003 ± 0.001 | | 0.004 ± 0.004 | |
| LysoPC | 0.000 ± 0.000 | | 0.003 ± 0.003 | | 0.006 ± 0.003 | | 0.004 ± 0.002 | | 0.001 ± 0.001 | | 0.001 ± 0.001 | | 0.007 ± 0.004 | |
| LysoPE | 0.002 ± 0.002 | | 0.001 ± 0.001 | | 0.003 ± 0.002 | | 0.002 ± 0.002 | | 0.001 ± 0.001 | | 0.001 ± 0.001 | | 0.005 ± 0.005 | |
| PC | 0.022 ± 0.003 | | 0.040 ± 0.017 | | 0.088 ± 0.046 | | 0.148 ± 0.042 | | 0.070 ± 0.031 | | 0.087 ± 0.032 | | 0.071 ± 0.053 | |
| PE | 0.007 ± 0.001 | | 0.016 ± 0.014 | | 0.026 ± 0.020 | | 0.042 ± 0.042 | | 0.051 ± 0.016 | | 0.046 ± 0.018 | | 0.046 ± 0.055 | |
| PI | 0.007 ± 0.003 | | 0.009 ± 0.009 | | 0.027 ± 0.009 | | 0.068 ± 0.018 | | 0.035 ± 0.027 | | 0.051 ± 0.030 | | 0.071 ± 0.081 | |
| PS | 0.001 ± 0.000 | | 0.004 ± 0.003 | | 0.007 ± 0.003 | | 0.011 ± 0.006 | | 0.016 ± 0.007 | | 0.013 ± 0.008 | | 0.015 ± 0.015 | |
| PA | 0.004 ± 0.002 | | 0.041 ± 0.03 | | 0.112 ± 0.049 | | 0.120 ± 0.047 | | 0.223 ± 0.125 | | 0.153 ± 0.090 | | 0.262 ± 0.144 | |
| Total | 0.181 ± 0.071 | | 0.179 ± 0.124 | | 0.633 ± 0.166 | | 0.494 ± 0.177 | | 0.447 ± 0.213 | | 0.447 ± 0.203 | | 0.517 ± 0.371 | |
| SQDG | n.d. | | 0.001 ± 0.001 | | n.d. | | 0.003 ± 0.000 | | 0.003 ± 0.001 | | 0.003 ± 0.001 | | 0.009 ± 0.003 | |
| Contamination controls |  |  |  |  |  |  |  |  |  |  |  |  |  |  |
| DGDG | 0.003 ± 0.001 | | 0.005 ± 0.007 | | 0.005 ± 0.004 | | 0.001 ± 0.001 | | 0.001 ± 0.002 | | 0.002 ± 0.001 | | 0.000 ± 0.000 | |
| MGDG | 0.004 ± 0.002 | | 0.004 ± 0.004 | | 0.010 ± 0.009 | | 0.001 ± 0.000 | | 0.002 ± 0.001 | | 0.002 ± 0.001 | | 0.001 ± 0.000 | |
| PG | 0.000 ± 0.000 | | 0.000 ± 0.000 | | 0.000 ± 0.001 | | 0.000 ± 0.000 | | 0.000 ± 0.000 | | 0.000 ± 0.000 | | 0.000 ± 0.000 | |
| LysoPC | 0.000 ± 0.000 | | 0.000 ± 0.000 | | 0.001 ± 0.000 | | 0.000 ± 0.000 | | 0.000 ± 0.000 | | 0.000 ± 0.000 | | 0.000 ± 0.000 | |
| LysoPE | 0.003 ± 0.002 | | 0.000 ± 0.000 | | 0.001 ± 0.001 | | 0.000 ± 0.000 | | 0.000 ± 0.000 | | 0.000 ± 0.000 | | 0.000 ± 0.000 | |
| PC | 0.002 ± 0.002 | | 0.003 ± 0.001 | | 0.008 ± 0.013 | | 0.008 ± 0.005 | | 0.007 ± 0.001 | | 0.003 ± 0.000 | | 0.012 ± 0.004 | |
| PE | 0.000 ± 0.000 | | 0.000 ± 0.000 | | 0.000 ± 0.000 | | 0.002 ± 0.002 | | 0.000 ± 0.000 | | 0.000 ± 0.000 | | 0.000 ± 0.000 | |
| PI | 0.001 ± 0.001 | | 0.000 ± 0.000 | | 0.001 ± 0.001 | | 0.002 ± 0.003 | | 0.000 ± 0.000 | | 0.000 ± 0.000 | | 0.000 ± 0.000 | |
| PS | 0.001 ± 0.001 | | 0.000 ± 0.000 | | 0.000 ± 0.000 | | 0.000 ± 0.000 | | 0.000 ± 0.000 | | 0.000 ± 0.001 | | 0.000 ± 0.000 | |
| PA | 0.000 ± 0.000 | | 0.007 ± 0.004 | | 0.002 ± 0.002 | | 0.011 ± 0.006 | | 0.009 ± 0.006 | | 0.002 ± 0.000 | | 0.006 ± 0.001 | |
| Total | 0.014 ± 0.008 | | 0.020 ± 0.015 | | 0.029 ± 0.029 | | 0.025 ± 0.016 | | 0.020 ± 0.008 | | 0.009 ± 0.003 | | 0.021 ± 0.003 | |
| SQDG | n.d. | | 0.001 ± 0.001 | | n.d. | | 0.003 ± 0.000 | | 0.003 ± 0.001 | | 0.003 ± 0.001 | | 0.009 ± 0.003 | |

Galactolipids: DGDG = digalactosyldiacylglycerol, MGDG = and monogalactosyldiacylglycerol. Phospholipids: LysoPG = lysophosphatidylglycerol, LysoPC = lysophosphatidylcholine, LysoPE = lysophosphatidylethanolamine, PA = phosphatidic acid, PC = phosphatidylcholine, PE = phosphatidylethanolamine, PI = phosphatidylinositol, PS = phosphatidylserine, PG = phosphatidylglycerol. SQDG = Sulfolipids.

### Lipid tracer studies to detect lipid contamination from cut surfaces (two species)

Concentrations of di17:0 PE in xylem sap of *Geijera* stems treated with di17:0 PE tracer at cut distal stem surfaces and in controls without tracer applications were below 1 pmol/mL in both treatments and not significantly different (*p* = 0.738). Concentrations of di17:0 PE in xylem sap of *Distictis* stems treated with di17:0 PE tracer at cut distal stem surfaces were 0.9 pmol/mL (±0.8 SE), 0.2 pmol/mL (± 0.1 SE) in controls without tracer applications, and not significantly different (*p* = 0.131).

### Xylem sap lipidomics (seven species)

Polar lipids extracted from xylem sap and cell contamination controls included the galactolipids digalactosyldiacylglycerol (DGDG, Fig. S1) and monogalactosyldiacylglycerol (MGDG, Fig. S2), as well as all common plant phospholipids. Total lipid concentrations for xylem sap varied between 0.18 nmol / mL for *Laurus* and *Liriodendron* and 0.63 nmol / mL for *Bambusa*, averaging 0.41 (± 0.17 SD) nmol / mL across species (Table 1; Fig. 2 A). Lipid concentrations in sap were not statistically related to the vessel volume opened by cutting stems, but there was a significant negative relationship to the total vessel perimeter at the xylem surface (Fig. 2 C).

**FIG. 2.**
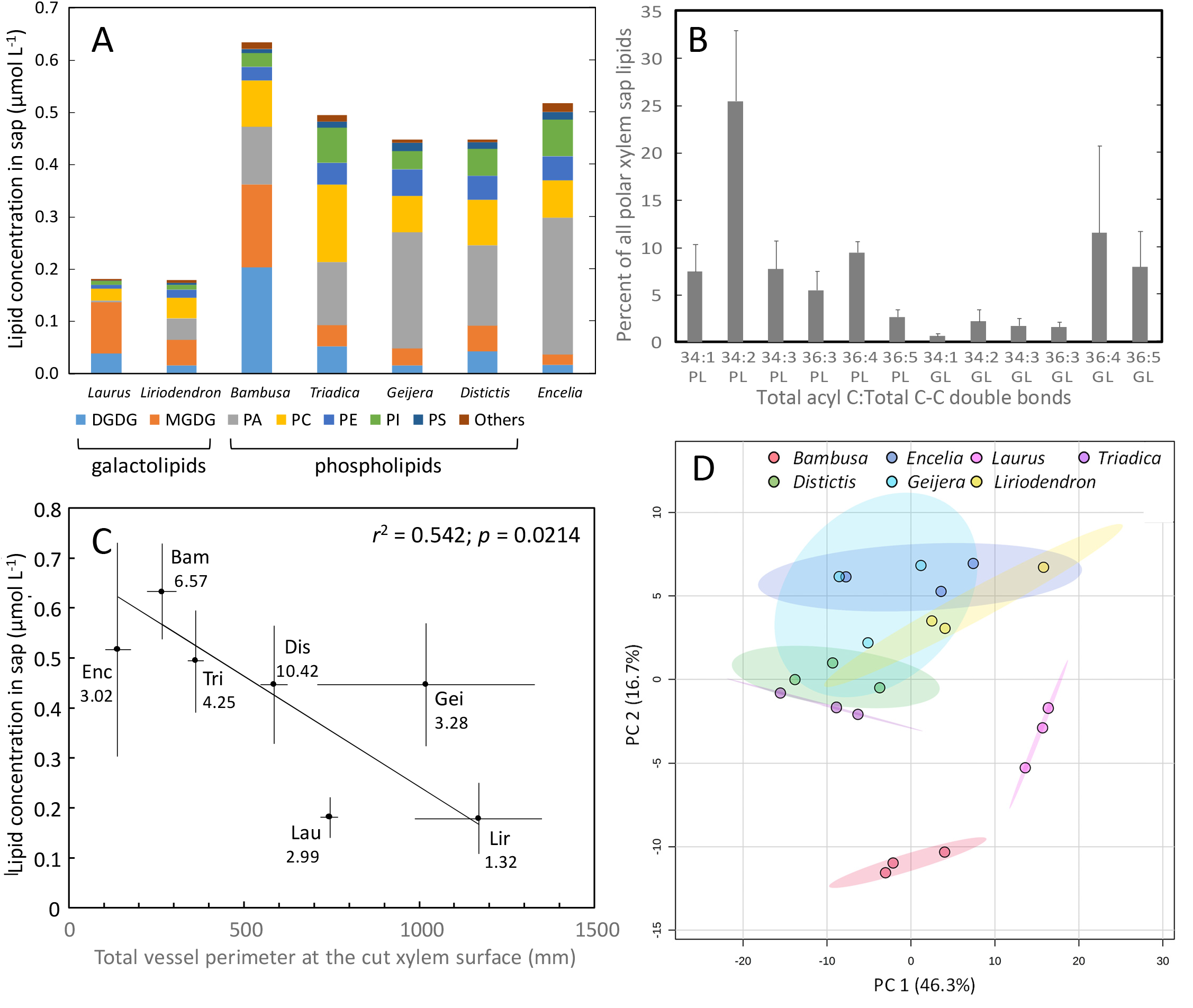
Polar lipid composition of xylem sap from seven angiosperm species (n = 3). A. Chemical composition of xylem sap lipids as determined by direct-infusion electrospray ionization triple-quadrupole mass spectrometry. See Table S1 for the standards used and Figures S1-9 and Tables S2-3 [**Supplementary Information**] for the complete data sets, including chains lengths and degrees of saturation. Sulfolipids were only found in trace amounts (Tables S2 and S3) and are not included in this figure. B. Most common numbers of acyl chain carbon atoms (mean ± SE) in xylem sap lipids (PL = phospholipids, GL = galactolipids) and their degree of unsaturation as determined by direct-infusion electrospray ionization triple-quadrupole mass spectrometry. See Table S2 [**Supplementary Information**] and Figures S1-9 for the complete data set. C. Total concentration of phospho- and galactolipids in xylem sap (mean ± SE) as a function of total vessel xylem perimeter at the cut xylem surface (mean ± SE). Numbers under the abbreviated generic names are the median vessel lengths in cm. Abbreviations: Bam = *Bambusa*, Dis = *Distictis*, Enc = *Encelia*, Gei = *Geijera*, Lau = *Laurus*, Lir = *Liriodendron*, Tri = *Triadica*. D. Polar lipid composition of xylem sap compared via principal component analysis (PCA), with the first two components shown that explain 63% of the variation among species. Shaded areas are 95% confidence regions.

The most common lipids grouped by their head groups (Fig. 2 A) were DGDG, MGDG, phosphatidylcholine (PC, Fig. S4), phosphatidic acid (PA, Fig. S8), phosphatidylethanolamine (PE, Fig. S5), phosphatidylinositol (PI, Fig. S6), with other phospholipids present in much lower amounts, including phosphatidylserine (PS, Fig. S7), phosphatidylglycerol (PG, Fig. S3), lysophosphatidylethanolamine (LPE, Fig. S9), lysophosphatidylcholine (LPC, Fig. S9), and lysophosphatidylglycerol (LPG). Trace amounts of a contaminant that was also found in solvent interfered with the detection of LPG, which was therefore excluded from all tables, figures, and analyses for stage 1 measurements. (The problem was solved for stage 2 measurements by using glass vials instead of LoBind Eppendorf tubes in all subsequent experiments.)

The most pronounced differences in chemical compositions between species were in the percentage of galactolipids (*Laurus* 75.4%, *Liriodendron* 35.9%, *Bambusa* 57.0%, *Triadica* 18.8%, *Geijera* 10.6%, *Distictis* 20.4%, *Encelia* 7.1%). Sulfolipids were detected only in trace amounts. In the PCA comparing the seven species, no species stood out as different in polar lipid compositions from the other six species on the PC1 axis, which explained 46.3% of the variation, while *Laurus* and *Bambusa* stood out on the PC2 axis, explaining 16.7% of the variation, as distinctly different (Fig. 2 D). The most pronounced differences between species were in the percentages contributed by DGDG, MGDG, PC, and PA (Table 1; Fig. 2 A). DGDG varied from about 3.0% of all lipids in *Encelia* to 32.0% in *Bambusa*, with a mean across species of 12.4 (±10.5 SD)%; MGDG varied from 4.1% in *Encelia* to 54.6% in *Laurus*, with a mean across species of 19.8 (±17.8 SD)%; PC varied from 12.3% in *Laurus* to 30.1% in *Triadica*, with a mean across species of 18.2 (±6.3 SD)%; PA varied from 1.9% in *Laurus* to 50.6% in *Encelia*, with a mean across species of 28.7 (±17.6 SD)%.

The most common number of acyl chain carbon atoms were 34 and 36, with galactolipids mostly having more 36-carbon combinations than phospholipids (Fig. 2 B, Fig. S1-2). Only on average 1.56 (± 0.70 SD)% of all lipids were fully saturated phospholipids, and the most common numbers of double bonds were 1-3 for the 34 carbon combination of 2 acyl chains and 3-5 for the 36 carbon combination (Fig. 2 B, Fig. S3-9). Averaged across species, lipids with 34:1-3 (carbon atoms:number of double bonds) and 36:3-5 accounted for 84.0 (± 4.9 SD)% of all apoplastic xylem lipids. PA 34:2 was especially abundant in all species, except for *Laurus*, making up about 23% of all polar lipids in *Encelia* and 25% in *Geijera*. The lipid composition of the galactolipids, with a total of four double bonds prominent, suggests that 18:2 fatty acids may be common.

### *Seasonal changes in lipid composition of xylem sap in* Geijera

Total concentrations of lipids in xylem sap were not different between samples collected in July and March, but concentrations of the galactolipids DGDG and MGDG were significantly higher in sap collected in March (Table 2). Concentrations of PA tended to be lower in March, but differences were not significant because of high variability among samples. Overall, 18 individual lipids differed significantly in their concentrations between July and March (Fig. 3 B), including seven types of DGDG, four of MGDG, and the remaining ones phospholipids. In PCA (Fig. 3 A), March samples were slightly more variable than July samples, and the 95% confidence regions for the two periods were clearly separated on the PC1 axis, which explained 27% of the variation. Overall, there were clear seasonal differences in lipid composition.

**Table 2.** Comparison of lipid composition categorized by headgroups in polar lipids in xylem sap of *Geijera* (n = 8) sampled in July 2019 and March 2020. Mean concentrations (nmol / mL) were compared between dates using FDR-adjusted t-tests in Metaboanalyst.ca.

| Lipid type | July 2019 | March 2020 | p-value |
| --- | --- | --- | --- |
|  | Mean ± STD | Mean ± STD | (FDR adjusted) |
| DGDG | 0.0143 ± 0.0046 | 0.0589 ± 0.0284 | 0.00027 |
| MGDG | 0.0230 ± 0.0096 | 0.0472 ± 0.0193 | 0.03036 |
| PG | 0.0137 ± 0.0059 | 0.0174 ± 0.0097 | n.s. |
| LysoPG | 0.0074 ± 0.0025 | 0.0072 ± 0.0045 | n.s. |
| LysoPC | 0.0033 ± 0.0024 | 0.0018 ± 0.0004 | n.s. |
| LysoPE | 0.0018 ± 0.0011 | 0.0009 ± 0.0002 | n.s. |
| PC | 0.0851 ± 0.0407 | 0.0860 ± 0.0348 | n.s. |
| PE | 0.0550 ± 0.0368 | 0.0512 ± 0.0181 | n.s. |
| PI | 0.0185 ± 0.0114 | 0.0224 ± 0.0119 | n.s. |
| PS | 0.0148 ± 0.0068 | 0.0114 ± 0.0031 | n.s. |
| PA | 0.5329 ± 0.3504 | 0.3050 ± 0.1206 | n.s. |
| Total Polar | 0.7698 ± 0.4546 | 0.6094 ± 0.2025 | n.s. |

**FIG. 3.**
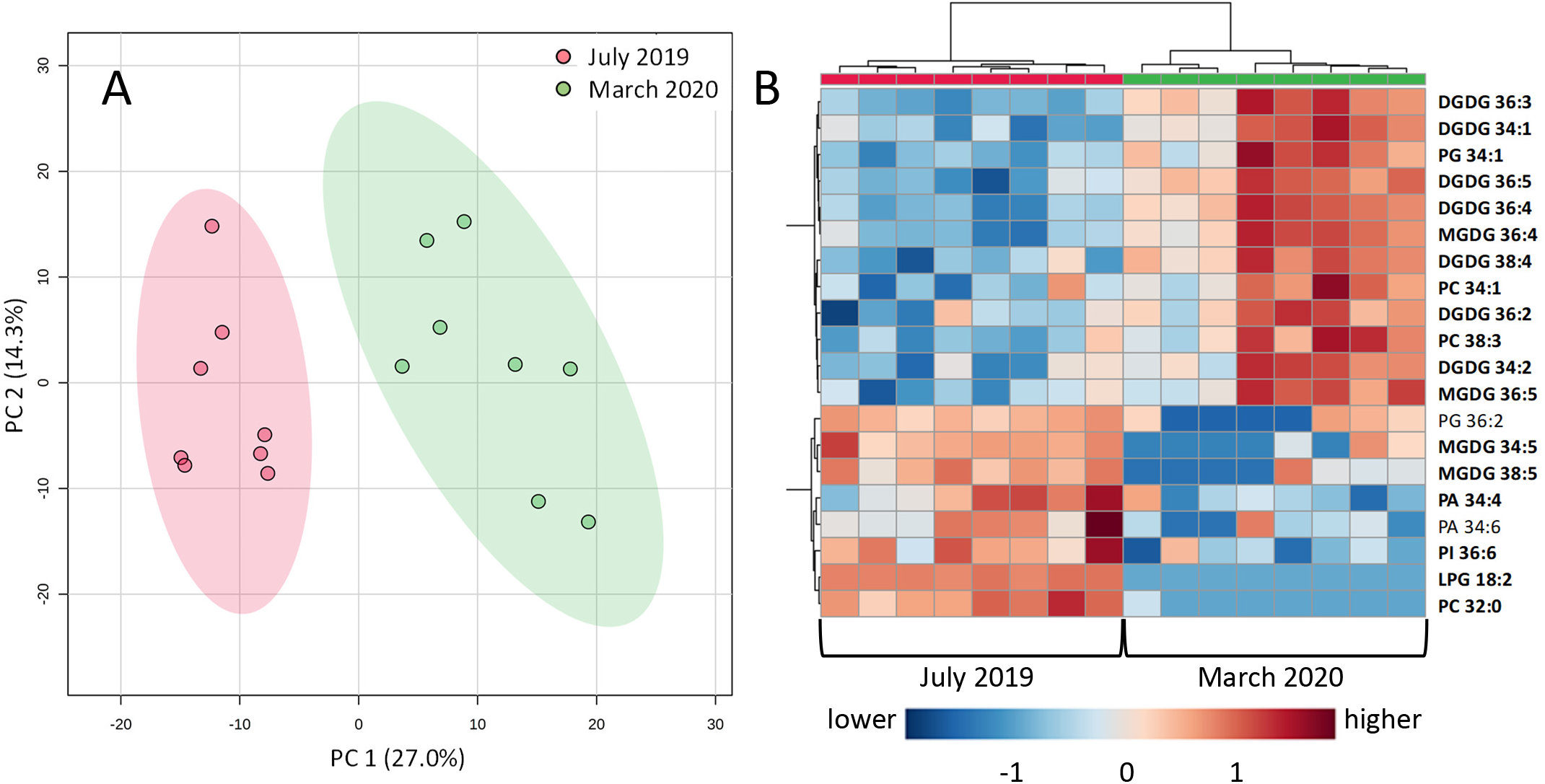
Polar lipid composition compared between xylem sap (n = 8) of *Geijera* sampled in July 2019 and March 2020. A. Results from a principal component analysis (PCA), with the first two components shown. Shaded areas are 95% confidence regions. B. Heatmaps for polar lipids in, based on Euclidean distance measures, a Ward clustering algorithm, and normalized data to visualize differences between sampling times. The lipids shown are the 17 most different between sampling times based on unpaired t-tests, adjusted for false discovery rates (FDR). All lipids in this heatmap were significantly different (p < 0.05) between dates after FDR adjustment.

### Comparisons of lipid compositions in wood and xylem sap (two species)

Wood samples include xylem sap lipids, so the samples are not independent and can be compared only qualitatively (Table 3). However, PCAs revealed some differences between wood and sap, especially for *Distictis* (Fig. 4 A), where FDR-adjusted t-tests identified five phospholipids (out of 125) as different between wood and sap. No headgroups, chain lengths, or unsaturation levels were consistently different between wood and sap for *Distictis*, but several types of PC stood out as being more abundant in sap than in wood, while several types of PS and lysophospholipids showed the opposite trend (Fig. 4 B). For *Geijera*, only one polar lipid, PI 34:3, was identified by t-tests as different between wood and sap (Fig. 4 B), and PE was slightly more abundant (by 3.3%) in wood than in sap, and lysoPC was slightly less abundant in wood than in sap (by −0.14%). PA was just as abundant in wood as in sap (Table 3), even though wood samples were treated immediately with hot isopropanol to inactive PLD enzyme activity. No galactolipids were flagged as being different between wood and sap of either species. Overall, differences in lipid composition between wood and sap were not pronounced.

**Table 3.** Comparison of lipid composition (in %) in paired samples (n = 4) of xylem sap and wood from the same stems collected in March 2020. All concentrations were converted to percentages of total polar lipids. Wood and sap samples are not independent from each other, so statistical comparisons were not possible.

|  | *Geijera* |  |  |  | *Distictis* |  |  |  |
| --- | --- | --- | --- | --- | --- | --- | --- | --- |
|  | Sap |  | Wood |  | Sap |  | Wood |  |
|  | mean | STD | mean | STD | Mean | STD | mean | STD |
| DGDG | 9.52 ± 0.84 |  | 7.23 ± 1.08 |  | 4.53 ± 1.93 |  | 6.05 ± 1.00 |  |
| MGDG | 7.74 ± 0.81 |  | 8.00 ± 1.54 |  | 4.33 ± 1.43 |  | 5.66 ± 0.78 |  |
| PG | 2.45 ± 0.55 |  | 5.78 ± 1.38 |  | 1.26 ± 0.30 |  | 2.73 ± 1.00 |  |
| LysoPG | 1.44 ± 1.66 |  | 0.16 ± 0.06 |  | 1.23 ± 0.75 |  | 0.50 ± 0.41 |  |
| LysoPC | 0.31 ± 0.15 |  | 0.18 ± 0.07 |  | 0.21 ± 0.07 |  | 0.50 ± 0.10 |  |
| LysoPE | 0.17 ± 0.08 |  | 0.15 ± 0.03 |  | 0.18 ± 0.08 |  | 0.30 ± 0.03 |  |
| PC | 11.64 ± 1.94 |  | 14.62 ± 5.38 |  | 23.55 ± 3.06 |  | 14.57 ± 3.23 |  |
| PE | 7.39 ± 2.40 |  | 10.66 ± 2.39 |  | 8.47 ± 1.37 |  | 9.01 ± 1.76 |  |
| PI | 3.73 ± 1.51 |  | 6.16 ± 0.83 |  | 7.35 ± 0.71 |  | 9.00 ± 0.87 |  |
| PS | 1.88 ± 0.62 |  | 1.57 ± 0.52 |  | 2.00 ± 0.77 |  | 1.68 ± 0.39 |  |
| PA | 53.74 ± 16.80 |  | 45.51 ± 4.85 |  | 46.88 ± 2.59 |  | 50.02 ± 3.85 |  |
| Total | 100 |  | 100 |  | 100 |  | 100 |  |

**FIG. 4.**
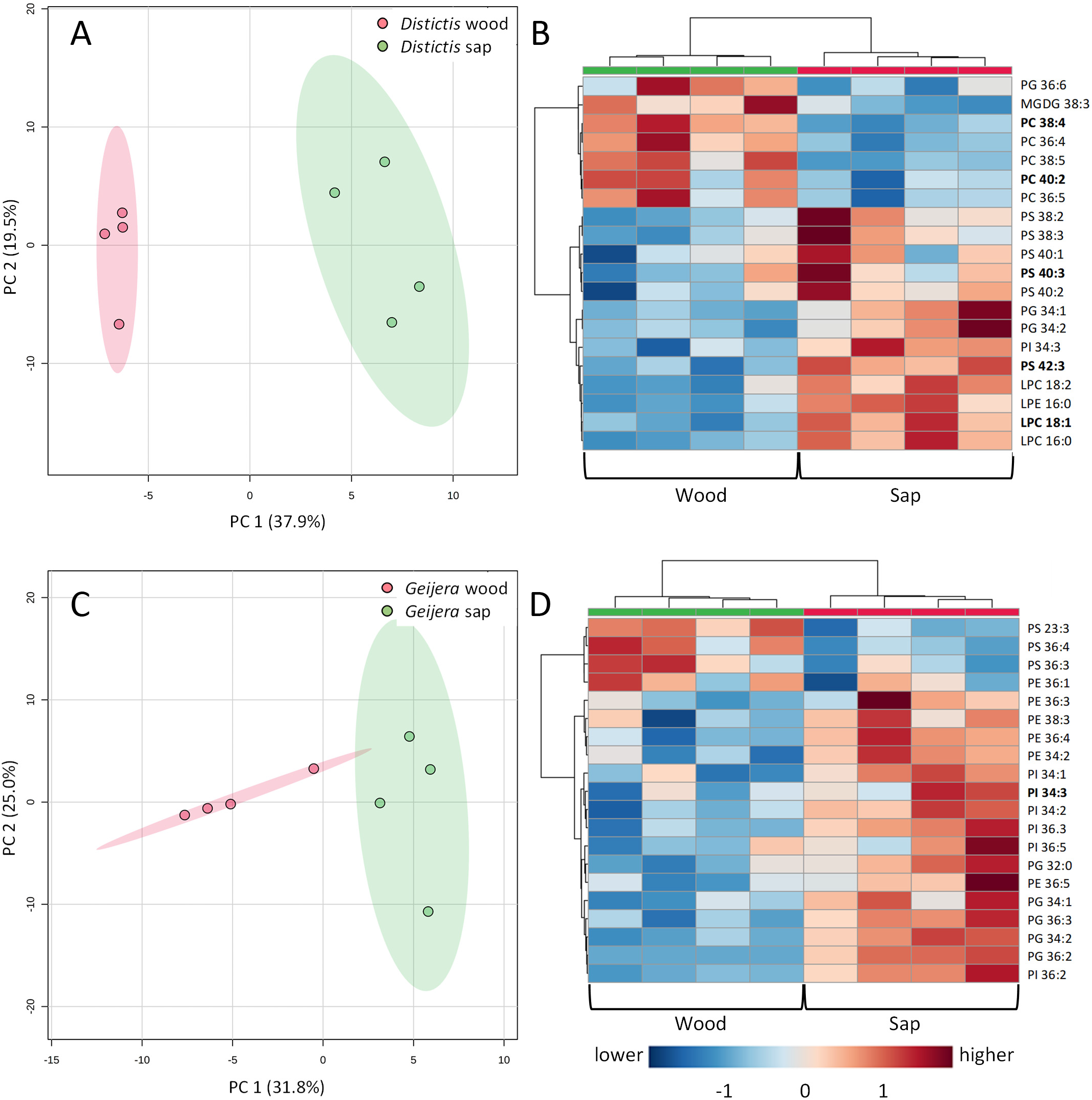
Polar lipid composition compared between paired samples of xylem sap and wood of the same stems in *Distictis* and *Geijera* (n = 4). A. and C. show results from principal component analyses (PCA) for *Distictis* and *Geijera*, respectively, with the first two components shown. Shaded areas are 95% confidence regions. B. and D. are heatmaps for polar lipids in *Distictis* and *Geijera*, respectively, based on normalized data, Euclidean distance measures, and a Ward clustering algorithm to visualize differences between xylem sap and wood. The lipids shown are the 20 most different between sap and wood based on paired t-tests, adjusted for false discovery rates (FDR). Lipids indicated in bold were flagged as different at *p* < 0.05 between xylem sap and wood after paired t-tests with FDR adjustment. Note that lipids in xylem sap and wood samples are not independent of each other, so t-tests were used only as qualitative indicators.

### *Visualization of lipids in xylem vessels of* Laurus

Treatment with FM1-43 showed continuous lipid layers on vessel surfaces of *Laurus nobilis*, with high concentrations in inter-vessel pits (Fig. 5 A-B). Pit membranes were clearly visible under confocal laser scanning microscopy as thick, pillow-shaped, black structures with lipid coatings on either side and some faint labelling inside the membranes. Never-dried pit membranes of *Laurus* were 1,252 nm (± 238 nm, standard deviation; n = 36 pit membranes) thick under confocal microscopy (Fig. 5 A-B), but 601 nm (± 150 nm; n = 19) under TEM (Fig. 5 D-G), showing that the dehydration required for TEM by propanol could result in shrinkage of the pit membrane. These observations were very similar to those published previously for five of the other study species, including *Liriodendron*, *Triadica*, *Geijera*, *Distictis*, and *Encelia* (Schenk *et al.*, 2018). TEM samples not treated with OsO_4_ showed a dark and homogeneous appearance (Fig. 5 D-E). After treatment with OSO_4_, the lipid layer appeared to be broken up into aggregates under TEM (Fig. 5 F-G) but was found in a continuous layer in the same locations observed under confocal microscopy on vessel and pit surfaces and on the pit membrane (Fig. 5 A-B).

**FIG. 5.**
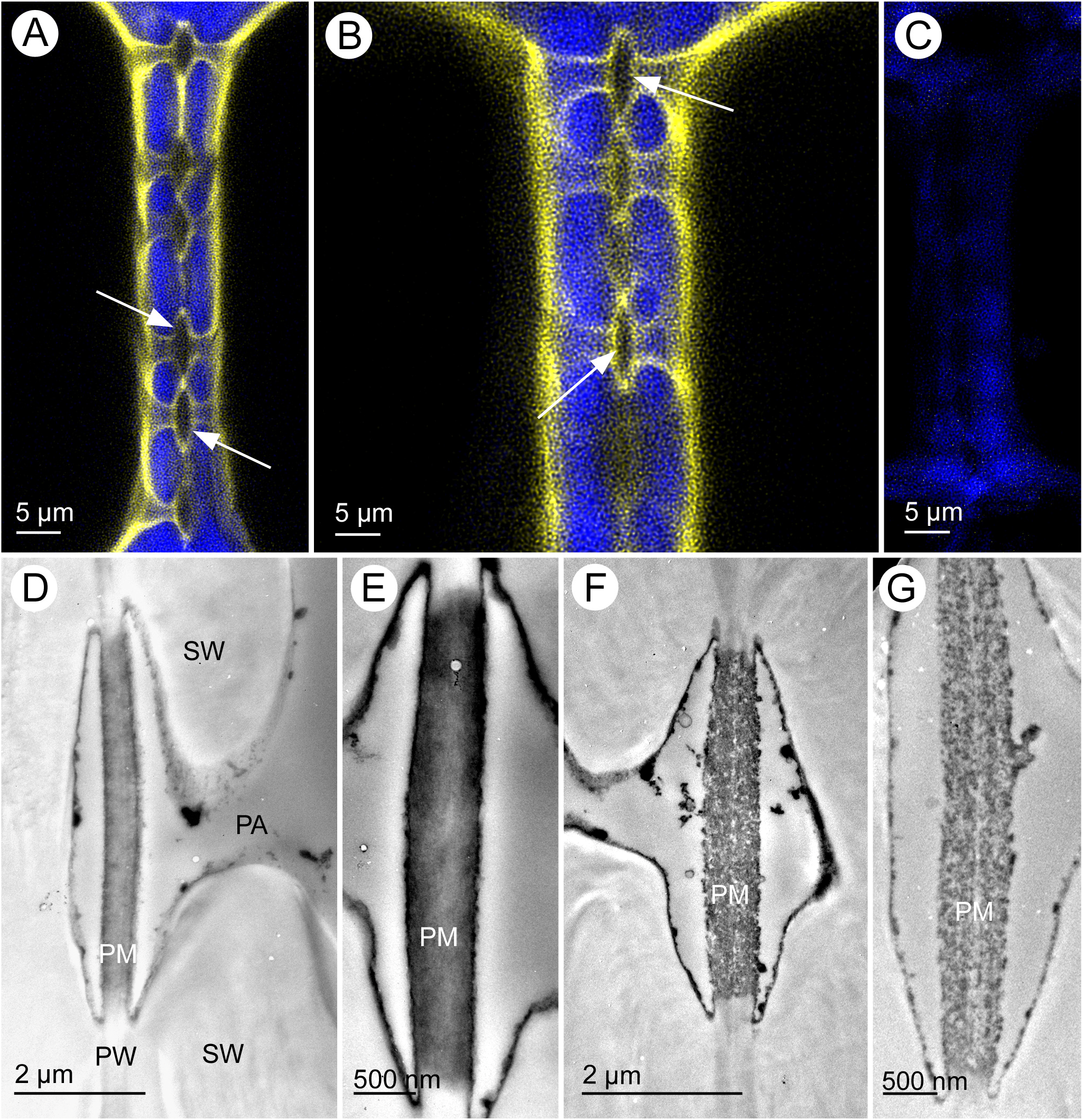
Lipids in xylem and inter-vessel pits of *Laurus nobilis*. A-C: Super-resolution confocal laser scanning microscopy images (false color) showing lignified cell walls in blue based on lignin autofluorescence and FM1-43 dye bound to lipids in yellow. White arrows = intervessel pit membranes. A-B: With FM1-43, C: Control without FM1-43. D-G: Transmission electron micrographs of intervessel pits. D-E: without OsO_4_; showing unusually dark, electron-dense homogeneous pit membranes and linings on secondary walls, but no granular appearance. F-G: with OsO_4_, lipids visible as black aggregates lining the wall surface, pit chamber and inside the pit membrane. Abbreviations: PA = pit aperture, PM = pit membrane, SW = secondary wall.

## Discussion

The most important take-home message of this study is that amphiphilic lipids exist in xylem conduits across the angiosperm phylogeny, including magnoliids, monocots, and eudicots. The available evidence based on our results indicate clearly that apoplastic lipids do not originate from cells cut open during sap extraction, as shown by contamination controls, lipid tracer experiments, and imaging. Phospholipids have been reported to occur in xylem sap (Gonorazky *et al.*, 2012, Schenk *et al.*, 2017), but not galactolipids. These findings raise questions about the chemical composition of xylem sap lipids, their origins, and functions of these lipids in angiosperm vessels.

### Chemical composition of xylem sap lipids

The concentrations of lipids in xylem sap were surprisingly consistent across species, with the two species from the magnoliid clade, *Laurus* and *Liriodendron*, having about a third of the concentrations of the other five species (Table 1; Fig. 2 A). Because lipid micelles or lipid-coated gas bubbles in xylem sap are mostly larger than 50 nm in diameter (Schenk *et al.*, 2017) and therefore too large to pass through pit membrane pores, which do not exceed 20 nm in hydrated pit membranes (Choat *et al.*, 2003, Zhang *et al.*, 2017, Zhang *et al.*, 2020), it is likely that the observed xylem sap lipids originate largely from vessels cut open for the sap extraction. To account for this, we analysed the opened vessel volumes of the seven species and found no significant relationship between lipid concentrations in sap and vessel volumes, but there was a significant negative relationship with the total vessel wall perimeter at the cut xylem surface (Fig. 2 C). It appears that the xylem functions like a filter during lipid extraction, so that xylem with higher wall surface area, i.e., more and smaller vessels, traps more surface-active lipids, leading to lower lipid yields in extracted sap. Because the open vessel volumes were typically smaller than the 1 to 2 mL of xylem sap extracted (except for the liana species *Distictis*), it is likely that the total amount of lipids in sap is underestimated due to a filtering and dilution effect.

Lipids are insoluble and in xylem conduits most are attached to surfaces (Fig. 4; Schenk *et al.*, 2018), so the amounts extracted in sap may only be a small fraction of all apoplastic xylem lipids. Quantification of all apoplastic lipids in xylem would require either complete extraction from individual conduits with organic solvents and/or detergents, which could cause artifacts if these substances penetrated into living vessel-associated cells (Morris *et al.*, 2018b), or quantification of lipids in vessel images via mass spectrometry imaging (Ellis *et al.*, 2013). The very low concentrations of lipids found in cell contamination controls from the basal xylem surface show that on average at least 95% of the lipids in xylem sap samples did not originate from cut parenchyma cells at basal xylem surfaces. *Bambusa* and *Distictis*, both of which have phloem embedded with xylem (*Distictis* has intraxylary phloem) had the lowest percentages of lipid concentrations in their control samples (4.3% in *Bambusa* and 1.1% in *Distictis*), demonstrating that hardly any lipids in xylem sap samples originated from the phloem. Lipid compositions of cell contamination controls were far more variable and different from those of sap (Fig. S12), which also confirms that sap lipids, except for traces, do not originate from cut living cells. Lipids in sap also clearly do not originate from cells cut at the distal end of stems, as lipid tracer applied to distal stem surfaces during xylem sap extraction did not show up in sap collections. Minute tracer amounts may have made their way through the exceptionally wide and long vessels of *Distictis*, but not enough to raise tracer levels significantly. Lipids from cut cells at the distal stem end are likely to attach to vessel surfaces as sap is extracted, or will be held up by inter-vessel pit membranes, instead of moving through >40 cm of vessel length and showing up in extracted sap.

The only other lipidomic analysis of xylem sap, for tomato (Gonorazky *et al.*, 2012), did not include analyses of galactolipids, but the chemical composition of phospholipids was roughly comparable to our findings, with PC and PA being the most abundant phospholipids. The xylem sap concentration of phospholipids in tomato (0.037 μmol kg^−1^) was similar to concentrations in *Laurus* (0.044 μmol L^−1^), but much lower than the average phospholipid concentrations across all species studied (0.296 ± 0.161 SD μmol L^−1^).

The presence of the galactolipids DGDG and MGDG in the xylem apoplast is especially notable, because galactolipids are synthesized exclusively in plastids (Dörmann and Benning, 2002, Botté and Maréchal, 2014) and are the most abundant lipids in chloroplasts, but they have never been found in any apoplastic compartment, presumably because there have been so few studies of “the black box of plant apoplast lipidomes” (Misra, 2016). Moreover, their physical properties, such as interactions of the head groups with water molecules, are different from phospholipids (Kanduč *et al.*, 2017). MGDG have very small head groups and form bilayers only in mixture with other lipids (Dörmann and Benning, 2002), where interactions between MGDG and DGDG head groups affect the physical properties of galactolipids at gas-water interfaces (Bottier *et al.*, 2007). It is also notable that sulfolipids, which are also abundant in plastids (Shimojima, 2011), were only found in trace amounts in xylem sap.

Regarding acyl chains, it was observed that galactolipids mostly had 36-carbon combinations, most likely consisting of two 18-carbon chains (Fig. 2 B). Some plants, termed “16:3 plants” can incorporate 16C-fatty acids into galactolipids and desaturate them to 16:3 (via the “prokaryotic pathway”), whereas others, termed “18:3 plants”, assemble the diacylglycerol backbone of galactolipids only in the E.R. 16:3 plants contain a mixture of 34C and 36C galactolipids, whether 18:3 plants contain only 36C galactolipids. Leaves of both *Laurus nobilis* and *Liriodendron tulipifera* contain small amounts (slightly over 1%) of plastid-synthesized 16:3 (Mongrand *et al.*, 1998). *Triadica*, *Geijera*, and *Distictis* leaves also may contain small amounts of 16:3, but *Bambusa* and *Encelia* are likely to be 18:3 plants. Still, the majority of xylem sap galactolipid backbones from all seven species are di-18C molecular species, consistent with synthesis in the ER (Mongrand *et al.*, 1998). Additionally, it has previously been shown that a 16:3 plant, such as *Arabidopsis thaliana*, favors the eukaryotic pathway in non-photosynthetic tissue (Devaiah *et al.*, 2006, Huynh *et al.*, 2012).

Along the same lines, the bulk of the observed lipid species appear to be less unsaturated than most leaf tissues, containing only minor amounts of 18:3 chains, which are the most prominent fatty acids in leaves. Whole tissue data demonstrate that generally *Arabidopsis* shoots are very high in 18:3, while roots and seeds are less unsaturated with more 18:2 and 18:1-containing galactolipids (Devaiah *et al.*, 2006, Huynh *et al*., 2012). Indeed, Gonorazky *et al.* (2011) observed that 18:3-containing phospholipids in leaves were less likely to be found in the extracellular fluid than in the whole leaf. The lipid composition of the xylem galactolipids, with 36:4 species prominent, suggests that 18:2 fatty acids may be common. Similarly, 34:2 species, which are likely 16:0/18:2 combinations are very common in phospholipids, suggesting an abundance of 18:2 in xylem sap.

Besides finding galactolipids in xylem sap, the presence of phosphatidic acid (PA) at high concentrations in some species is also notable (Fig. 2 A). High concentrations of PA in plant samples are usually attributed to phospholipase D (PLD) activity, which cleaves head groups such as the choline head-group of phosphatidylcholine (PC) off phospholipids to create PA (Christie, 1993, Welti *et al.*, 2002). PLD is active when bound to cell membranes (Kolesnikov *et al.*, 2012), and cutting living cells causes PLD activation as part of a wounding response (Bargmann *et al.*, 2009). High PA concentrations can be artifacts of wounding, and thorough cleaning of cut xylem surfaces is therefore essential for keeping fragments of cell membranes containing PLD out of xylem sap collections. PLD can remain active even in organic solvents such as chloroform:methanol (Christie, 1993), possibly because membrane fragments form inverted vesicles in organic solvents that allow for continued PLD activity in the hydrophilic interior of the vesicle, as has been suggested for phospholipase C (PLC) (Kates, 1957).

The high PA concentrations found in xylem sap of some species, such as *Encelia*, *Geijera*, and *Distictis* (Fig. 2 A) were initially a concern, because it was suspected that they could be sampling artifacts. However, treatments of cut xylem surfaces with the PLD inhibitors 1-butanol, 5-fluoro-2-indolyl des-chlorohalopemide (FIPI) (Su *et al.*, 2009), and a lipase inhibitor cocktail (Furse *et al.*, 2013) had no effect on PA concentrations in xylem sap of *Distictis* (data not shown). Moreover, PLD or other phospholipases have never been detected in proteomic analyses of xylem sap (Alvarez *et al.*, 2006, Djordjevic *et al.*, 2007, Aki *et al.*, 2008, Krishnan *et al.*, 2011, Ligat *et al.*, 2011, Dugé de Bernonville *et al.*, 2014), which may not be surprising, because PLD is often bound to membranes and therefore unlikely to occur naturally in xylem sap. Moreover, thin wood samples also contained high amounts of PA (Table 3), even though the slices were immediately dropped into hot isopropanol to inactivate PLD.

These lines of argument all support PA as a major natural ingredient of xylem saps, except for that of *Laurus* (Fig. 2 A; Table 1). PA has also been found in relatively high concentrations in xylem sap of tomato plants (Gonorazky *et al.*, 2012). PA has many functions in plants, interacts with many proteins, and has important roles in lipid metabolism and signalling (Testerink and Munnik, 2011), wounding responses (Bargmann *et al.*, 2009), and in responses to stress, such as drought and salinity (McLoughlin and Testerink, 2013). The physical properties of PA are also unique when compared to other amphiphilic lipids in having the smallest and negatively charged head-group, binding of cations such as Ca^2+^ and Mg^2+^ (Ohki and Ohshima, 1985), and causing PA to have lower surface tension in monolayers than other phospholipids (Weschayanwiwat *et al.*, 2005). Of all the various roles of PA, its interactions with cations and effects on surface tension at gas interfaces are likely to be most important in the xylem apoplast environment.

The amounts of choline-containing phospholipids (PC and LPC) in xylem sap reported here were much lower than estimated in previous research using an enzymatic assay (Schenk *et al.*, 2017), where all choline quantified in xylem sap was assumed to originate via PLD activity from choline-containing phospholipids. However, that assumption appears to have been incorrect. Choline has been found in xylem sap at molar concentrations far exceeding those of lipids (Lima *et al.*, 2017) and is likely to originate from sources other than phospholipids in sap.

### Where do apoplastic xylem lipids come from?

There are two possible and not mutually exclusive origins for apoplastic xylem lipids: Lipids have been shown to remain from the living cell content of conduits during conduit development (Scott *et al.*, 1960, Esau *et al.*, 1966). If this includes lipids from plastids, then this would easily explain the presence of galactolipids in xylem sap. Lipids could also be transported into conduits from conduit-associated parenchyma cells (Czaninski, 1977, Morris *et al.*, 2018b), but it seems unlikely that plastid lipids would be moved into vessels. We compared the lipid composition in sap to that of the wood that the sap was extracted from (Fig. 4) and found only minor differences in composition and none in galactolipids. To some degree, this is expected, as wood includes sap, so it is impossible for sap to include any lipids not found in wood samples. Out of 157 polar lipids tested, only one for *Geijera* and five for *Distictis* stood out as different in relative amounts between sap and wood (Table 2). Overall, the differences between sap and wood were small, so it seems reasonable to conclude that sap and wood lipids are essentially the same, supporting the origin of apoplastic xylem lipids, including galactolipids, from living vessel contents, as shown previously in microscopic studies of developing vessels (Scott *et al.*, 1960, Esau *et al.*, 1966).

That said, there were clear seasonal differences in the polar lipid composition of *Geijera* xylem sap between July and March (Fig. 3), with elevated galactolipid concentrations in March. Seasonal changes could be a reflection of wood development, with newer vessels containing different lipids than older ones, could be caused by apoplastic enzyme activities, or by lipid transport from vessel-associated cells. This would require transport of lipids across a cell membrane, the protective layer surrounding such cells, which consists mainly of polysaccharides (Chafe and Chauret, 1974, Fujii *et al.*, 1981, Wisniewski *et al.*, 1991a, Wisniewski *et al.*, 1991b, Wisniewski and Davis, 1995), as well as across the pit membrane, which mainly consists of cellulose. The transport could involve non-specific lipid transfer proteins (nsLTPs), which can transport lipids across cell walls (Domínguez *et al.*, 2015, Fich *et al.*, 2016, Li *et al.*, 2016, Misra, 2016) and which have been found in xylem cell walls and xylem sap (Buhtz *et al.*, 2004, Djordjevic *et al.*, 2007, Krasikov *et al.*, 2011, Ligat *et al.*, 2011). However, it is exceedingly unlikely that plastid lipids would be transported into vessels, so seasonal differences in galactolipids in xylem sap are more likely to be caused either by apoplastic enzyme activities or by immobilization of galactolipids on vessel surfaces and in pit membranes, which would prevent them from being extracted in sap. Further research will be required to understand these seasonal changes.

Lipid transport or secretion of apoplastic lipid-altering enzymes from conduit-associated cells would allow for changes in lipid concentration and composition in conduits seasonally or in response to environmental conditions, which could possibly affect hydraulic functions and embolism resistance (Charrier *et al.*, 2018). Plants species vary widely in the amount and degree of contact between conduits and living parenchyma cells in secondary xylem (Martínez-Cabrera *et al.*, 2009, Morris and Jansen, 2016, Morris *et al.*, 2016, Morris *et al.*, 2018a), and it could be that some species undergo seasonal changes in apoplastic lipids while others do not. Seasonal changes in the electron-density of pit membranes have been observed (Schmid and Machado, 1968), and this could potentially indicate lipid accumulation in conduits over time.

### Apoplastic xylem lipids and functional implications

The presence of lipids on xylem conduit surfaces and pit membranes is incompatible with the widely accepted assumption that gas bubble formation in xylem is prevented by the high surface tension of water (Oertli, 1971, Sperry and Tyree, 1988, Tyree and Zimmermann, 2002, Stroock *et al.*, 2014). There is unequivocal evidence for negative pressure in xylem, and this aspect of the CT theory is not in question, although many open questions remain (Jansen and Schenk, 2015, Schenk *et al.*, 2015, Schenk *et al.*, 2017, Venturas *et al.*, 2017). Molecular dynamics modelling of lipid bilayers under negative pressure confirmed that cavitation in lipid bilayers and micelles is unlikely to occur within the normal range of pressures (above −10 MPa) experienced in plants (Kanduč *et al.*, 2020).

However, layers of phospho- and galactolipids on conduit surfaces and in pit membranes (Fig. 5) will reduce the surface tension at gas-water interfaces substantially below that of pure water, with the dynamic surface tension depending on the local concentration of lipids at the surface (Zuo *et al.*, 2004, Kwan and Borden, 2010). Measurements of xylem lipids extracted from four of the species studied in this paper found average equilibrium surface tensions to be about 25 mN/m, roughly a third of the surface tension of pure water (Yang *et al.*, 2020). These findings are entirely compatible with the air seeding pressure required to force air through pore constrictions in angiosperm pit membranes. Pore constrictions rarely exceed 20 nm in diameter (Zhang *et al.*, 2020), and it would take about 7 MPa of pressure to force air through such a constriction if surface tension were that of pure water (Kaack *et al.*, 2019). Actual air seeding pressures through pit membranes tend to be in the range of −0.4 to −2 MPa (Bartlett *et al.*, 2016), thereby supporting a role for lipids in pit membranes in the air seeding process.

In an earlier paper, we proposed a hypothesis to reconcile the presence of apoplastic xylem lipids with the CT theory (Schenk *et al.*, 2015) and later amended and modified that hypothesis in the light of new data (Schenk *et al.*, 2017). To summarize that hypothesis briefly, gas penetrating from embolized conduits into pit membranes of sap-filled conduits may snap off nanobubbles inside confined, lipid-coated pore spaces, resulting in the creation of lipid-coated nanobubbles (Fig. 6), which have in fact been found and visualized in xylem sap (Schenk *et al.*, 2017). Key to this hypothesis is that both the low dynamic surface tension of lipids and snapping off of bubbles at pore constrictions inside fibrous pit membranes would limit the bubbles’ sizes (Park *et al.*, 2019) and keep them below a critical threshold that prevents bubble expansion into embolism (Schenk *et al.*, 2015, Schenk *et al.*, 2017). This process could happen under normal xylem water potentials and would allow xylem to operate without spreading embolisms via air seeding from gas-filled conduits. The population of lipid-coated nanobubbles could also function as a gas reservoir that serves as a buffer when dissolved gas in sap exceeds saturation under increasing temperatures (Schenk *et al.*, 2016).

**FIG. 6.**
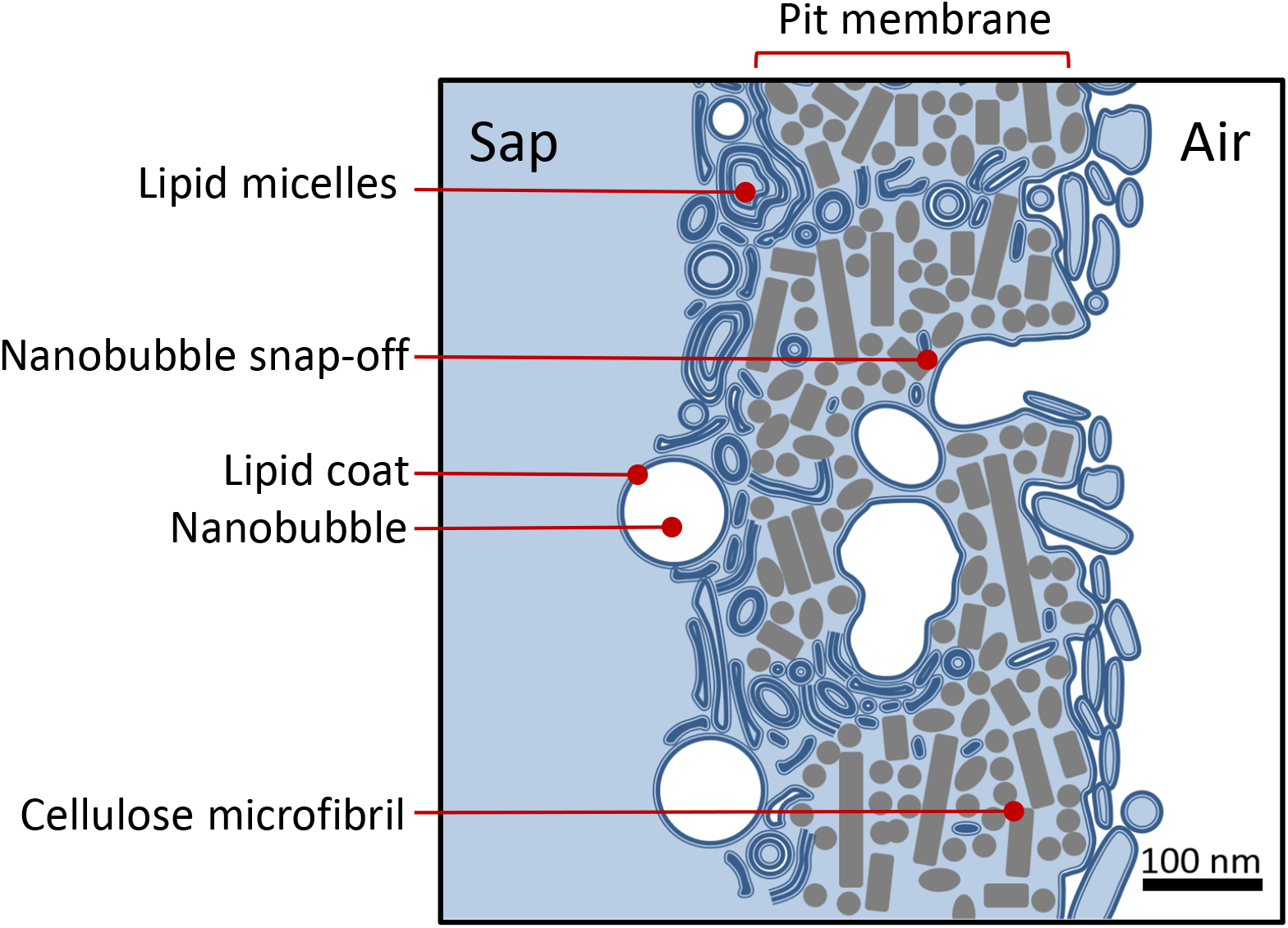
Conceptual model of the formation of lipid-coated nanobubbles via snap-off inside pit membranes that are coated and infiltrated with surface-active lipids. The different sizes of micelles shown in this image reflect the wide range of nanoparticle sizes detected in xylem sap and presumed to be bilayer to multilayer micelles, vesicles, and lipid-coated nanobubbles (Schenk *et al*. 2017).

Below a more negative air seeding pressure threshold, gas could potentially still penetrate through the entire length of pit membrane pores, causing a growing bubble to emerge from the pit membrane, where nothing would restrict the bubble’s growth and resulting in embolism. This is the widely accepted air seeding process (Sperry and Tyree, 1988, Tyree and Zimmermann, 2002, Stroock *et al.*, 2014, Brodribb *et al.*, 2016), the underlying mechanism of which is still poorly understood (Jansen *et al.*, 2018), but which is supported by a large body of research, including measurements of pit membrane pore constrictions and air seeding pressures (Kaack *et al.*, 2019). What the new lipid-nanobubble hypothesis could explain is how plants can operate under negative pressure at all without forming embolism even under mildly negative pressure, a feat that engineers have found almost impossible to replicate (Smith, 1994). If this hypothesis, which is not contradictory but complementary to the cohesion-tension theory, is correct, then lipids are required for water transport under negative pressure. In fact, Dixon (1914) already suggested the colloids increase the tensile strength of xylem sap over that of pure water.

## Conclusions

This study provides conclusive evidence for substantial amounts of phospho- and galactolipids in xylem conduits across the angiosperm phylogeny, and these findings force major questions about the functions of these lipids for plant water transport and the cohesion tension theory, as well as about the origins of lipids. Research on the functions of lipids in water that is under negative pressure and in the presence of gas-water interfaces in pit membranes and bubbles will benefit from an interdisciplinary approach to involve surface scientists and physicists with expertise on the behavior of lipids at surfaces such as pulmonary surfactant layers (Zuo *et al.*, 2008, Zhang *et al.*, 2011, Kanduč *et al.*, 2013, Kanduč *et al.*, 2016). Moreover, possible interactions between lipids and proteins, including enzymes, in xylem conduits need to be investigated. Even in the absence of gas in conduits, lipids accumulated in pit membranes are likely to affect sap flow and could be involved in the regulation of hydraulic conductance, the so-called ionic effect (Nardini *et al.*, 2011, Nardini *et al.*, 2012). Finally, if lipids are crucial for plant water transport, then they should be found as well in gymnosperms, ferns, and perhaps even moss hydroids. The discovery, or rather rediscovery (Scott *et al.*, 1960, Esau, 1965, Esau *et al.*, 1966, Wagner *et al.*, 2000), of lipids in xylem conduits opens a vast field of new research questions.

## Supporting information

Supplemantary information combined into one file

## Funding

The research was funded by research grants from the National Science Foundation (EAGER IOS-1558108 and IOS-1754850). SJ and LK acknowledge funding from the German Research Foundation (DFG, project No. 383393940).

## Acknowledgements

The authors thank Anne Basilio, Zoe Cuevas, Jessica Garcia, Ryan Cochoit, Alec Hunt, and Tilly Duong for assistance with xylem sap extractions and vessel volume analyses, Jim Henrich at the Los Angeles County Arboretum and Greg Pongetti at the Fullerton Arboretum for allowing sampling of plants in their living collections, the Electron Microscopy Section of Ulm University for preparing TEM samples, and the Core Facility for Confocal and Multiphoton Microscopy of Ulm University for assistance with confocal imaging.

## Supplementary Materials

### Supporting figures

Figure S1. . Xylem sap concentrations of digalactosyldiacylglycerol (DGDG) in seven angiosperms species.

Figure S2. Xylem sap concentrations of monogalactosyldiacylglycerol (MGDG) in seven angiosperms species.

Figure S3. Xylem sap concentration of phosphatidylglycerol (PG) in seven angiosperms species.

Figure S4. Xylem sap concentration of phosphatidylcholine (PC) in seven angiosperms species.

Figure S5. Xylem sap concentration of phosphatidylethanolamine (PE) in seven angiosperms species.

Figure S6. Xylem sap concentrations of phosphatidylinositol (PI) in seven angiosperms species.

Figures S7. Xylem sap concentrations of phosphatidylserine (PS) in seven angiosperms species.

Figures S8. Xylem sap concentrations of phosphatidic acid (PA) in seven angiosperms species.

Figure S9. Xylem sap concentrations of lysophosphatidylethanolamine (LPE) and lysophosphatidylcholine (LPC) in seven angiosperms species.

Figure S10. Stems infiltrated with blue-colored silicone, cut at 0.7 cm from the injected surface. The four spoke-like structures in *Distictis buccinatoria* xylem are areas of interxylary phloem.

Figure S11. Total vessel area filled with silicone as a function of distance from the infiltrated surface, shown for a sample of *Triadica sebifera*. The silicone-filled vessel volume was calculated from these filled areas by linear interpolation.

Fig. S12. Principal component analyses (PCA) comparing polar lipid composition in xylem sap for six angiosperm species to paired cell contamination controls from the same stems.

### Supporting tables

Table S1. Standards used for mass spectrometry of xylem sap lipids.

Table S2. Concentrations of polar lipids and sulfolipids in xylem sap.

Table S3. Concentrations of polar lipids and sulfolipids in cell contamination controls for xylem sap.

### Supporting experimental procedures

Methods S1. Methods comparison of lipid extraction by partial freeze drying vs. SpeedVac.

## Notes

### Competing Interest Statement

The authors have declared no competing interest.

### Summary of Updates

This revision includes new results and analyses not included in the previous version, presented in new figures, tables, and supplementary information. Coauthor Kerri Mocko has been added.

