## Supplementary material for "Lipids in xylem sap of woody plants across the angiosperm phylogeny": Supplemantary information combined into one file

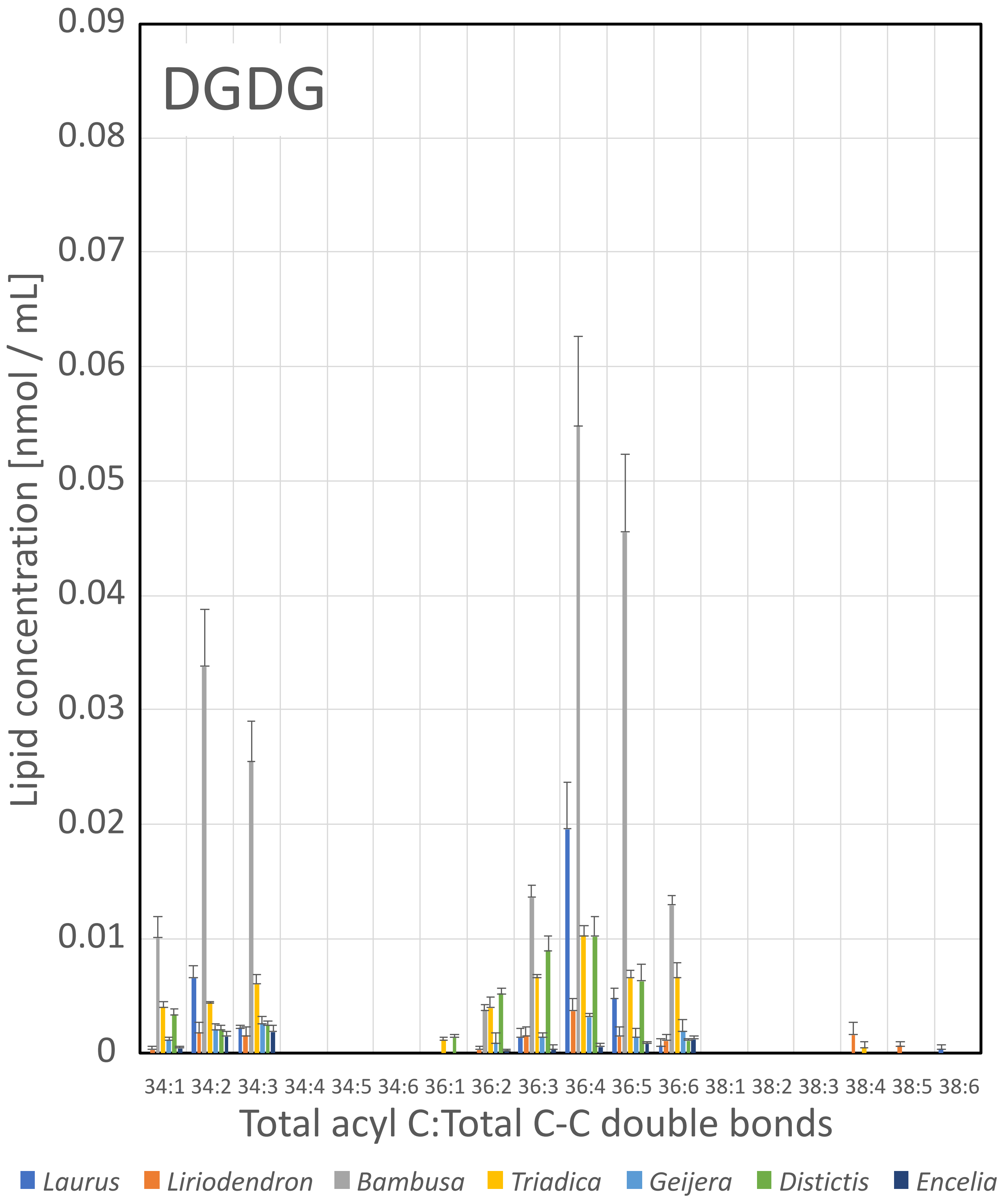
DGDG
Lipid concentration [nmol / mL]
0.09
0.08
0.07
0.06
0.05
0.04
0.03
0.02
0.01
0
34:1 34:2 34:3 34:4 34:5 34:6 36:1 36:2 36:3 36:4 36:5 36:6 38:1 38:2 38:3 38:4 38:5 38:6
Total acyl C:Total C-C double bonds
Laurus
Liriodendron
Bambusa
Triadica
Geijera
Distictis
Encelia

Fig. S1.

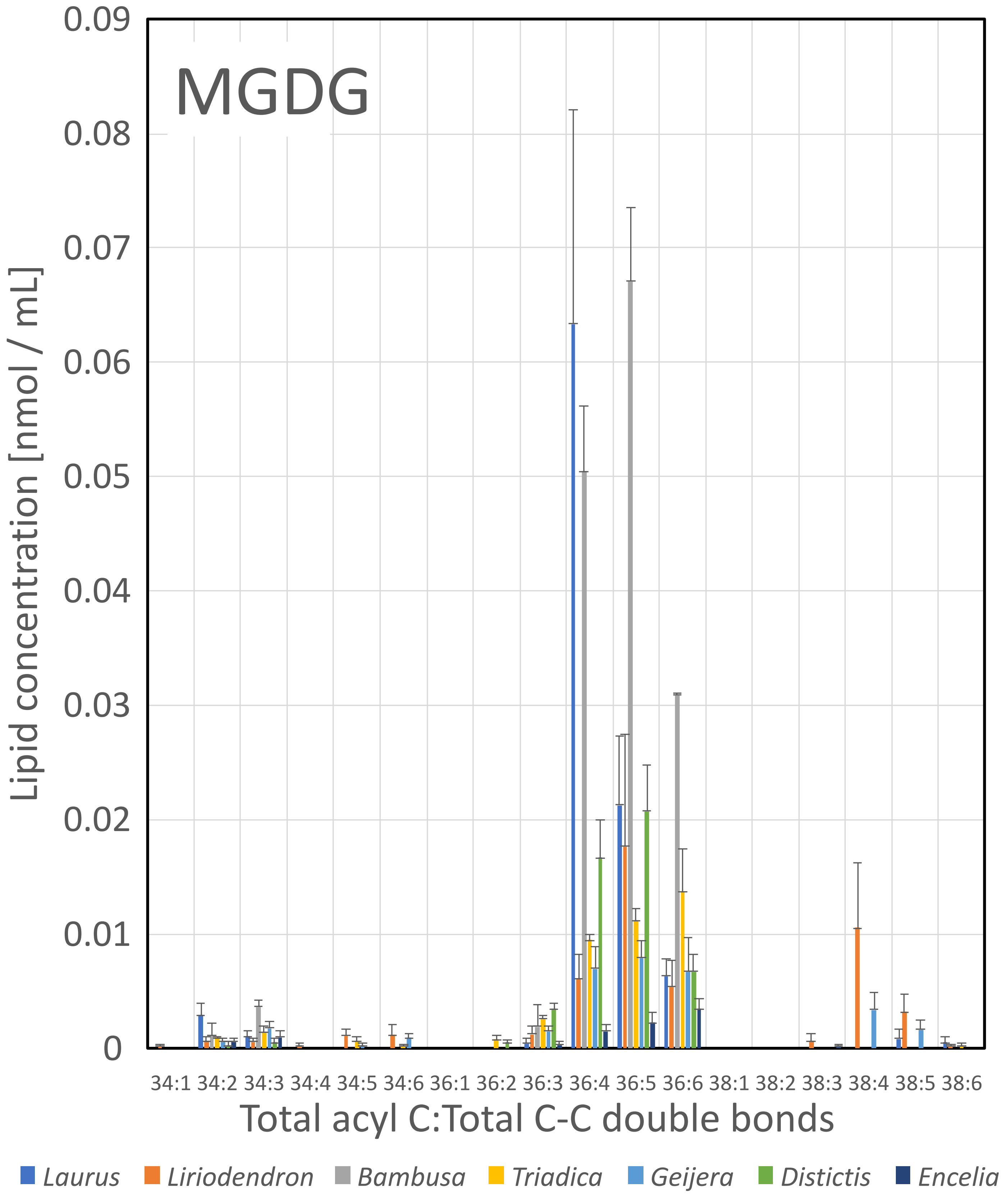
MGDG
Lipid concentration [nmol / mL]
0.09
0.08
0.07
0.06
0.05
0.04
0.03
0.02
0.01
0
34:1 34:2 34:3 34:4 34:5 34:6 36:1 36:2 36:3 36:4 36:5 36:6 38:1 38:2 38:3 38:4 38:5 38:6
Total acyl C:Total C-C double bonds
Laurus
Liriodendron
Bambusa
Triadica
Geijera
Distictis
Encelia

Fig. S2

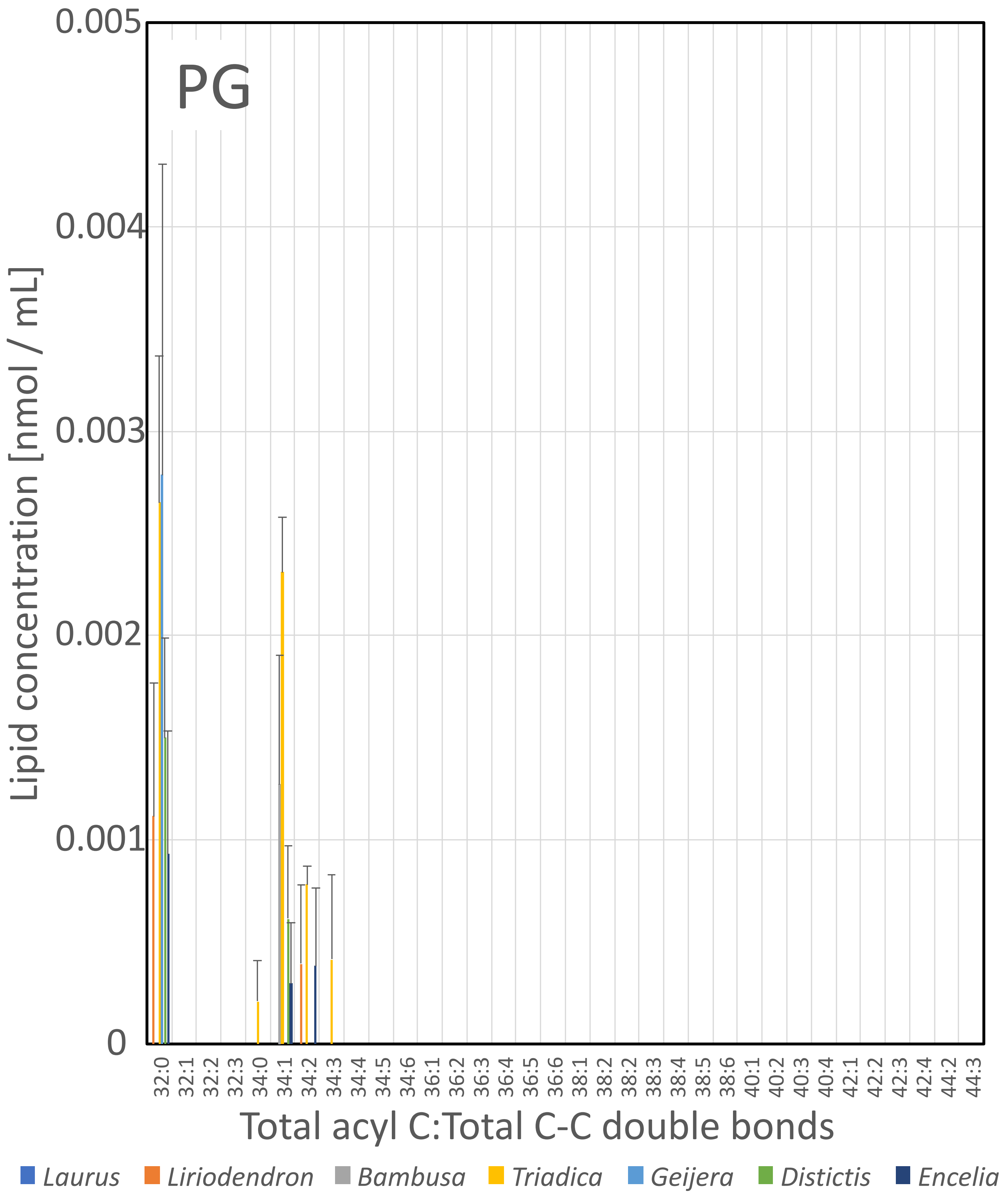
PG
0.005
0.004
0.003
0.002
0.001
0
Lipid concentration [nmol / mL]
32:0 32:1 32:2 32:3 34:0 34:1 34:2 34:3 34:4 34:5 34:6 36:1 36:2 36:3 36:4 36:5 36:6 38:1 38:2 38:2 38:3 38:4 38:5 38:6 40:1 40:2 40:3 40:4 42:1 42:2 42:3 42:4 44:2 44:3
Total acyl C:Total C-C double bonds
Laurus
Liriodendron
Bambusa
Triadica
Geijera
Distictis
Encelia

Fig. S3

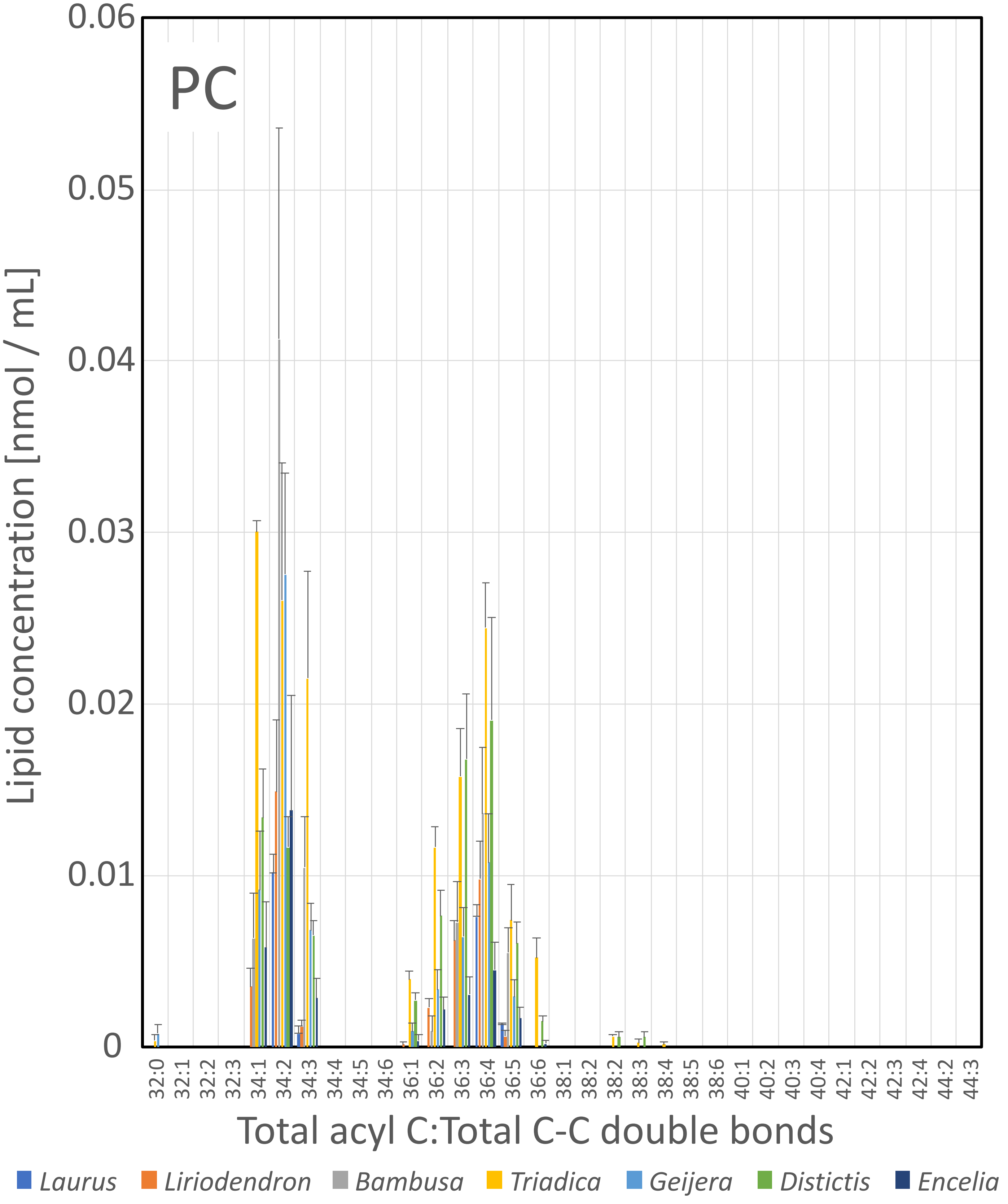
PC
Lipid concentration [nmol / mL]
0.06
0.05
0.04
0.03
0.02
0.01
0
32:0
32:1
32:2
32:3
34:1
34:2
34:3
34:4
34:5
34:6
36:1
36:2
36:3
36:4
36:5
36:6
38:1
38:2
38:2
38:3
38:4
38:5
38:6
40:1
40:2
40:3
40:4
42:1
42:2
42:3
42:4
44:2
44:3
Total acyl C:Total C-C double bonds
Laurus
Liriodendron
Bambusa
Triadica
Geijera
Distictis
Encelia

Fig. S4

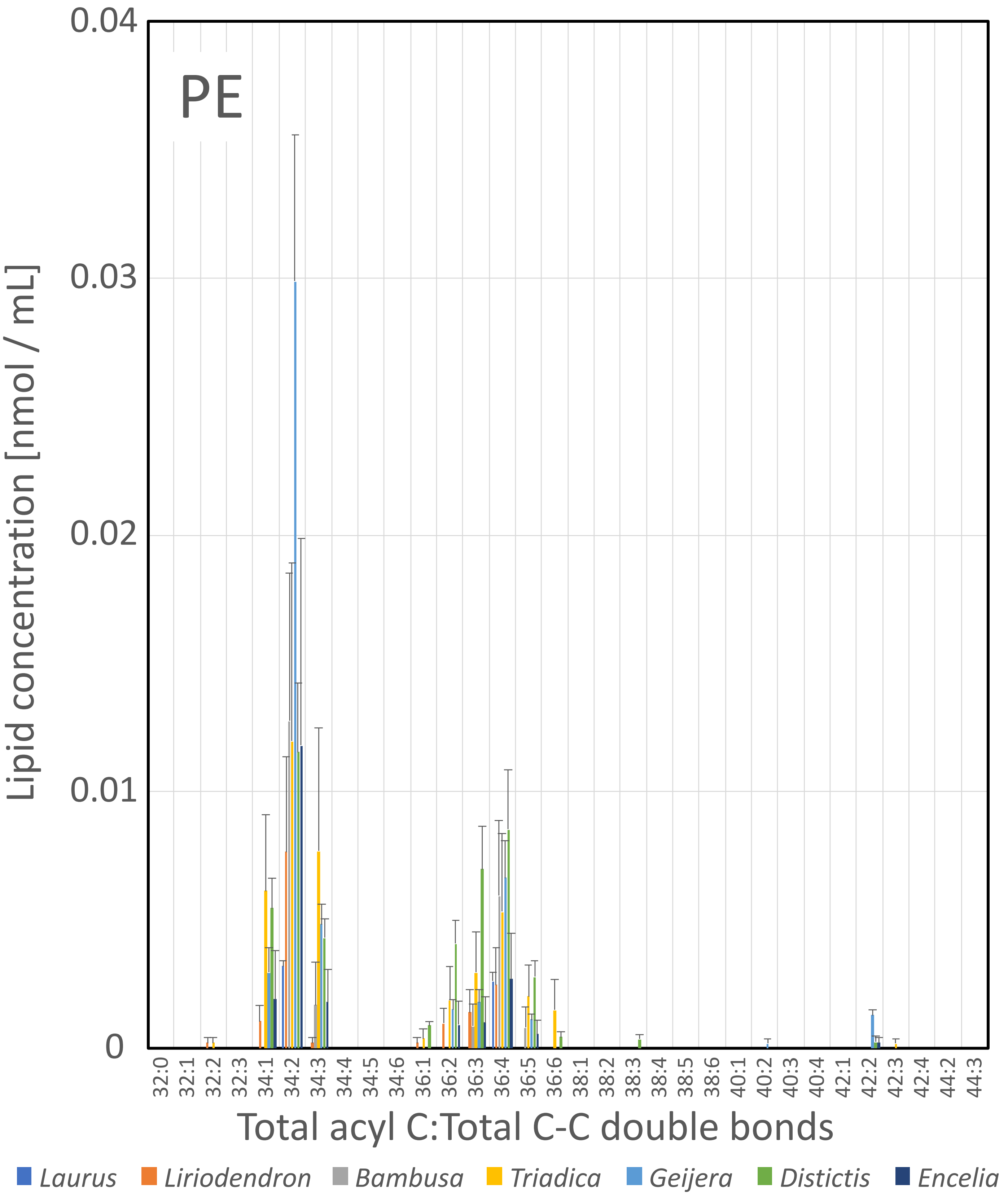

PE
Lipid concentration [nmol / mL]
0.04
0.03
0.02
0.01
0
32:0
32:1
32:2
32:3
34:1
34:2
34:3
34:4
34:5
34:6
36:1
36:2
36:3
36:4
36:5
36:6
38:1
38:2
38:3
38:4
38:5
38:6
40:1
40:2
40:3
40:4
42:1
42:2
42:3
42:4
44:2
44:3
Total acyl C:Total C-C double bonds
Laurus
Liriodendron
Bambusa
Triadica
Geijera
Distictis
Encelia


Fig. S5

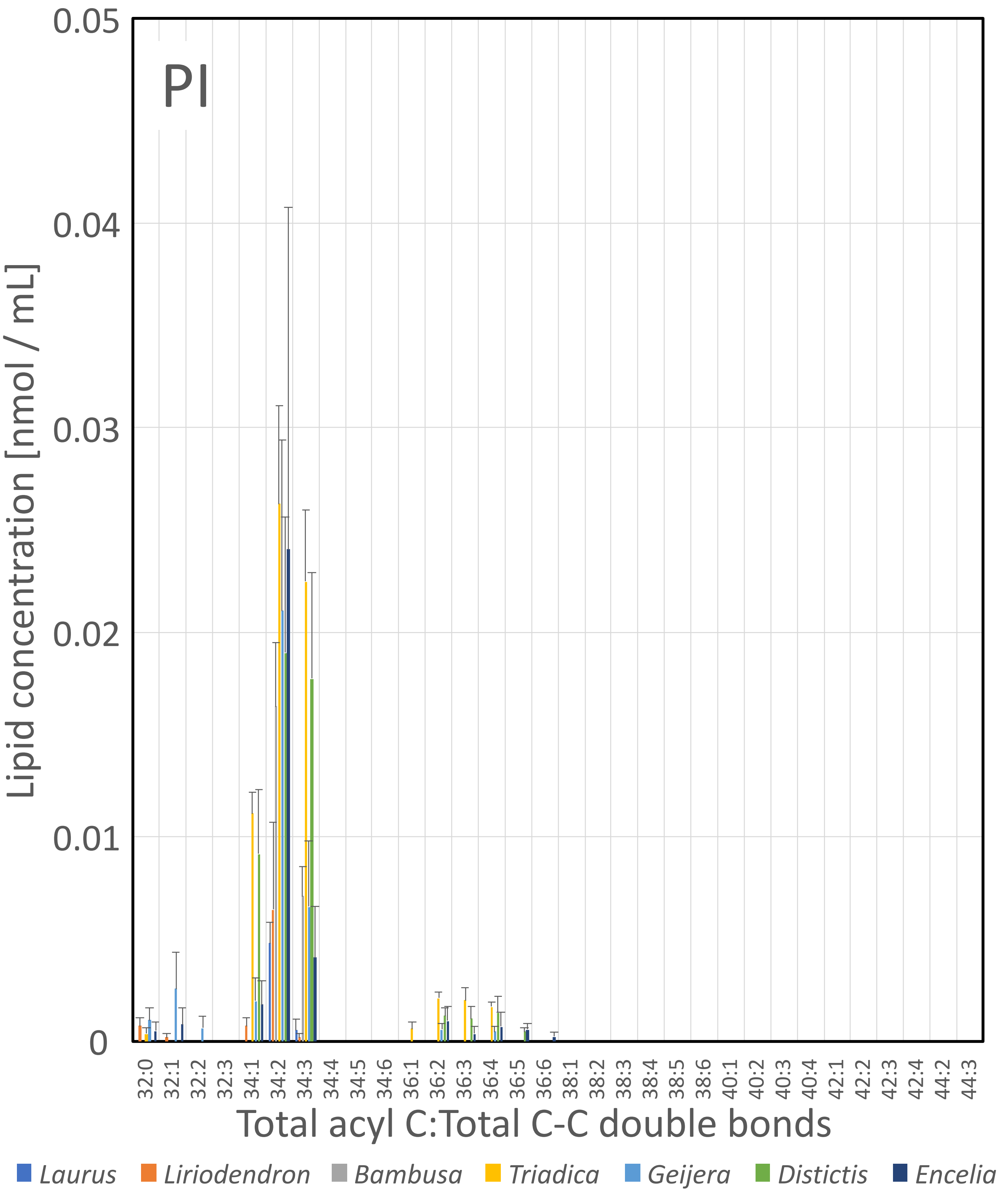

PI
0.05
0.04
0.03
0.02
0.01
0
Lipid concentration [nmol / mL]
32:0
32:1
32:2
32:3
34:1
34:2
34:3
34:4
34:5
34:6
36:1
36:2
36:3
36:4
36:5
36:6
38:1
38:2
38:3
38:4
38:5
38:6
40:1
40:2
40:3
40:4
42:1
42:2
42:3
42:4
44:2
44:3
Total acyl C:Total C-C double bonds
Laurus
Liriodendron
Bambusa
Triadica
Geijera
Distictis
Encelia


Fig. S6

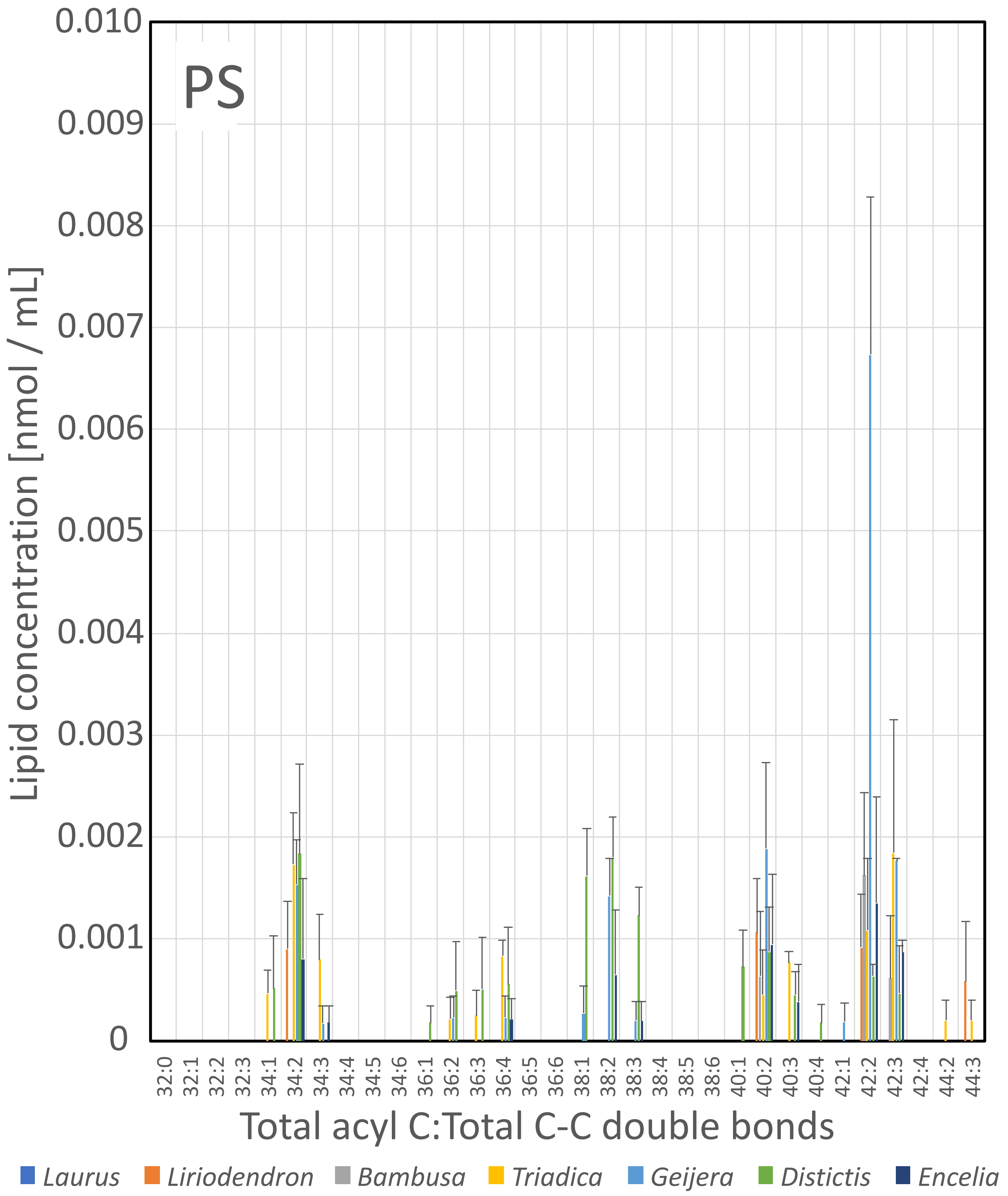
PS
0.010
0.009
0.008
0.007
0.006
0.005
0.004
0.003
0.002
0.001
0
Lipid concentration [nmol / mL]
32:0
32:1
32:2
32:3
34:1
34:2
34:3
34:4
34:5
34:6
36:1
36:2
36:3
36:4
36:5
36:6
38:1
38:2
38:3
38:4
38:5
38:6
40:1
40:2
40:3
40:4
42:1
42:2
42:3
42:4
44:2
44:3
Total acyl C:Total C-C double bonds
Laurus
Liriodendron
Bambusa
Triadica
Geijera
Distictis
Encelia

Fig. S7

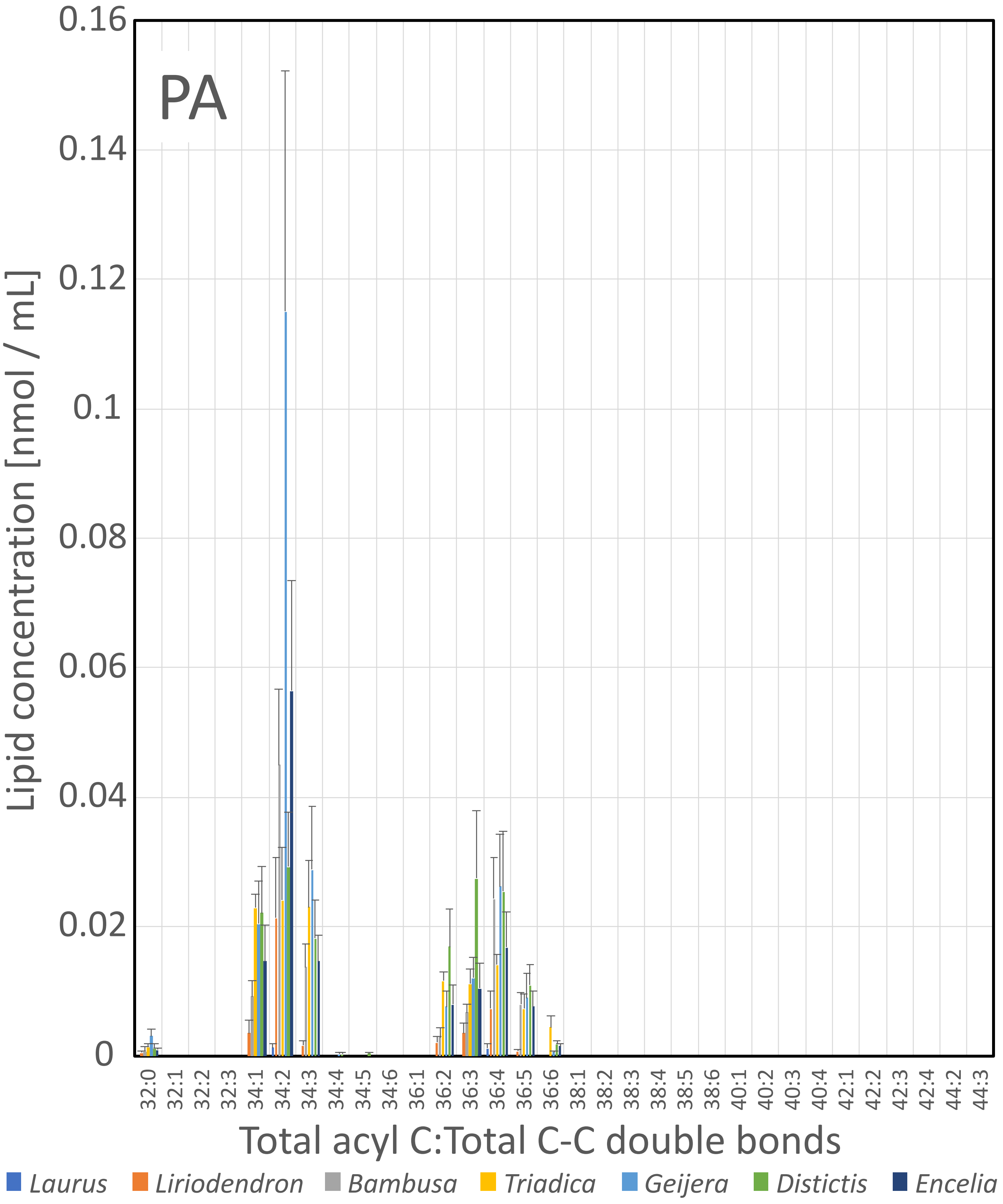
PA
Lipid concentration [nmol / mL]
0.16
0.14
0.12
0.1
0.08
0.06
0.04
0.02
0
32:0
32:1
32:2
32:3
34:1
34:2
34:3
34:4
34:5
34:6
36:1
36:2
36:3
36:4
36:5
36:6
38:1
38:2
38:3
38:4
38:5
38:6
40:1
40:2
40:3
40:4
42:1
42:2
42:3
42:4
44:2
44:3
Total acyl C:Total C-C double bonds
Laurus
Liriodendron
Bambusa
Triadica
Geijera
Distictis
Encelia

Fig. S8

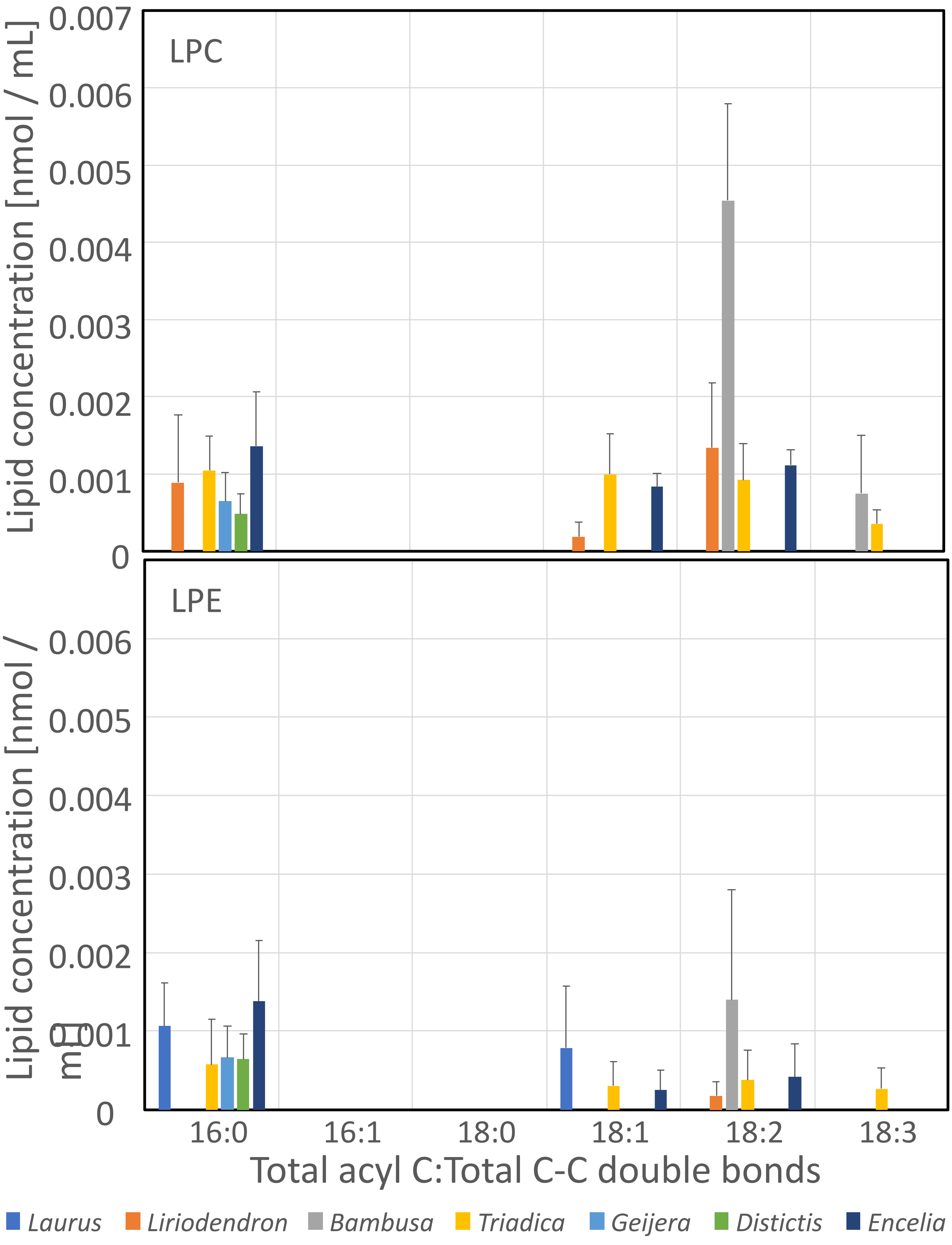

LPC
LPE
Lipid concentration [nmol / mL]
Lipid concentration [nmol / mL]
0.007
0.006
0.005
0.004
0.003
0.002
0.001
0
16:0
16:1
18:0
18:1
18:2
18:3
Total acyl C:Total C-C double bonds
Laurus
Liriodendron
Bambusa
Triadica
Geijera
Distictis
Encelia


Fig. S9

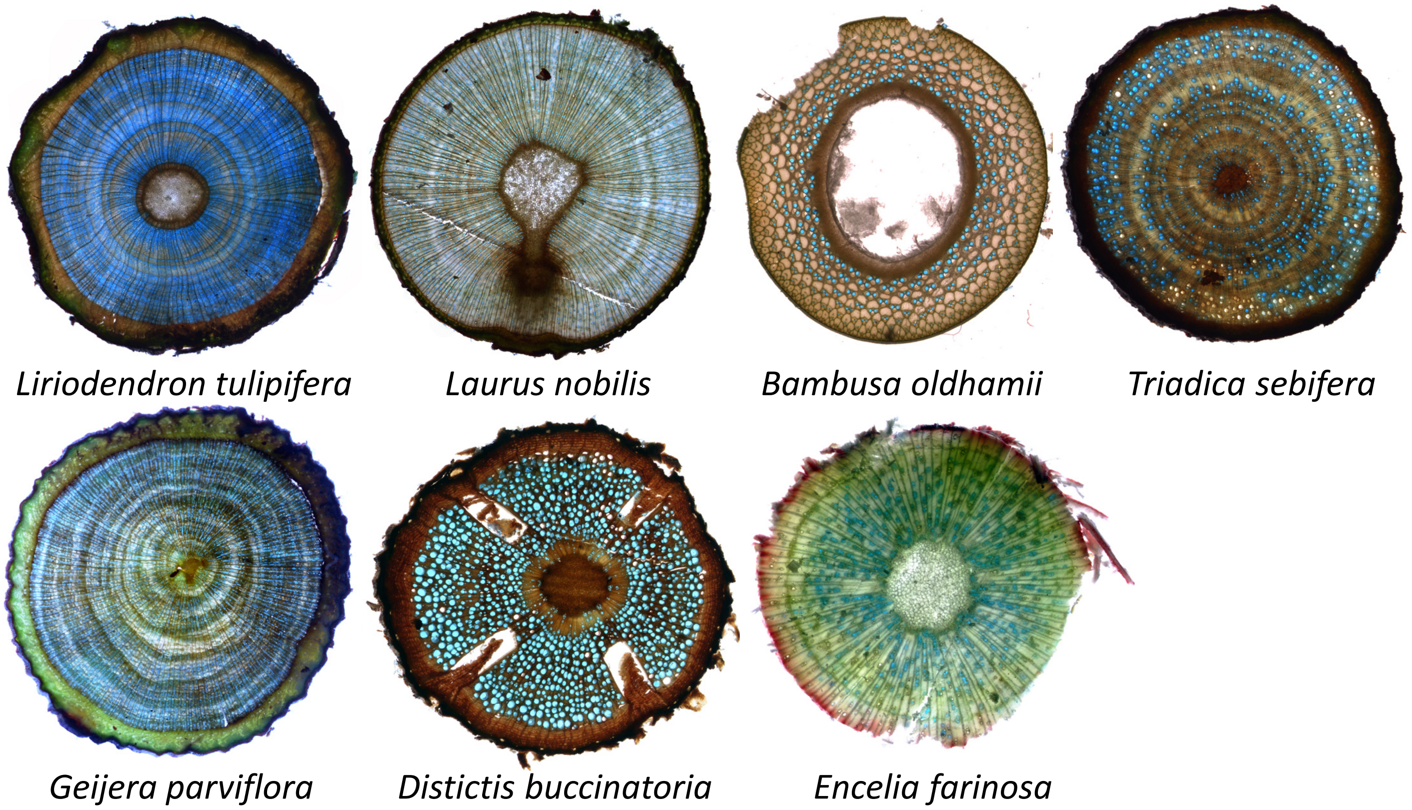
*Liriodendron tulipifera*
*Laurus nobilis*
*Bambusa oldhamii*
*Triadica sebifera*
*Geijera parviflora*
*Distictis buccinatoria*
*Encelia farinosa*

Fig. S10

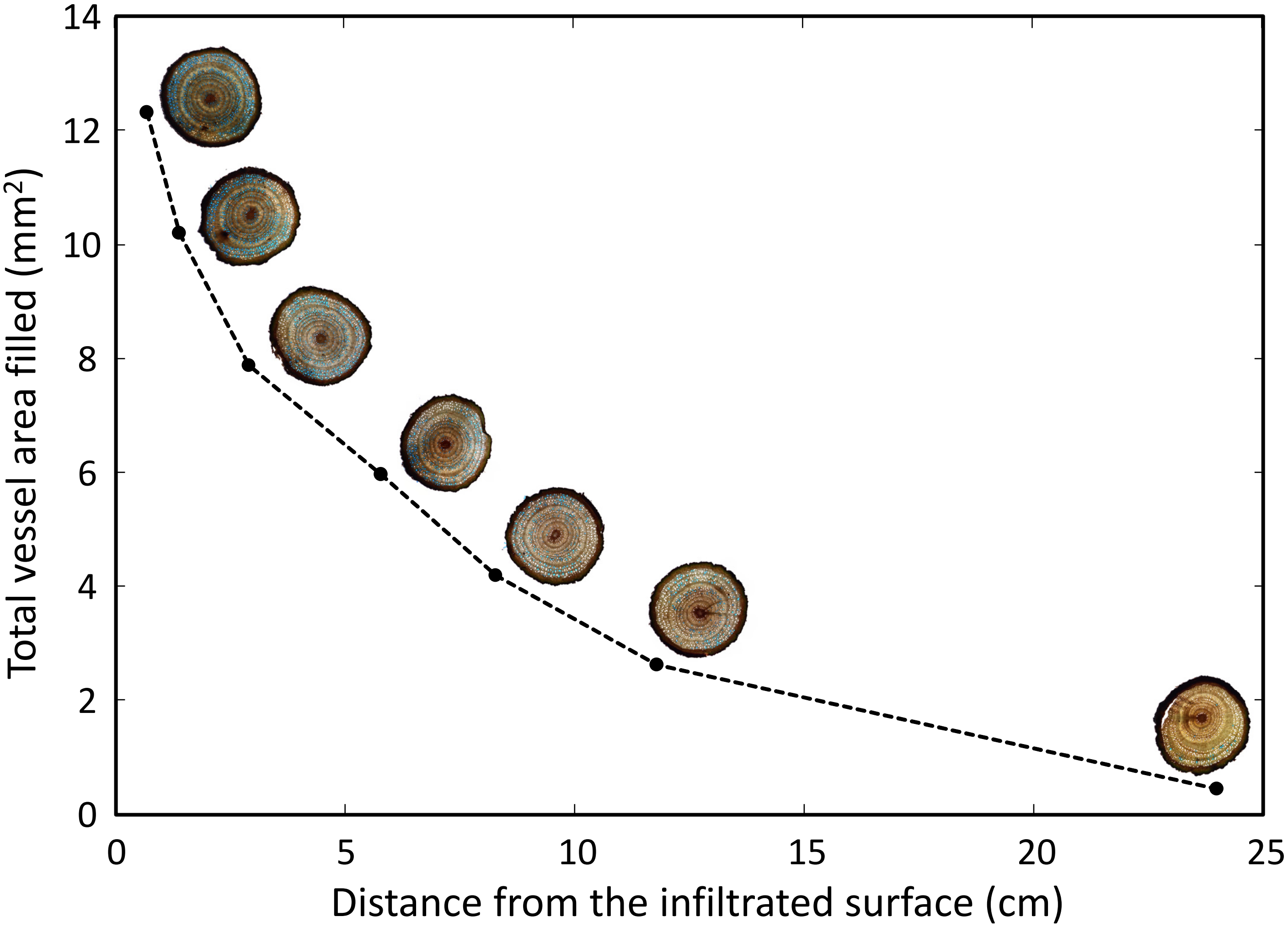

14
12
10
8
6
4
2
0
0
5
10
15
20
25
Total vessel area filled (mm2)
Distance from the infiltrated surface (cm)


Fig. S11

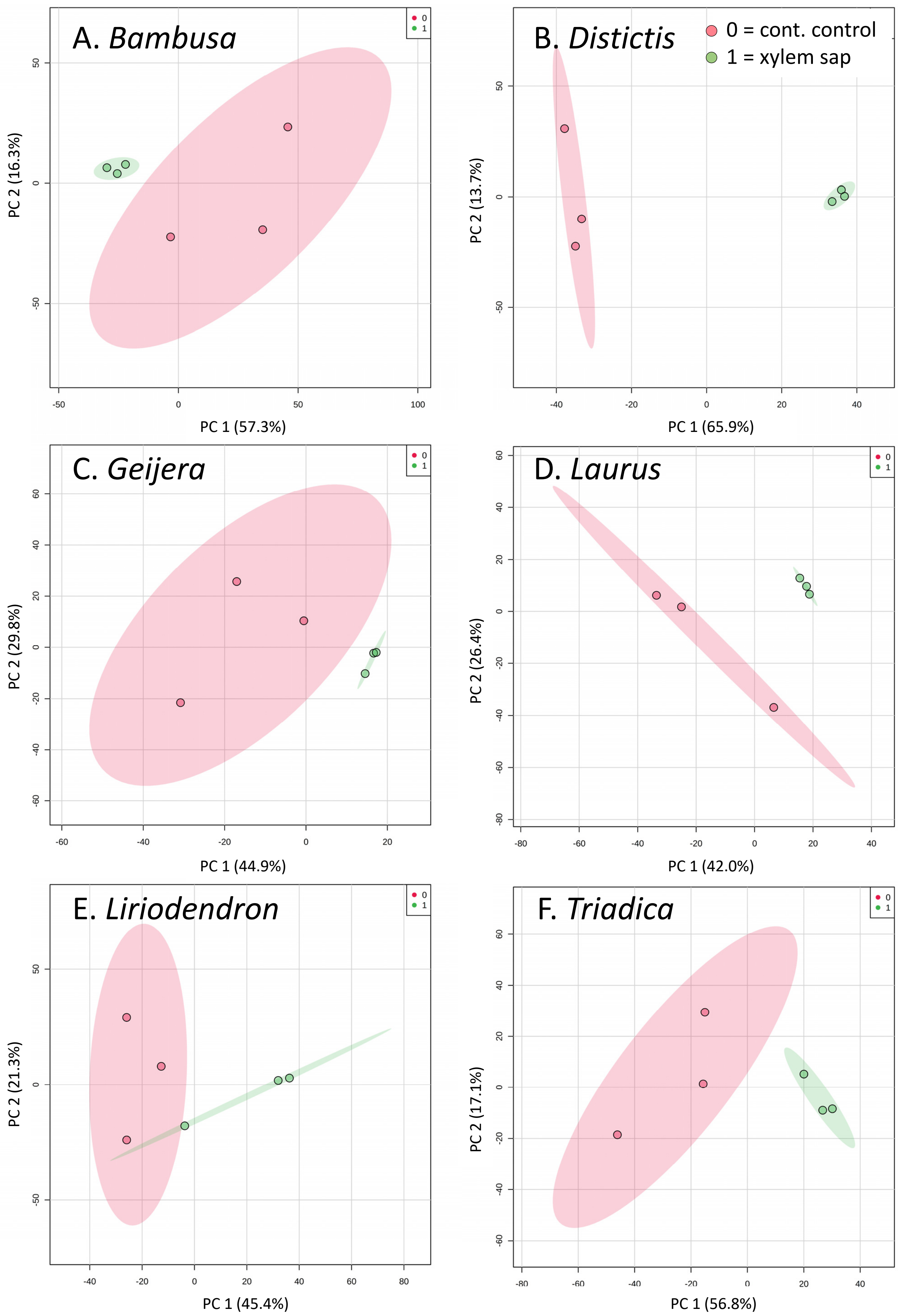

A. Bambusa
B. Distictis
C. Geijera
D. Laurus
E. Liriodendron
F. Triadica
0 = cont. control
1 = xylem sap
0
1
PC 1 (57.3%)
PC 2 (16.3%)
PC 1 (65.9%)
PC 2 (13.7%)
PC 1 (44.9%)
PC 2 (29.8%)
PC 1 (42.0%)
PC 2 (26.4%)
PC 1 (45.4%)
PC 2 (21.3%)
PC 1 (56.8%)
PC 2 (17.1%)


Fig. S12

**Supplementary Information**

**Methods S1: Methods comparison of lipid extraction by partial freeze drying vs. SpeedVac**

*Introduction*

In the course of the research reported in this paper, a SpeedVac evaporator instrument designed for evaporation of chloroform became available in 2018, before stage 2 measurements were conducted, which allowed us to simplify procedures of lipid extraction. To validate this new method against the partial freeze dry method used in earlier measurements, we conducted a methods comparison.

*Materials and Methods*

Xylem sap was extracted from eight *Geijera parviflora* stems (2 mL per stem) as described in the methods chapter. To create paired samples, sap from two stems was combined into a 4 mL sample, mixed, and then divided into two identical samples to create two groups of paired samples, four that were then extracted via the partial freeze dry protocol and four extracted using the SpeedVac. The only difference in protocols to those described in the Methods section was that glass vials were used instead of LoBind Eppendorf tubes for the partial freeze dry protocol.

Samples were analyzed via mass spectrometry for 156 polar lipids and 39 triglycerides (TAGs), as described in the methods chapter. Total concentrations of polar lipids and TAGs were compared between the methods using paired t-tests with equal variance in Microsoft Excel. Lipid composition was compared using PCA and t-tests adjusted for false discovery rate (FDR) in MetaboAnalyst after cube-root transforming the data to normalize distributions.

*Results and Discussion*

Total concentrations of polar lipids were not different between the two methods (Partial freeze dry: 0.207 ± 0.041 SE nmol / mL; SpeedVac: 0.247 ± 0.032 SE nmol / mL, paired t-test: $p$ = 0.220). Lipid composition was not different between the two methods, as their 95% confidence regions in PCA overlapped completely (See Figure), but the SpeedVac method produced higher variability in composition between samples. No lipids were identified as significantly different ($p < 0.05$ after adjustment for FDR) when using paired t-tests.

Lipid extraction using a SpeedVac evaporator is much less labor-intensive and much faster than the partial freeze dry method, which remains a viable choice for research labs that do not have access to a SpeedVac model that is designed for evaporating chloroform.

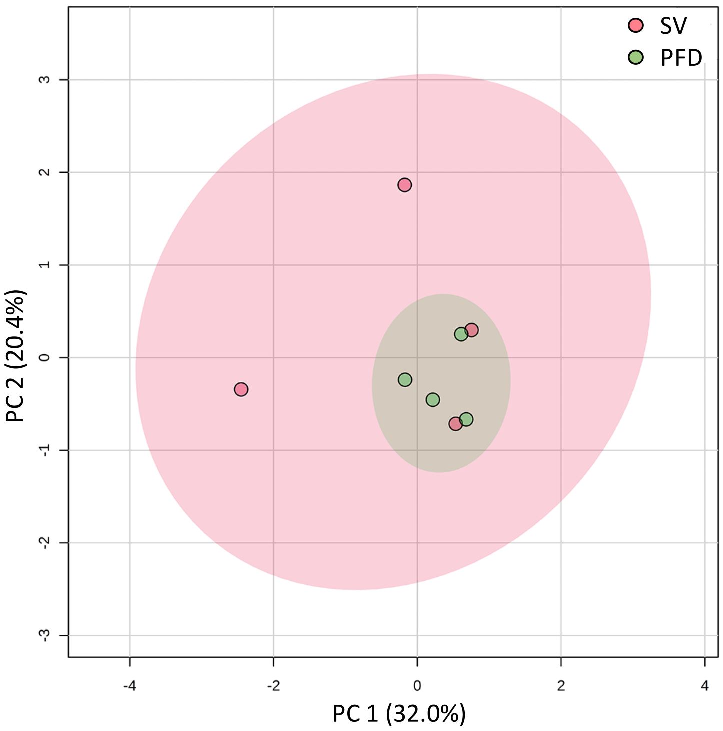


Figure.  Principal component analysis (PCA) of polar lipid composition in xylem sap samples processed by partial free drying (PFD, green) or SpeedVac evaporation (SV, pink). The shaded areas show 95% confidence regions.

Table S1. Standards used for mass spectrometry of xylem sap lipids.

| Internal standard | Amount added (nmol) | Standard for |
|---|---|---|
| | Phospholipids | |
| LPC (13:0) | 0.300 | LPC |
| LPC (19:0) | 0.300 | LPC |
| LPE (14:0) | 0.150 | LPE |
| LPE (18:0) | 0.150 | LPE |
| LPG (14:0) | 0.150 | LPG |
| LPG (18:0) | 0.150 | LPG |
| PA (14:0/14:0) | 0.150 | PA |
| PA (diphytanoyl, i.e., 20:0/20:0) | 0.150 | PA |
| PC (12:0/12:0) | 0.300 | PC |
| PC (24:1/24:1) | 0.300 | PC |
| PE (12:0/12:0) | 0.150 | PE |
| PE (diphytanoyl, i.e., 20:0/20:0) | 0.150 | PE |
| PG (14:0/14:0) | 0.150 | PG |
| PG (diphytanoyl, i.e., 20:0/20:0) | 0.150 | PG |
| PI (16:0/18:0) | 0.144 | PI |
| PI (18:0/18:0) | 0.053 | PI |
| PS (diphytanoyl, i.e., 20:0/20:0) | 0.100 | PS |
| PS14:0/14:0) | 0.100 | PS |
| | Galactolipids | |
| DGDG (18:0/16:0) | 0.220 | DGDG |
| DGDG (18:0/18:0) | 0.740 | DGDG |
| MGDG (18:0/16:0) | 0.833 | MGDG and SQDG |
| MGDG (18:0/18:0) | 0.703 | MGDG |

Table S2. Lipids in xylem sap (mean of n = 4, µmol $L^{-1}$). In parentheses: Number of carbon atoms in acyl chain:number of double bonds.

| | *Laurus nobilis* | | *Liriodendron tulipifera* | | *Bambusa oldhamii* | | *Triadica sebifera* | | *Geijera parviflora* | | *Distictis buccinatoria* | | *Encelia farinosa* | |
|---|---|---|---|---|---|---|---|---|---|---|---|---|---|---|
| **Compound*** | **mean** | **std** | **mean** | **std** | **mean** | **std** | **mean** | **std** | **mean** | **std** | **mean** | **std** | **mean** | **std** |
| DGDG(34:4) | 0.000 | 0.000 | 0.000 | 0.000 | 0.001 | 0.000 | 0.000 | 0.000 | 0.000 | 0.000 | 0.000 | 0.000 | 0.000 | 0.000 |
| DGDG(34:3) | 0.002 | 0.000 | 0.002 | 0.001 | 0.026 | 0.006 | 0.006 | 0.001 | 0.003 | 0.001 | 0.002 | 0.001 | 0.004 | 0.002 |
| DGDG(34:2) | 0.007 | 0.002 | 0.002 | 0.001 | 0.034 | 0.009 | 0.004 | 0.000 | 0.002 | 0.001 | 0.002 | 0.001 | 0.003 | 0.002 |
| DGDG(34:1) | 0.000 | 0.000 | 0.000 | 0.000 | 0.010 | 0.003 | 0.004 | 0.001 | 0.001 | 0.000 | 0.003 | 0.001 | 0.001 | 0.001 |
| DGDG(36:6) | 0.001 | 0.001 | 0.001 | 0.001 | 0.013 | 0.001 | 0.007 | 0.002 | 0.002 | 0.002 | 0.001 | 0.000 | 0.002 | 0.001 |
| DGDG(36:5) | 0.005 | 0.001 | 0.002 | 0.001 | 0.046 | 0.012 | 0.007 | 0.001 | 0.002 | 0.001 | 0.006 | 0.002 | 0.002 | 0.001 |
| DGDG(36:4) | 0.020 | 0.007 | 0.004 | 0.002 | 0.055 | 0.014 | 0.010 | 0.002 | 0.003 | 0.001 | 0.010 | 0.003 | 0.001 | 0.001 |
| DGDG(36:3) | 0.002 | 0.001 | 0.002 | 0.001 | 0.014 | 0.002 | 0.007 | 0.000 | 0.001 | 0.001 | 0.009 | 0.002 | 0.001 | 0.001 |
| DGDG(36:2) | 0.001 | 0.000 | 0.000 | 0.000 | 0.004 | 0.001 | 0.004 | 0.002 | 0.001 | 0.001 | 0.005 | 0.001 | 0.001 | 0.001 |
| DGDG(36:1) | 0.000 | 0.000 | 0.000 | 0.000 | 0.001 | 0.000 | 0.001 | 0.000 | 0.000 | 0.000 | 0.001 | 0.001 | 0.000 | 0.000 |
| DGDG(38:6) | 0.000 | 0.001 | 0.000 | 0.000 | 0.000 | 0.000 | 0.000 | 0.000 | 0.000 | 0.000 | 0.000 | 0.000 | 0.000 | 0.000 |
| DGDG(38:5) | 0.000 | 0.000 | 0.001 | 0.001 | 0.001 | 0.000 | 0.000 | 0.000 | 0.000 | 0.000 | 0.000 | 0.000 | 0.000 | 0.000 |
| DGDG(38:4) | 0.000 | 0.000 | 0.002 | 0.002 | 0.001 | 0.001 | 0.001 | 0.001 | 0.000 | 0.000 | 0.000 | 0.000 | 0.000 | 0.000 |
| **Total DGDG** | **0.038** | **0.013** | **0.015** | **0.010** | **0.203** | **0.046** | **0.051** | **0.009** | **0.015** | **0.006** | **0.041** | **0.010** | **0.015** | **0.008** |
| MGDG(34:6) | 0.000 | 0.000 | 0.001 | 0.001 | 0.000 | 0.000 | 0.000 | 0.000 | 0.001 | 0.001 | 0.000 | 0.000 | 0.000 | 0.000 |
| MGDG(34:5) | 0.000 | 0.000 | 0.001 | 0.001 | 0.000 | 0.000 | 0.001 | 0.001 | 0.000 | 0.000 | 0.000 | 0.000 | 0.000 | 0.000 |
| MGDG(34:4) | 0.000 | 0.000 | 0.000 | 0.000 | 0.000 | 0.000 | 0.000 | 0.000 | 0.000 | 0.000 | 0.000 | 0.000 | 0.001 | 0.000 |
| MGDG(34:3) | 0.001 | 0.001 | 0.001 | 0.000 | 0.004 | 0.001 | 0.001 | 0.001 | 0.002 | 0.001 | 0.001 | 0.000 | 0.002 | 0.001 |
| MGDG(34:2) | 0.003 | 0.002 | 0.001 | 0.001 | 0.002 | 0.001 | 0.001 | 0.000 | 0.001 | 0.000 | 0.000 | 0.000 | 0.001 | 0.001 |
| MGDG(34:1) | 0.000 | 0.000 | 0.000 | 0.000 | 0.000 | 0.000 | 0.000 | 0.000 | 0.000 | 0.000 | 0.000 | 0.000 | 0.000 | 0.000 |
| MGDG(36:6) | 0.006 | 0.003 | 0.005 | 0.004 | 0.031 | 0.000 | 0.014 | 0.006 | 0.007 | 0.005 | 0.007 | 0.002 | 0.007 | 0.003 |
| MGDG(36:5) | 0.021 | 0.011 | 0.018 | 0.017 | 0.067 | 0.011 | 0.011 | 0.002 | 0.008 | 0.003 | 0.021 | 0.007 | 0.005 | 0.003 |
| MGDG(36:4) | 0.063 | 0.032 | 0.006 | 0.004 | 0.050 | 0.010 | 0.009 | 0.001 | 0.007 | 0.003 | 0.017 | 0.006 | 0.003 | 0.002 |
| MGDG(36:3) | 0.001 | 0.000 | 0.001 | 0.001 | 0.003 | 0.002 | 0.003 | 0.000 | 0.002 | 0.001 | 0.003 | 0.001 | 0.001 | 0.001 |
| MGDG(36:2) | 0.000 | 0.000 | 0.000 | 0.000 | 0.000 | 0.000 | 0.001 | 0.001 | 0.000 | 0.000 | 0.001 | 0.000 | 0.000 | 0.000 |
| MGDG(38:6) | 0.001 | 0.001 | 0.000 | 0.000 | 0.000 | 0.000 | 0.000 | 0.000 | 0.000 | 0.000 | 0.000 | 0.000 | 0.000 | 0.000 |
| MGDG(38:5) | 0.001 | 0.001 | 0.003 | 0.002 | 0.000 | 0.000 | 0.000 | 0.000 | 0.002 | 0.001 | 0.000 | 0.000 | 0.000 | 0.000 |
| MGDG(38:4) | 0.000 | 0.000 | 0.011 | 0.010 | 0.000 | 0.000 | 0.000 | 0.000 | 0.003 | 0.003 | 0.000 | 0.000 | 0.000 | 0.000 |
| MGDG(38:3) | 0.000 | 0.000 | 0.001 | 0.001 | 0.000 | 0.000 | 0.000 | 0.000 | 0.000 | 0.000 | 0.000 | 0.000 | 0.000 | 0.001 |
| **Total MGDG** | **0.099** | **0.052** | **0.050** | **0.041** | **0.158** | **0.022** | **0.042** | **0.007** | **0.032** | **0.012** | **0.050** | **0.016** | **0.021** | **0.010** |

| Compound* | *Laurus nobilis* mean | std | *Liriodendron tulipifera* mean | std | *Bambusa oldhamii* mean | std | *Triadica sebifera* mean | std | *Geijera parviflora* mean | std | *Distictis buccinatoria* mean | std | *Encelia farinosa* mean | std |
|---|---|---|---|---|---|---|---|---|---|---|---|---|---|---|
| PG(32:0) | 0.000 | 0.000 | 0.001 | 0.001 | 0.000 | 0.000 | 0.003 | 0.001 | 0.003 | 0.003 | 0.001 | 0.001 | 0.002 | 0.002 |
| PG(34:4) | 0.000 | 0.000 | 0.000 | 0.000 | 0.000 | 0.000 | 0.000 | 0.000 | 0.000 | 0.000 | 0.000 | 0.000 | 0.000 | 0.000 |
| PG(34:3) | 0.000 | 0.000 | 0.000 | 0.000 | 0.000 | 0.000 | 0.001 | 0.001 | 0.000 | 0.000 | 0.000 | 0.000 | 0.000 | 0.000 |
| PG(34:2) | 0.000 | 0.000 | 0.001 | 0.001 | 0.001 | 0.000 | 0.001 | 0.000 | 0.000 | 0.000 | 0.000 | 0.000 | 0.001 | 0.001 |
| PG(34:1) | 0.000 | 0.000 | 0.000 | 0.000 | 0.002 | 0.001 | 0.002 | 0.000 | 0.000 | 0.000 | 0.001 | 0.000 | 0.001 | 0.001 |
| **Total PG** | **0.001** | **0.000** | **0.002** | **0.002** | **0.003** | **0.001** | **0.007** | **0.002** | **0.003** | **0.003** | **0.003** | **0.001** | **0.004** | **0.004** |
| LPG(16:1) | 0.000 | 0.000 | 0.000 | 0.000 | 0.002 | 0.003 | 0.002 | 0.001 | 0.003 | 0.003 | 0.000 | 0.000 | 0.001 | 0.001 |
| LPG(16:0) | 0.008 | 0.002 | 0.006 | 0.003 | 0.012 | 0.004 | 0.006 | 0.002 | 0.004 | 0.003 | 0.004 | 0.001 | 0.006 | 0.001 |
| LPG(18:3) | 0.000 | 0.000 | 0.000 | 0.000 | 0.000 | 0.000 | 0.000 | 0.000 | 0.000 | 0.000 | 0.000 | 0.000 | 0.001 | 0.001 |
| LPG(18:2) | 0.001 | 0.000 | 0.001 | 0.001 | 0.004 | 0.003 | 0.000 | 0.000 | 0.001 | 0.000 | 0.001 | 0.001 | 0.001 | 0.000 |
| LPG(18:1) | 0.001 | 0.000 | 0.001 | 0.001 | 0.005 | 0.007 | 0.001 | 0.001 | 0.001 | 0.000 | 0.001 | 0.001 | 0.001 | 0.001 |
| **Total LysoPG** | **0.010** | **0.002** | **0.008** | **0.003** | **0.024** | **0.013** | **0.010** | **0.004** | **0.008** | **0.006** | **0.005** | **0.003** | **0.010** | **0.003** |
| LPC(16:0) | 0.000 | 0.000 | 0.001 | 0.001 | 0.000 | 0.000 | 0.001 | 0.001 | 0.001 | 0.001 | 0.001 | 0.000 | 0.003 | 0.002 |
| LPC(18:3) | 0.000 | 0.000 | 0.000 | 0.000 | 0.002 | 0.001 | 0.000 | 0.000 | 0.000 | 0.000 | 0.000 | 0.000 | 0.000 | 0.000 |
| LPC(18:2) | 0.000 | 0.000 | 0.001 | 0.001 | 0.005 | 0.002 | 0.001 | 0.001 | 0.000 | 0.000 | 0.000 | 0.000 | 0.002 | 0.001 |
| LPC(18:1) | 0.000 | 0.000 | 0.000 | 0.000 | 0.000 | 0.000 | 0.001 | 0.001 | 0.000 | 0.000 | 0.000 | 0.000 | 0.002 | 0.001 |
| **Total LysoPC** | **0.000** | **0.000** | **0.003** | **0.003** | **0.006** | **0.003** | **0.004** | **0.002** | **0.001** | **0.001** | **0.001** | **0.001** | **0.007** | **0.004** |
| LPE(16:0) | 0.001 | 0.001 | 0.000 | 0.000 | 0.001 | 0.000 | 0.001 | 0.001 | 0.001 | 0.001 | 0.001 | 0.000 | 0.003 | 0.003 |
| LPE(18:3) | 0.000 | 0.000 | 0.000 | 0.000 | 0.000 | 0.000 | 0.000 | 0.000 | 0.000 | 0.000 | 0.000 | 0.000 | 0.000 | 0.000 |
| LPE(18:2) | 0.000 | 0.000 | 0.000 | 0.000 | 0.002 | 0.002 | 0.000 | 0.001 | 0.000 | 0.000 | 0.000 | 0.000 | 0.001 | 0.001 |
| LPE(18:1) | 0.001 | 0.001 | 0.000 | 0.000 | 0.000 | 0.000 | 0.000 | 0.000 | 0.000 | 0.000 | 0.000 | 0.000 | 0.001 | 0.001 |
| **Total LysoPE** | **0.002** | **0.002** | **0.001** | **0.001** | **0.003** | **0.002** | **0.002** | **0.002** | **0.001** | **0.001** | **0.001** | **0.001** | **0.005** | **0.005** |
| PC(32:0) | 0.000 | 0.000 | 0.000 | 0.000 | 0.000 | 0.000 | 0.001 | 0.001 | 0.001 | 0.001 | 0.000 | 0.000 | 0.001 | 0.000 |
| PC(34:4) | 0.000 | 0.000 | 0.000 | 0.000 | 0.000 | 0.000 | 0.000 | 0.000 | 0.000 | 0.000 | 0.000 | 0.000 | 0.000 | 0.000 |
| PC(34:3) | 0.001 | 0.000 | 0.001 | 0.001 | 0.010 | 0.005 | 0.022 | 0.011 | 0.007 | 0.003 | 0.007 | 0.001 | 0.006 | 0.004 |
| PC(34:2) | 0.010 | 0.002 | 0.015 | 0.007 | 0.041 | 0.021 | 0.026 | 0.014 | 0.028 | 0.010 | 0.012 | 0.003 | 0.028 | 0.023 |
| PC(34:1) | 0.000 | 0.000 | 0.004 | 0.002 | 0.006 | 0.005 | 0.030 | 0.001 | 0.009 | 0.006 | 0.013 | 0.005 | 0.012 | 0.009 |
| PC(36:6) | 0.000 | 0.000 | 0.000 | 0.000 | 0.001 | 0.001 | 0.005 | 0.002 | 0.000 | 0.000 | 0.002 | 0.000 | 0.001 | 0.000 |
| PC(36:5) | 0.001 | 0.000 | 0.001 | 0.000 | 0.006 | 0.003 | 0.007 | 0.004 | 0.003 | 0.002 | 0.006 | 0.002 | 0.003 | 0.002 |
| PC(36:4) | 0.008 | 0.001 | 0.010 | 0.004 | 0.014 | 0.007 | 0.024 | 0.005 | 0.011 | 0.005 | 0.019 | 0.010 | 0.009 | 0.006 |
| PC(36:3) | 0.001 | 0.000 | 0.006 | 0.002 | 0.007 | 0.004 | 0.016 | 0.005 | 0.006 | 0.003 | 0.017 | 0.007 | 0.006 | 0.004 |

| Compound* | *Laurus nobilis* mean | std | *Liriodendron tulipifera* mean | std | *Bambusa oldhamii* mean | std | *Triadica sebifera* mean | std | *Geijera parviflora* mean | std | *Distictis buccinatoria* mean | std | *Encelia farinosa* mean | std |
|---|---|---|---|---|---|---|---|---|---|---|---|---|---|---|
| PC(36:2) | 0.000 | 0.000 | 0.002 | 0.001 | 0.002 | 0.001 | 0.012 | 0.002 | 0.003 | 0.002 | 0.008 | 0.003 | 0.004 | 0.003 |
| PC(36:1) | 0.000 | 0.000 | 0.000 | 0.000 | 0.000 | 0.000 | 0.004 | 0.001 | 0.001 | 0.001 | 0.003 | 0.001 | 0.001 | 0.001 |
| PC(38:6) | 0.000 | 0.000 | 0.000 | 0.000 | 0.000 | 0.000 | 0.000 | 0.000 | 0.000 | 0.000 | 0.000 | 0.000 | 0.000 | 0.000 |
| PC(38:5) | 0.000 | 0.000 | 0.000 | 0.000 | 0.000 | 0.000 | 0.000 | 0.000 | 0.000 | 0.000 | 0.000 | 0.000 | 0.000 | 0.000 |
| PC(38:4) | 0.000 | 0.000 | 0.000 | 0.000 | 0.000 | 0.000 | 0.000 | 0.000 | 0.000 | 0.000 | 0.000 | 0.000 | 0.000 | 0.000 |
| PC(38:3) | 0.000 | 0.000 | 0.000 | 0.000 | 0.000 | 0.000 | 0.000 | 0.000 | 0.000 | 0.000 | 0.001 | 0.000 | 0.000 | 0.000 |
| PC(38:2) | 0.000 | 0.000 | 0.000 | 0.000 | 0.000 | 0.000 | 0.001 | 0.000 | 0.000 | 0.000 | 0.001 | 0.000 | 0.000 | 0.000 |
| **Total PC** | **0.022** | **0.003** | **0.040** | **0.017** | **0.088** | **0.046** | **0.148** | **0.042** | **0.070** | **0.031** | **0.087** | **0.032** | **0.071** | **0.053** |
| PE(32:1) | 0.000 | 0.000 | 0.000 | 0.000 | 0.000 | 0.000 | 0.000 | 0.000 | 0.000 | 0.000 | 0.000 | 0.000 | 0.000 | 0.001 |
| PE(32:0) | 0.000 | 0.000 | 0.000 | 0.000 | 0.000 | 0.000 | 0.000 | 0.000 | 0.000 | 0.000 | 0.000 | 0.000 | 0.000 | 0.000 |
| PE(34:4) | 0.000 | 0.000 | 0.000 | 0.000 | 0.000 | 0.000 | 0.000 | 0.000 | 0.000 | 0.000 | 0.000 | 0.000 | 0.000 | 0.000 |
| PE(34:3) | 0.000 | 0.000 | 0.000 | 0.000 | 0.003 | 0.002 | 0.008 | 0.008 | 0.005 | 0.001 | 0.004 | 0.001 | 0.004 | 0.004 |
| PE(34:2) | 0.003 | 0.000 | 0.008 | 0.006 | 0.013 | 0.010 | 0.012 | 0.012 | 0.030 | 0.010 | 0.012 | 0.005 | 0.024 | 0.028 |
| PE(34:1) | 0.000 | 0.000 | 0.001 | 0.001 | 0.000 | 0.000 | 0.006 | 0.005 | 0.003 | 0.002 | 0.005 | 0.002 | 0.004 | 0.006 |
| PE(36:6) | 0.000 | 0.000 | 0.000 | 0.000 | 0.000 | 0.000 | 0.002 | 0.002 | 0.000 | 0.000 | 0.001 | 0.000 | 0.000 | 0.000 |
| PE(36:5) | 0.000 | 0.000 | 0.000 | 0.000 | 0.001 | 0.001 | 0.002 | 0.002 | 0.001 | 0.000 | 0.003 | 0.001 | 0.001 | 0.002 |
| PE(36:4) | 0.003 | 0.001 | 0.003 | 0.002 | 0.006 | 0.005 | 0.005 | 0.005 | 0.007 | 0.003 | 0.009 | 0.004 | 0.005 | 0.006 |
| PE(36:3) | 0.000 | 0.000 | 0.001 | 0.001 | 0.001 | 0.001 | 0.003 | 0.003 | 0.002 | 0.001 | 0.007 | 0.003 | 0.002 | 0.003 |
| PE(36:2) | 0.000 | 0.000 | 0.001 | 0.001 | 0.000 | 0.000 | 0.002 | 0.002 | 0.001 | 0.001 | 0.004 | 0.002 | 0.002 | 0.003 |
| PE(36:1) | 0.000 | 0.000 | 0.000 | 0.000 | 0.000 | 0.000 | 0.001 | 0.001 | 0.000 | 0.000 | 0.001 | 0.000 | 0.000 | 0.000 |
| PE(42:2) | 0.000 | 0.000 | 0.000 | 0.000 | 0.000 | 0.000 | 0.000 | 0.000 | 0.001 | 0.000 | 0.000 | 0.000 | 0.001 | 0.001 |
| **Total PE** | **0.007** | **0.001** | **0.016** | **0.014** | **0.026** | **0.020** | **0.042** | **0.042** | **0.051** | **0.016** | **0.046** | **0.018** | **0.046** | **0.055** |
| PI(32:2) | 0.000 | 0.000 | 0.000 | 0.000 | 0.000 | 0.000 | 0.000 | 0.000 | 0.001 | 0.001 | 0.000 | 0.000 | 0.000 | 0.000 |
| PI(32:1) | 0.000 | 0.000 | 0.000 | 0.000 | 0.000 | 0.000 | 0.000 | 0.000 | 0.003 | 0.003 | 0.000 | 0.000 | 0.002 | 0.003 |
| PI(32:0) | 0.000 | 0.000 | 0.001 | 0.001 | 0.001 | 0.000 | 0.001 | 0.000 | 0.001 | 0.001 | 0.000 | 0.000 | 0.001 | 0.001 |
| PI(34:4) | 0.000 | 0.000 | 0.000 | 0.000 | 0.000 | 0.000 | 0.000 | 0.000 | 0.000 | 0.000 | 0.000 | 0.000 | 0.000 | 0.000 |
| PI(34:3) | 0.001 | 0.000 | 0.000 | 0.000 | 0.007 | 0.003 | 0.022 | 0.006 | 0.007 | 0.006 | 0.018 | 0.009 | 0.008 | 0.009 |
| PI(34:2) | 0.005 | 0.002 | 0.006 | 0.007 | 0.016 | 0.005 | 0.026 | 0.008 | 0.021 | 0.014 | 0.019 | 0.012 | 0.048 | 0.058 |
| PI(34:1) | 0.000 | 0.000 | 0.001 | 0.001 | 0.001 | 0.000 | 0.011 | 0.002 | 0.002 | 0.002 | 0.009 | 0.005 | 0.004 | 0.004 |
| PI(36:6) | 0.000 | 0.000 | 0.000 | 0.000 | 0.000 | 0.000 | 0.000 | 0.000 | 0.000 | 0.000 | 0.000 | 0.000 | 0.001 | 0.000 |
| PI(36:5) | 0.000 | 0.000 | 0.000 | 0.000 | 0.000 | 0.000 | 0.000 | 0.000 | 0.000 | 0.000 | 0.001 | 0.000 | 0.001 | 0.001 |
| PI(36:4) | 0.000 | 0.000 | 0.000 | 0.000 | 0.001 | 0.001 | 0.002 | 0.000 | 0.001 | 0.000 | 0.002 | 0.001 | 0.002 | 0.002 |

| Compound* | *Laurus nobilis* mean | std | *Liriodendron tulipifera* mean | std | *Bambusa oldhamii* mean | std | *Triadica sebifera* mean | std | *Geijera parviflora* mean | std | *Distictis buccinatoria* mean | std | *Encelia farinosa* mean | std |
|---|---|---|---|---|---|---|---|---|---|---|---|---|---|---|
| PI(36:3) | 0.000 | 0.000 | 0.000 | 0.000 | 0.001 | 0.000 | 0.002 | 0.001 | 0.000 | 0.000 | 0.001 | 0.001 | 0.001 | 0.001 |
| PI(36:2) | 0.000 | 0.000 | 0.000 | 0.000 | 0.000 | 0.000 | 0.002 | 0.000 | 0.001 | 0.001 | 0.001 | 0.001 | 0.002 | 0.002 |
| PI(36:1) | 0.000 | 0.000 | 0.000 | 0.000 | 0.000 | 0.000 | 0.001 | 0.000 | 0.000 | 0.000 | 0.000 | 0.000 | 0.000 | 0.000 |
| **Total PI** | **0.007** | **0.003** | **0.009** | **0.009** | **0.027** | **0.009** | **0.068** | **0.018** | **0.035** | **0.027** | **0.051** | **0.030** | **0.071** | **0.081** |
| PS(34:3) | 0.000 | 0.000 | 0.000 | 0.000 | 0.000 | 0.000 | 0.001 | 0.001 | 0.000 | 0.000 | 0.000 | 0.000 | 0.000 | 0.001 |
| PS(34:2) | 0.000 | 0.000 | 0.001 | 0.001 | 0.001 | 0.000 | 0.002 | 0.001 | 0.002 | 0.001 | 0.002 | 0.002 | 0.002 | 0.003 |
| PS(34:1) | 0.000 | 0.000 | 0.000 | 0.000 | 0.000 | 0.000 | 0.001 | 0.000 | 0.000 | 0.000 | 0.001 | 0.001 | 0.000 | 0.000 |
| PS(36:6) | 0.000 | 0.000 | 0.000 | 0.000 | 0.000 | 0.000 | 0.000 | 0.000 | 0.000 | 0.000 | 0.000 | 0.000 | 0.000 | 0.000 |
| PS(36:5) | 0.000 | 0.000 | 0.000 | 0.000 | 0.000 | 0.000 | 0.000 | 0.000 | 0.000 | 0.000 | 0.000 | 0.000 | 0.000 | 0.000 |
| PS(36:4) | 0.000 | 0.000 | 0.000 | 0.000 | 0.000 | 0.000 | 0.001 | 0.000 | 0.000 | 0.000 | 0.001 | 0.001 | 0.000 | 0.001 |
| PS(36:3) | 0.000 | 0.000 | 0.000 | 0.000 | 0.000 | 0.000 | 0.000 | 0.000 | 0.000 | 0.000 | 0.001 | 0.001 | 0.000 | 0.000 |
| PS(36:2) | 0.000 | 0.000 | 0.000 | 0.000 | 0.000 | 0.000 | 0.000 | 0.000 | 0.000 | 0.000 | 0.001 | 0.001 | 0.000 | 0.000 |
| PS(38:3) | 0.000 | 0.000 | 0.000 | 0.000 | 0.000 | 0.000 | 0.000 | 0.000 | 0.000 | 0.000 | 0.001 | 0.000 | 0.001 | 0.001 |
| PS(38:2) | 0.000 | 0.000 | 0.000 | 0.000 | 0.001 | 0.000 | 0.000 | 0.000 | 0.001 | 0.001 | 0.002 | 0.001 | 0.002 | 0.002 |
| PS(38:1) | 0.000 | 0.000 | 0.000 | 0.000 | 0.000 | 0.000 | 0.000 | 0.000 | 0.000 | 0.000 | 0.002 | 0.001 | 0.000 | 0.000 |
| PS(40:3) | 0.000 | 0.000 | 0.000 | 0.000 | 0.000 | 0.000 | 0.001 | 0.000 | 0.000 | 0.000 | 0.001 | 0.000 | 0.001 | 0.001 |
| PS(40:2) | 0.000 | 0.000 | 0.001 | 0.001 | 0.001 | 0.001 | 0.001 | 0.001 | 0.002 | 0.001 | 0.001 | 0.001 | 0.002 | 0.002 |
| PS(40:1) | 0.000 | 0.000 | 0.000 | 0.000 | 0.000 | 0.000 | 0.000 | 0.000 | 0.000 | 0.000 | 0.001 | 0.001 | 0.000 | 0.000 |
| PS(42:4) | 0.000 | 0.000 | 0.000 | 0.000 | 0.000 | 0.000 | 0.000 | 0.000 | 0.000 | 0.000 | 0.000 | 0.000 | 0.000 | 0.000 |
| PS(42:3) | 0.000 | 0.000 | 0.000 | 0.000 | 0.001 | 0.001 | 0.002 | 0.002 | 0.002 | 0.000 | 0.001 | 0.001 | 0.002 | 0.000 |
| PS(42:2) | 0.000 | 0.000 | 0.001 | 0.001 | 0.002 | 0.001 | 0.001 | 0.001 | 0.007 | 0.003 | 0.001 | 0.000 | 0.003 | 0.003 |
| PS(44:3) | 0.000 | 0.000 | 0.001 | 0.001 | 0.000 | 0.000 | 0.000 | 0.000 | 0.000 | 0.000 | 0.000 | 0.000 | 0.000 | 0.000 |
| **Total PS** | **0.001** | **0.000** | **0.004** | **0.003** | **0.007** | **0.003** | **0.011** | **0.006** | **0.016** | **0.007** | **0.013** | **0.008** | **0.015** | **0.015** |
| PA(32:0) | 0.000 | 0.000 | 0.000 | 0.000 | 0.001 | 0.001 | 0.001 | 0.001 | 0.003 | 0.002 | 0.001 | 0.001 | 0.002 | 0.001 |
| PA(34:4) | 0.000 | 0.000 | 0.000 | 0.000 | 0.000 | 0.000 | 0.000 | 0.000 | 0.000 | 0.000 | 0.000 | 0.000 | 0.001 | 0.000 |
| PA(34:3) | 0.000 | 0.000 | 0.002 | 0.001 | 0.014 | 0.006 | 0.023 | 0.012 | 0.029 | 0.017 | 0.018 | 0.010 | 0.029 | 0.014 |
| PA(34:2) | 0.002 | 0.001 | 0.021 | 0.016 | 0.045 | 0.020 | 0.024 | 0.014 | 0.115 | 0.065 | 0.029 | 0.015 | 0.113 | 0.059 |
| PA(34:1) | 0.000 | 0.000 | 0.004 | 0.003 | 0.009 | 0.004 | 0.023 | 0.003 | 0.020 | 0.011 | 0.022 | 0.012 | 0.029 | 0.019 |
| PA(36:6) | 0.000 | 0.000 | 0.000 | 0.000 | 0.000 | 0.000 | 0.004 | 0.003 | 0.001 | 0.000 | 0.002 | 0.001 | 0.003 | 0.002 |
| PA(36:5) | 0.000 | 0.000 | 0.001 | 0.001 | 0.008 | 0.003 | 0.007 | 0.004 | 0.009 | 0.006 | 0.011 | 0.006 | 0.015 | 0.008 |
| PA(36:4) | 0.001 | 0.001 | 0.007 | 0.005 | 0.024 | 0.011 | 0.014 | 0.003 | 0.026 | 0.014 | 0.025 | 0.016 | 0.033 | 0.019 |
| PA(36:3) | 0.000 | 0.000 | 0.004 | 0.002 | 0.007 | 0.002 | 0.011 | 0.004 | 0.012 | 0.005 | 0.027 | 0.018 | 0.021 | 0.014 |

| Compound* | *Laurus nobilis* mean | std | *Liriodendron tulipifera* mean | std | *Bambusa oldhamii* mean | std | *Triadica sebifera* mean | std | *Geijera parviflora* mean | std | *Distictis buccinatoria* mean | std | *Encelia farinosa* mean | std |
|---|---|---|---|---|---|---|---|---|---|---|---|---|---|---|
| PA(36:2) | 0.000 | 0.000 | 0.002 | 0.001 | 0.003 | 0.002 | 0.012 | 0.002 | 0.008 | 0.004 | 0.017 | 0.010 | 0.016 | 0.010 |
| **Total PA** | **0.004** | **0.002** | **0.041** | **0.030** | **0.112** | **0.049** | **0.120** | **0.047** | **0.223** | **0.125** | **0.153** | **0.090** | **0.262** | **0.144** |
| SQDG(34:3) | n.d. | | 0.000 | 0.000 | n.d. | | 0.001 | 0.001 | 0.000 | 0.000 | 0.001 | 0.000 | 0.001 | 0.000 |
| SQDG(34:2) | n.d. | | 0.000 | 0.000 | n.d. | | 0.001 | 0.000 | 0.001 | 0.002 | 0.001 | 0.000 | 0.004 | 0.002 |
| SQDG(34:1) | n.d. | | 0.000 | 0.000 | n.d. | | 0.001 | 0.000 | 0.000 | 0.000 | 0.001 | 0.000 | 0.001 | 0.000 |
| SQDG(36:5) | n.d. | | 0.000 | 0.000 | n.d. | | 0.000 | 0.000 | 0.000 | 0.000 | 0.000 | 0.000 | 0.001 | 0.001 |
| SQDG(36:4) | n.d. | | 0.000 | 0.000 | n.d. | | 0.000 | 0.000 | 0.000 | 0.000 | 0.000 | 0.000 | 0.001 | 0.000 |
| **Total SQDG** | n.d. | | **0.001** | **0.001** | n.d. | | **0.003** | **0.000** | **0.003** | **0.001** | **0.003** | **0.001** | **0.009** | **0.003** |

* DGDG = digalactosyldiacylglycerol; LPC = lysophosphatidylcholine; LPE = lysophosphatidylethanolamine; LPG = lysophosphatidylglycerol; MGDG = monogalactosyldiacylglycerol; PA = phosphatidic acid; PC = phosphatidylcholine; PE = phosphatidylethanolamine; PG = phosphatidylglycerol; PI = phosphatidylinositol; PS = phosphatidylserine

Table S3. Lipids in cell contamination controls for xylem sap (mean of n = 4, µmol $L^{-1}$). In parentheses: Number of carbon atoms in acyl chain:number of double bonds.

| | *Laurus nobilis* | | *Liriodendron tulipifera* | | *Bambusa oldhamii* | | *Triadica sebifera* | | *Geijera parviflora* | | *Distictis buccinatoria* | | *Encelia farinosa* | |
|---|---|---|---|---|---|---|---|---|---|---|---|---|---|---|
| **Compound*** | **mean** | **std** | **mean** | **std** | **mean** | **std** | **mean** | **std** | **mean** | **std** | **mean** | **std** | **mean** | **std** |
| DGDG((34:4) | 0.000 | 0.000 | 0.000 | 0.000 | 0.000 | 0.000 | 0.000 | 0.000 | 0.000 | 0.000 | 0.000 | 0.000 | 0.000 | 0.000 |
| DGDG((34:3) | 0.000 | 0.000 | 0.000 | 0.000 | 0.001 | 0.001 | 0.000 | 0.000 | 0.000 | 0.000 | 0.000 | 0.000 | 0.000 | 0.000 |
| DGDG((34:2) | 0.000 | 0.000 | 0.000 | 0.000 | 0.001 | 0.001 | 0.000 | 0.000 | 0.000 | 0.000 | 0.000 | 0.000 | 0.000 | 0.000 |
| DGDG((34:1) | 0.000 | 0.000 | 0.000 | 0.000 | 0.000 | 0.000 | 0.000 | 0.000 | 0.000 | 0.000 | 0.000 | 0.000 | 0.000 | 0.000 |
| DGDG((36:6) | 0.000 | 0.000 | 0.000 | 0.000 | 0.000 | 0.000 | 0.000 | 0.000 | 0.000 | 0.000 | 0.000 | 0.000 | 0.000 | 0.000 |
| DGDG((36:5) | 0.001 | 0.000 | 0.000 | 0.001 | 0.001 | 0.001 | 0.000 | 0.000 | 0.000 | 0.000 | 0.000 | 0.000 | 0.000 | 0.000 |
| DGDG((36:4) | 0.001 | 0.000 | 0.001 | 0.001 | 0.001 | 0.001 | 0.000 | 0.000 | 0.000 | 0.000 | 0.000 | 0.000 | 0.000 | 0.000 |
| DGDG((36:3) | 0.000 | 0.000 | 0.001 | 0.002 | 0.000 | 0.000 | 0.000 | 0.000 | 0.000 | 0.000 | 0.000 | 0.000 | 0.000 | 0.000 |
| DGDG((36:2) | 0.000 | 0.000 | 0.000 | 0.000 | 0.000 | 0.000 | 0.000 | 0.000 | 0.000 | 0.000 | 0.000 | 0.000 | 0.000 | 0.000 |
| DGDG((36:1) | 0.000 | 0.000 | 0.000 | 0.000 | 0.000 | 0.000 | 0.000 | 0.000 | 0.000 | 0.000 | 0.000 | 0.000 | 0.000 | 0.000 |
| DGDG((38:6) | 0.000 | 0.000 | 0.000 | 0.000 | 0.000 | 0.000 | 0.000 | 0.000 | 0.000 | 0.000 | 0.000 | 0.000 | 0.000 | 0.000 |
| DGDG((38:5) | 0.000 | 0.000 | 0.000 | 0.000 | 0.000 | 0.000 | 0.000 | 0.000 | 0.000 | 0.000 | 0.000 | 0.000 | 0.000 | 0.000 |
| DGDG((38:4) | 0.000 | 0.000 | 0.001 | 0.001 | 0.000 | 0.000 | 0.000 | 0.000 | 0.000 | 0.000 | 0.000 | 0.000 | 0.000 | 0.000 |
| **Total DGDG** | **0.003** | **0.001** | **0.005** | **0.007** | **0.005** | **0.004** | **0.001** | **0.001** | **0.001** | **0.002** | **0.002** | **0.001** | **0.000** | **0.000** |
| MGDG((34:6) | 0.000 | 0.000 | 0.000 | 0.000 | 0.000 | 0.000 | 0.000 | 0.000 | 0.000 | 0.000 | 0.000 | 0.000 | 0.000 | 0.000 |
| MGDG((34:5) | 0.000 | 0.000 | 0.002 | 0.003 | 0.000 | 0.000 | 0.000 | 0.000 | 0.000 | 0.000 | 0.000 | 0.000 | 0.000 | 0.000 |
| MGDG((34:4) | 0.000 | 0.000 | 0.000 | 0.000 | 0.000 | 0.000 | 0.000 | 0.000 | 0.000 | 0.000 | 0.000 | 0.000 | 0.000 | 0.000 |
| MGDG((34:3) | 0.000 | 0.000 | 0.000 | 0.000 | 0.000 | 0.000 | 0.000 | 0.000 | 0.000 | 0.000 | 0.000 | 0.000 | 0.000 | 0.000 |
| MGDG((34:2) | 0.000 | 0.000 | 0.000 | 0.000 | 0.000 | 0.000 | 0.000 | 0.000 | 0.000 | 0.000 | 0.000 | 0.000 | 0.000 | 0.000 |
| MGDG((34:1) | 0.000 | 0.000 | 0.000 | 0.000 | 0.000 | 0.000 | 0.000 | 0.000 | 0.000 | 0.000 | 0.000 | 0.000 | 0.000 | 0.000 |
| MGDG((36:6) | 0.001 | 0.000 | 0.000 | 0.001 | 0.002 | 0.001 | 0.001 | 0.000 | 0.001 | 0.000 | 0.000 | 0.000 | 0.000 | 0.000 |
| MGDG((36:5) | 0.001 | 0.000 | 0.001 | 0.000 | 0.004 | 0.004 | 0.000 | 0.000 | 0.000 | 0.000 | 0.001 | 0.001 | 0.000 | 0.000 |
| MGDG((36:4) | 0.002 | 0.000 | 0.000 | 0.000 | 0.003 | 0.003 | 0.000 | 0.000 | 0.000 | 0.000 | 0.001 | 0.001 | 0.000 | 0.000 |
| MGDG((36:3) | 0.000 | 0.000 | 0.000 | 0.000 | 0.000 | 0.000 | 0.000 | 0.000 | 0.000 | 0.000 | 0.000 | 0.000 | 0.000 | 0.000 |
| MGDG((36:2) | 0.000 | 0.000 | 0.000 | 0.000 | 0.000 | 0.000 | 0.000 | 0.000 | 0.000 | 0.000 | 0.000 | 0.000 | 0.000 | 0.000 |
| MGDG((38:6) | 0.000 | 0.000 | 0.000 | 0.000 | 0.000 | 0.000 | 0.000 | 0.000 | 0.000 | 0.000 | 0.000 | 0.000 | 0.000 | 0.000 |
| MGDG((38:5) | 0.000 | 0.000 | 0.000 | 0.000 | 0.000 | 0.000 | 0.000 | 0.000 | 0.000 | 0.000 | 0.000 | 0.000 | 0.000 | 0.000 |
| MGDG((38:4) | 0.000 | 0.000 | 0.000 | 0.000 | 0.000 | 0.000 | 0.000 | 0.000 | 0.000 | 0.000 | 0.000 | 0.000 | 0.000 | 0.000 |
| MGDG((38:3) | 0.000 | 0.000 | 0.000 | 0.000 | 0.000 | 0.000 | 0.000 | 0.000 | 0.000 | 0.000 | 0.000 | 0.000 | 0.000 | 0.000 |
| **Total MGDG** | **0.004** | **0.002** | **0.004** | **0.004** | **0.010** | **0.009** | **0.001** | **0.000** | **0.002** | **0.001** | **0.002** | **0.001** | **0.001** | **0.000** |

| Compound* | *Laurus nobilis* mean | std | *Liriodendron tulipifera* mean | std | *Bambusa oldhamii* mean | std | *Triadica sebifera* mean | std | *Geijera parviflora* mean | std | *Distictis buccinatoria* mean | std | *Encelia farinosa* mean | std |
|---|---|---|---|---|---|---|---|---|---|---|---|---|---|---|
| PG((32:0) | 0.000 | 0.000 | 0.000 | 0.000 | 0.000 | 0.000 | 0.000 | 0.000 | 0.000 | 0.000 | 0.000 | 0.000 | 0.000 | 0.000 |
| PG((34:4) | 0.000 | 0.000 | 0.000 | 0.000 | 0.000 | 0.000 | 0.000 | 0.000 | 0.000 | 0.000 | 0.000 | 0.000 | 0.000 | 0.000 |
| PG((34:3) | 0.000 | 0.000 | 0.000 | 0.000 | 0.000 | 0.000 | 0.000 | 0.000 | 0.000 | 0.000 | 0.000 | 0.000 | 0.000 | 0.000 |
| PG((34:2) | 0.000 | 0.000 | 0.000 | 0.000 | 0.000 | 0.000 | 0.000 | 0.000 | 0.000 | 0.000 | 0.000 | 0.000 | 0.000 | 0.000 |
| PG((34:1) | 0.000 | 0.000 | 0.000 | 0.000 | 0.000 | 0.000 | 0.000 | 0.000 | 0.000 | 0.000 | 0.000 | 0.000 | 0.000 | 0.000 |
| **Total PG** | **0.000** | **0.000** | **0.000** | **0.000** | **0.000** | **0.001** | **0.000** | **0.000** | **0.000** | **0.000** | **0.000** | **0.000** | **0.000** | **0.000** |
| LPG((16:1) | 0.000 | 0.000 | 0.000 | 0.000 | 0.000 | 0.000 | 0.000 | 0.000 | 0.001 | 0.000 | 0.000 | 0.000 | 0.000 | 0.000 |
| LPG((16:0) | 0.005 | 0.001 | 0.003 | 0.003 | 0.008 | 0.001 | 0.002 | 0.001 | 0.003 | 0.001 | 0.003 | 0.001 | 0.002 | 0.000 |
| LPG((18:3) | 0.000 | 0.000 | 0.000 | 0.000 | 0.000 | 0.000 | 0.000 | 0.000 | 0.000 | 0.000 | 0.000 | 0.000 | 0.000 | 0.000 |
| LPG((18:2) | 0.001 | 0.000 | 0.000 | 0.000 | 0.000 | 0.000 | 0.000 | 0.000 | 0.000 | 0.000 | 0.000 | 0.000 | 0.000 | 0.000 |
| LPG((18:1) | 0.001 | 0.001 | 0.001 | 0.000 | 0.000 | 0.000 | 0.000 | 0.000 | 0.000 | 0.000 | 0.000 | 0.000 | 0.000 | 0.000 |
| **Total LysoPG** | **0.006** | **0.002** | **0.004** | **0.002** | **0.008** | **0.001** | **0.002** | **0.001** | **0.004** | **0.001** | **0.003** | **0.002** | **0.002** | **0.000** |
| LPC((16:0) | 0.000 | 0.000 | 0.000 | 0.000 | 0.000 | 0.000 | 0.000 | 0.000 | 0.000 | 0.000 | 0.000 | 0.000 | 0.000 | 0.000 |
| LPC((18:3) | 0.000 | 0.000 | 0.000 | 0.000 | 0.000 | 0.000 | 0.000 | 0.000 | 0.000 | 0.000 | 0.000 | 0.000 | 0.000 | 0.000 |
| LPC((18:2) | 0.000 | 0.000 | 0.000 | 0.000 | 0.000 | 0.000 | 0.000 | 0.000 | 0.000 | 0.000 | 0.000 | 0.000 | 0.000 | 0.000 |
| LPC((18:1) | 0.000 | 0.000 | 0.000 | 0.000 | 0.000 | 0.000 | 0.000 | 0.000 | 0.000 | 0.000 | 0.000 | 0.000 | 0.000 | 0.000 |
| **Total LysoPC** | **0.000** | **0.000** | **0.000** | **0.000** | **0.001** | **0.000** | **0.000** | **0.000** | **0.000** | **0.000** | **0.000** | **0.000** | **0.000** | **0.000** |
| LPE((16:0) | 0.002 | 0.001 | 0.000 | 0.000 | 0.001 | 0.001 | 0.000 | 0.000 | 0.000 | 0.000 | 0.000 | 0.000 | 0.000 | 0.000 |
| LPE((18:3) | 0.000 | 0.000 | 0.000 | 0.000 | 0.000 | 0.000 | 0.000 | 0.000 | 0.000 | 0.000 | 0.000 | 0.000 | 0.000 | 0.000 |
| LPE((18:2) | 0.000 | 0.000 | 0.000 | 0.000 | 0.000 | 0.000 | 0.000 | 0.000 | 0.000 | 0.000 | 0.000 | 0.000 | 0.000 | 0.000 |
| LPE((18:1) | 0.000 | 0.001 | 0.000 | 0.000 | 0.000 | 0.000 | 0.000 | 0.000 | 0.000 | 0.000 | 0.000 | 0.000 | 0.000 | 0.000 |
| **Total LysoPE** | **0.003** | **0.002** | **0.000** | **0.000** | **0.001** | **0.001** | **0.000** | **0.000** | **0.000** | **0.000** | **0.000** | **0.000** | **0.000** | **0.000** |
| PC((32:0) | 0.000 | 0.000 | 0.000 | 0.000 | 0.000 | 0.000 | 0.000 | 0.000 | 0.000 | 0.000 | 0.000 | 0.000 | 0.000 | 0.000 |
| PC((34:4) | 0.000 | 0.000 | 0.000 | 0.000 | 0.000 | 0.000 | 0.000 | 0.000 | 0.000 | 0.000 | 0.000 | 0.000 | 0.000 | 0.000 |
| PC((34:3) | 0.000 | 0.000 | 0.000 | 0.000 | 0.001 | 0.002 | 0.001 | 0.001 | 0.001 | 0.000 | 0.000 | 0.000 | 0.001 | 0.000 |
| PC((34:2) | 0.001 | 0.001 | 0.001 | 0.000 | 0.004 | 0.006 | 0.001 | 0.000 | 0.002 | 0.000 | 0.000 | 0.000 | 0.004 | 0.000 |
| PC((34:1) | 0.000 | 0.000 | 0.000 | 0.000 | 0.001 | 0.001 | 0.002 | 0.001 | 0.001 | 0.000 | 0.001 | 0.000 | 0.001 | 0.000 |
| PC((36:6) | 0.000 | 0.000 | 0.000 | 0.000 | 0.000 | 0.000 | 0.000 | 0.000 | 0.000 | 0.000 | 0.000 | 0.000 | 0.000 | 0.000 |
| PC((36:5) | 0.000 | 0.000 | 0.000 | 0.000 | 0.001 | 0.001 | 0.000 | 0.000 | 0.000 | 0.000 | 0.000 | 0.000 | 0.001 | 0.000 |
| PC((36:4) | 0.000 | 0.001 | 0.001 | 0.000 | 0.001 | 0.002 | 0.002 | 0.001 | 0.001 | 0.001 | 0.001 | 0.000 | 0.003 | 0.002 |
| PC((36:3) | 0.000 | 0.000 | 0.001 | 0.000 | 0.001 | 0.001 | 0.001 | 0.001 | 0.001 | 0.000 | 0.001 | 0.000 | 0.001 | 0.000 |

| | *Laurus nobilis* | | *Liriodendron tulipifera* | | *Bambusa oldhamii* | | *Triadica sebifera* | | *Geijera parviflora* | | *Distictis buccinatoria* | | *Encelia farinosa* | |
|---|---|---|---|---|---|---|---|---|---|---|---|---|---|---|
| **Compound*** | **mean** | **std** | **mean** | **std** | **mean** | **std** | **mean** | **std** | **mean** | **std** | **mean** | **std** | **mean** | **std** |
| PC((36:2) | 0.000 | 0.000 | 0.000 | 0.000 | 0.000 | 0.000 | 0.001 | 0.000 | 0.000 | 0.000 | 0.000 | 0.000 | 0.001 | 0.000 |
| PC((36:1) | 0.000 | 0.000 | 0.000 | 0.000 | 0.000 | 0.000 | 0.000 | 0.000 | 0.000 | 0.000 | 0.000 | 0.000 | 0.000 | 0.000 |
| PC((38:6) | 0.000 | 0.000 | 0.000 | 0.000 | 0.000 | 0.000 | 0.000 | 0.000 | 0.000 | 0.000 | 0.000 | 0.000 | 0.000 | 0.000 |
| PC((38:5) | 0.000 | 0.000 | 0.000 | 0.000 | 0.000 | 0.000 | 0.000 | 0.000 | 0.000 | 0.000 | 0.000 | 0.000 | 0.000 | 0.000 |
| PC((38:4) | 0.000 | 0.000 | 0.000 | 0.000 | 0.000 | 0.000 | 0.000 | 0.000 | 0.000 | 0.000 | 0.000 | 0.000 | 0.000 | 0.000 |
| PC((38:3) | 0.000 | 0.000 | 0.000 | 0.000 | 0.000 | 0.000 | 0.000 | 0.000 | 0.000 | 0.000 | 0.000 | 0.000 | 0.000 | 0.000 |
| PC((38:2) | 0.000 | 0.000 | 0.000 | 0.000 | 0.000 | 0.000 | 0.000 | 0.000 | 0.000 | 0.000 | 0.000 | 0.000 | 0.000 | 0.000 |
| **Total PC** | **0.002** | **0.002** | **0.003** | **0.001** | **0.008** | **0.013** | **0.008** | **0.005** | **0.007** | **0.001** | **0.003** | **0.000** | **0.012** | **0.004** |
| PE((32:1) | 0.000 | 0.000 | 0.000 | 0.000 | 0.000 | 0.000 | 0.000 | 0.000 | 0.000 | 0.000 | 0.000 | 0.000 | 0.000 | 0.000 |
| PE((32:0) | 0.000 | 0.000 | 0.000 | 0.000 | 0.000 | 0.000 | 0.000 | 0.000 | 0.000 | 0.000 | 0.000 | 0.000 | 0.000 | 0.000 |
| PE((34:4) | 0.000 | 0.000 | 0.000 | 0.000 | 0.000 | 0.000 | 0.000 | 0.000 | 0.000 | 0.000 | 0.000 | 0.000 | 0.000 | 0.000 |
| PE((34:3) | 0.000 | 0.000 | 0.000 | 0.000 | 0.000 | 0.000 | 0.000 | 0.000 | 0.000 | 0.000 | 0.000 | 0.000 | 0.000 | 0.000 |
| PE((34:2) | 0.000 | 0.000 | 0.000 | 0.000 | 0.000 | 0.000 | 0.000 | 0.001 | 0.000 | 0.000 | 0.000 | 0.000 | 0.000 | 0.000 |
| PE((34:1) | 0.000 | 0.000 | 0.000 | 0.000 | 0.000 | 0.000 | 0.000 | 0.000 | 0.000 | 0.000 | 0.000 | 0.000 | 0.000 | 0.000 |
| PE((36:6) | 0.000 | 0.000 | 0.000 | 0.000 | 0.000 | 0.000 | 0.000 | 0.000 | 0.000 | 0.000 | 0.000 | 0.000 | 0.000 | 0.000 |
| PE((36:5) | 0.000 | 0.000 | 0.000 | 0.000 | 0.000 | 0.000 | 0.000 | 0.000 | 0.000 | 0.000 | 0.000 | 0.000 | 0.000 | 0.000 |
| PE((36:4) | 0.000 | 0.000 | 0.000 | 0.000 | 0.000 | 0.000 | 0.000 | 0.000 | 0.000 | 0.000 | 0.000 | 0.000 | 0.000 | 0.000 |
| PE((36:3) | 0.000 | 0.000 | 0.000 | 0.000 | 0.000 | 0.000 | 0.000 | 0.000 | 0.000 | 0.000 | 0.000 | 0.000 | 0.000 | 0.000 |
| PE((36:2) | 0.000 | 0.000 | 0.000 | 0.000 | 0.000 | 0.000 | 0.000 | 0.000 | 0.000 | 0.000 | 0.000 | 0.000 | 0.000 | 0.000 |
| PE((36:1) | 0.000 | 0.000 | 0.000 | 0.000 | 0.000 | 0.000 | 0.000 | 0.000 | 0.000 | 0.000 | 0.000 | 0.000 | 0.000 | 0.000 |
| PE((42:2) | 0.000 | 0.000 | 0.000 | 0.000 | 0.000 | 0.000 | 0.000 | 0.000 | 0.000 | 0.000 | 0.000 | 0.000 | 0.000 | 0.000 |
| **Total PE** | **0.000** | **0.000** | **0.000** | **0.000** | **0.000** | **0.000** | **0.002** | **0.002** | **0.000** | **0.000** | **0.000** | **0.000** | **0.000** | **0.000** |
| PI((32:2) | 0.000 | 0.000 | 0.000 | 0.000 | 0.000 | 0.000 | 0.000 | 0.000 | 0.000 | 0.000 | 0.000 | 0.000 | 0.000 | 0.000 |
| PI((32:1) | 0.000 | 0.000 | 0.000 | 0.000 | 0.000 | 0.000 | 0.000 | 0.000 | 0.000 | 0.000 | 0.000 | 0.000 | 0.000 | 0.000 |
| PI((32:0) | 0.000 | 0.000 | 0.000 | 0.000 | 0.000 | 0.000 | 0.000 | 0.000 | 0.000 | 0.000 | 0.000 | 0.000 | 0.000 | 0.000 |
| PI((34:4) | 0.000 | 0.000 | 0.000 | 0.000 | 0.000 | 0.000 | 0.000 | 0.000 | 0.000 | 0.000 | 0.000 | 0.000 | 0.000 | 0.000 |
| PI((34:3) | 0.000 | 0.000 | 0.000 | 0.000 | 0.000 | 0.000 | 0.000 | 0.001 | 0.000 | 0.000 | 0.000 | 0.000 | 0.000 | 0.000 |
| PI((34:2) | 0.000 | 0.001 | 0.000 | 0.000 | 0.000 | 0.001 | 0.001 | 0.001 | 0.000 | 0.000 | 0.000 | 0.000 | 0.000 | 0.000 |
| PI((34:1) | 0.000 | 0.000 | 0.000 | 0.000 | 0.000 | 0.000 | 0.000 | 0.001 | 0.000 | 0.000 | 0.000 | 0.000 | 0.000 | 0.000 |
| PI((36:6) | 0.000 | 0.000 | 0.000 | 0.000 | 0.000 | 0.000 | 0.000 | 0.000 | 0.000 | 0.000 | 0.000 | 0.000 | 0.000 | 0.000 |
| PI((36:5) | 0.000 | 0.000 | 0.000 | 0.000 | 0.000 | 0.000 | 0.000 | 0.000 | 0.000 | 0.000 | 0.000 | 0.000 | 0.000 | 0.000 |
| PI((36:4) | 0.000 | 0.000 | 0.000 | 0.000 | 0.000 | 0.000 | 0.000 | 0.000 | 0.000 | 0.000 | 0.000 | 0.000 | 0.000 | 0.000 |

| Compound* | *Laurus nobilis* mean | std | *Liriodendron tulipifera* mean | std | *Bambusa oldhamii* mean | std | *Triadica sebifera* mean | std | *Geijera parviflora* mean | std | *Distictis buccinatoria* mean | std | *Encelia farinosa* mean | std |
|---|---|---|---|---|---|---|---|---|---|---|---|---|---|---|
| PI((36:3) | 0.000 | 0.000 | 0.000 | 0.000 | 0.000 | 0.000 | 0.000 | 0.000 | 0.000 | 0.000 | 0.000 | 0.000 | 0.000 | 0.000 |
| PI((36:2) | 0.000 | 0.000 | 0.000 | 0.000 | 0.000 | 0.000 | 0.000 | 0.000 | 0.000 | 0.000 | 0.000 | 0.000 | 0.000 | 0.000 |
| PI((36:1) | 0.000 | 0.000 | 0.000 | 0.000 | 0.000 | 0.000 | 0.000 | 0.000 | 0.000 | 0.000 | 0.000 | 0.000 | 0.000 | 0.000 |
| **Total PI** | **0.001** | **0.001** | **0.000** | **0.000** | **0.001** | **0.001** | **0.002** | **0.003** | **0.000** | **0.000** | **0.000** | **0.000** | **0.000** | **0.000** |
| PS((34:3) | 0.000 | 0.000 | 0.000 | 0.000 | 0.000 | 0.000 | 0.000 | 0.000 | 0.000 | 0.000 | 0.000 | 0.000 | 0.000 | 0.000 |
| PS((34:2) | 0.000 | 0.000 | 0.000 | 0.000 | 0.000 | 0.000 | 0.000 | 0.000 | 0.000 | 0.000 | 0.000 | 0.000 | 0.000 | 0.000 |
| PS((34:1) | 0.000 | 0.000 | 0.000 | 0.000 | 0.000 | 0.000 | 0.000 | 0.000 | 0.000 | 0.000 | 0.000 | 0.000 | 0.000 | 0.000 |
| PS((36:6) | 0.000 | 0.000 | 0.000 | 0.000 | 0.000 | 0.000 | 0.000 | 0.000 | 0.000 | 0.000 | 0.000 | 0.000 | 0.000 | 0.000 |
| PS((36:5) | 0.000 | 0.000 | 0.000 | 0.000 | 0.000 | 0.000 | 0.000 | 0.000 | 0.000 | 0.000 | 0.000 | 0.000 | 0.000 | 0.000 |
| PS((36:4) | 0.000 | 0.000 | 0.000 | 0.000 | 0.000 | 0.000 | 0.000 | 0.000 | 0.000 | 0.000 | 0.000 | 0.000 | 0.000 | 0.000 |
| PS((36:3) | 0.000 | 0.000 | 0.000 | 0.000 | 0.000 | 0.000 | 0.000 | 0.000 | 0.000 | 0.000 | 0.000 | 0.000 | 0.000 | 0.000 |
| PS((36:2) | 0.000 | 0.000 | 0.000 | 0.000 | 0.000 | 0.000 | 0.000 | 0.000 | 0.000 | 0.000 | 0.000 | 0.000 | 0.000 | 0.000 |
| PS((38:3) | 0.000 | 0.000 | 0.000 | 0.000 | 0.000 | 0.000 | 0.000 | 0.000 | 0.000 | 0.000 | 0.000 | 0.000 | 0.000 | 0.000 |
| PS((38:2) | 0.000 | 0.000 | 0.000 | 0.000 | 0.000 | 0.000 | 0.000 | 0.000 | 0.000 | 0.000 | 0.000 | 0.000 | 0.000 | 0.000 |
| PS((38:1) | 0.000 | 0.001 | 0.000 | 0.000 | 0.000 | 0.000 | 0.000 | 0.000 | 0.000 | 0.000 | 0.000 | 0.000 | 0.000 | 0.000 |
| PS((40:3) | 0.000 | 0.000 | 0.000 | 0.000 | 0.000 | 0.000 | 0.000 | 0.000 | 0.000 | 0.000 | 0.000 | 0.000 | 0.000 | 0.000 |
| PS((40:2) | 0.000 | 0.000 | 0.000 | 0.000 | 0.000 | 0.000 | 0.000 | 0.000 | 0.000 | 0.000 | 0.000 | 0.000 | 0.000 | 0.000 |
| PS((40:1) | 0.000 | 0.000 | 0.000 | 0.000 | 0.000 | 0.000 | 0.000 | 0.000 | 0.000 | 0.000 | 0.000 | 0.000 | 0.000 | 0.000 |
| PS((42:4) | 0.000 | 0.000 | 0.000 | 0.000 | 0.000 | 0.000 | 0.000 | 0.000 | 0.000 | 0.000 | 0.000 | 0.000 | 0.000 | 0.000 |
| PS((42:3) | 0.000 | 0.000 | 0.000 | 0.000 | 0.000 | 0.000 | 0.000 | 0.000 | 0.000 | 0.000 | 0.000 | 0.000 | 0.000 | 0.000 |
| PS((42:2) | 0.000 | 0.000 | 0.000 | 0.000 | 0.000 | 0.000 | 0.000 | 0.000 | 0.000 | 0.000 | 0.000 | 0.000 | 0.000 | 0.000 |
| PS((44:3) | 0.000 | 0.000 | 0.000 | 0.000 | 0.000 | 0.000 | 0.000 | 0.000 | 0.000 | 0.000 | 0.000 | 0.000 | 0.000 | 0.000 |
| **Total PS** | **0.001** | **0.001** | **0.000** | **0.000** | **0.000** | **0.000** | **0.000** | **0.000** | **0.000** | **0.000** | **0.000** | **0.001** | **0.000** | **0.000** |
| PA((32:0) | 0.000 | 0.000 | 0.000 | 0.000 | 0.000 | 0.000 | 0.000 | 0.000 | 0.000 | 0.000 | 0.000 | 0.000 | 0.000 | 0.000 |
| PA((34:4) | 0.000 | 0.000 | 0.000 | 0.000 | 0.000 | 0.000 | 0.000 | 0.000 | 0.000 | 0.000 | 0.000 | 0.000 | 0.000 | 0.000 |
| PA((34:3) | 0.000 | 0.000 | 0.000 | 0.000 | 0.000 | 0.000 | 0.002 | 0.001 | 0.002 | 0.001 | 0.000 | 0.000 | 0.001 | 0.000 |
| PA((34:2) | 0.000 | 0.000 | 0.002 | 0.001 | 0.001 | 0.001 | 0.002 | 0.001 | 0.003 | 0.002 | 0.000 | 0.000 | 0.003 | 0.001 |
| PA((34:1) | 0.000 | 0.000 | 0.001 | 0.001 | 0.000 | 0.000 | 0.002 | 0.002 | 0.001 | 0.001 | 0.000 | 0.000 | 0.001 | 0.000 |
| PA((36:6) | 0.000 | 0.000 | 0.000 | 0.000 | 0.000 | 0.000 | 0.000 | 0.000 | 0.000 | 0.000 | 0.000 | 0.000 | 0.000 | 0.000 |
| PA((36:5) | 0.000 | 0.000 | 0.000 | 0.000 | 0.000 | 0.000 | 0.000 | 0.000 | 0.001 | 0.001 | 0.000 | 0.000 | 0.001 | 0.000 |
| PA((36:4) | 0.000 | 0.000 | 0.002 | 0.001 | 0.000 | 0.000 | 0.002 | 0.001 | 0.001 | 0.001 | 0.000 | 0.000 | 0.001 | 0.000 |
| PA((36:3) | 0.000 | 0.000 | 0.001 | 0.000 | 0.000 | 0.000 | 0.001 | 0.001 | 0.001 | 0.001 | 0.000 | 0.000 | 0.001 | 0.000 |

| Compound* | *Laurus nobilis* mean | std | *Liriodendron tulipifera* mean | std | *Bambusa oldhamii* mean | std | *Triadica sebifera* mean | std | *Geijera parviflora* mean | std | *Distictis buccinatoria* mean | std | *Encelia farinosa* mean | std |
|---|---|---|---|---|---|---|---|---|---|---|---|---|---|---|
| PA((36:2) | 0.000 | 0.000 | 0.001 | 0.000 | 0.000 | 0.000 | 0.001 | 0.001 | 0.000 | 0.000 | 0.000 | 0.000 | 0.000 | 0.000 |
| **Total PA** | **0.000** | **0.000** | **0.007** | **0.004** | **0.002** | **0.002** | **0.011** | **0.006** | **0.009** | **0.006** | **0.002** | **0.000** | **0.006** | **0.001** |
| SQDG((34:3) | | | 0.000 | 0.000 | | | 0.000 | 0.000 | 0.000 | 0.000 | 0.000 | 0.000 | 0.000 | 0.000 |
| SQDG((34:2) | | | 0.000 | 0.000 | | | 0.000 | 0.000 | 0.000 | 0.000 | 0.000 | 0.000 | 0.000 | 0.000 |
| SQDG((34:1) | | | 0.000 | 0.000 | | | 0.000 | 0.000 | 0.000 | 0.000 | 0.000 | 0.000 | 0.000 | 0.000 |
| SQDG((36:5) | | | 0.000 | 0.000 | | | 0.000 | 0.000 | 0.000 | 0.000 | 0.000 | 0.000 | 0.000 | 0.000 |
| SQDG((36:4) | | | 0.000 | 0.000 | | | 0.000 | 0.000 | 0.000 | 0.000 | 0.000 | 0.000 | 0.000 | 0.000 |
| **Total SQDG** | | | **0.001** | **0.001** | | | **0.003** | **0.000** | **0.003** | **0.001** | **0.003** | **0.001** | **0.009** | **0.003** |

* DGDG = digalactosyldiacylglycerol; LPC = lysophosphatidylcholine; LPE = lysophosphatidylethanolamine; LPG = lysophosphatidylglycerol; MGDG = monogalactosyldiacylglycerol; PA = phosphatidic acid; PC = phosphatidylcholine; PE = phosphatidylethanolamine; PG = phosphatidylglycerol; PI = phosphatidylinositol; PS = phosphatidylserine
